# Impacts of genome architecture on the repeatability of polygenic adaptation

**DOI:** 10.64898/2026.03.26.714499

**Authors:** Zhenyong Du, Johannes Wirtz, Qilin Li, Alexander Taylor, Lydia Larsen, Samuel Lu, David B. Stern, Carol Eunmi Lee

## Abstract

A central but poorly resolved question regards how genome architecture shapes the selection response and repeatability of polygenic adaptation. We addressed this question using Evolve-and-Resequence experiments under rapid salinity decline in two sibling species (clades) of the invasive copepod *Eurytemora affinis* complex that differ strikingly in chromosome number (15 versus 4). The 4-chromosome genome arose from chromosomal fusions of the ancestral 15-chromosome genome, bringing together coadapted alleles at fusion sites. Across 10–20 generations of selection, both clades adapted to low salinity but followed divergent evolutionary trajectories. The selection lines of the 15-chromosome clade exhibited highly parallel genomic responses, whereas the 4-chromosome clade lines showed delayed and less repeatable responses. Forward genetic simulations revealed that strong synergistic epistasis among beneficial alleles best explained elevated parallelism in the 15-chromosome clade. Additional simulations varying chromosome number and epistasis revealed that strong positive epistasis, combined with high chromosome numbers, enhances parallelism by enabling recombination to assemble coadapted allelic combinations. In contrast, the high starting frequency of beneficial alleles in the 4-chromosome clade lines indicated selection on standing variation, likely acting on alternative linked haplotypes. These findings demonstrate that genome architecture and gene–gene interactions jointly determine the dynamics and predictability of polygenic adaptation.

## Introduction

A fundamental question in evolutionary biology regards how selection acts on complex traits and the mechanisms and repeatability of polygenic adaptation^1–9^. When faced with sudden environmental change, populations can adapt surprisingly quickly, even when selection acts on polygenic traits encoded by many genes^4,10–15^. Polygenic adaptation is crucially important in response to environmental change, as selection in response to abiotic variables often acts on physiological traits, which are typically polygenic in nature. Despite its importance, until recently mechanisms of polygenic adaptation had remained poorly understood, as dissecting polygenic responses requires information on genome architecture and genome-wide selection responses of populations.

Independent populations, or experimental replicates, may involve similar genetic solutions that reflect parallel evolutionary responses^4,13^. Alternatively, populations may experience divergent genetic paths that nevertheless produce comparable adaptive outcomes at the phenotypic level^3,4,9,16–18^. Dissecting the specific mechanisms of parallel evolution of complex traits is crucial for understanding how natural selection acts on populations and which factors constrain or promote repeatability^3,17^. In addition, determining the repeatability of selection responses is crucial for understanding the speed and extent to which populations could evolve in response to rapid environmental stress and avoid extinctions in the face of rapid global change^19–21^.

The roles of genome architecture underlying polygenic traits in impacting selection responses and adaptive repeatability have remained poorly explored. While the number of loci contributing to a trait is an important factor influencing constraints on natural selection (e.g., pleiotropy) and the extent of parallelism among replicate evolutionary events^5,8,17,22^, the localization and arrangement of these loci within the genome are also likely to be critically important^23,24^. In particular, genomic features such as the positions and arrangement of genes, chromosome number, and recombination rate influence gene–gene interactions and determine the extent to which loci are co-inherited and co-regulated^25–31^.

Among genomic features, the number of chromosomes, for a given genome size, could have profound impacts on how selection acts on coadapted gene complexes. On the one hand, genomes with fewer chromosomes and lower recombination rates may preserve combinations of coadapted alleles, making independent populations more likely to reuse the same multi-locus linked haplotypes, thereby increasing parallelism. However, low recombination can also lead to the maintenance of distinct beneficial haplotypes within a population, enabling different replicate lines or populations to fix alternative haplotypes and thereby reducing parallelism^32,33^. In addition, linked selection would likely reduce parallelism in genomes with longer chromosomes, as beneficial allelic combinations would be linked to long stretches of neutral and potentially segregating loci, leading to selection favoring different allelic combinations^34–37^. By contrast, genomes with many chromosomes, and consequently higher recombination rates, would enable greater combinations of beneficial alleles at different loci, allowing natural selection to more effectively assemble the most favorable allelic combinations within a genome^23,38,39^. If positive epistasis favors specific sets of alleles, recombination can consistently generate the same adaptive combinations across replicates, strongly promoting parallel evolution^4,40,41^. Thus, the impact of genome architecture on the repeatability of adaptation could depend in part on the interplay between linkage among beneficial alleles and the recombination required to assemble them^11,25–27^.

To explore how genome architecture impacts the selection response and repeatability of polygenic adaptation, we examined the selection responses of sibling species with differing chromosome numbers in the copepod *Eurytemora affinis* species complex^29,42^. Populations from this species complex are native to saline habitats, but have repeatedly invaded freshwater habitats within the past 80 years, likely due to shipping and ballast water transport^43^ (Figure 1a). This species complex is composed of sibling species (clades) that differ drastically in haploid chromosome number (4–15 chromosomes; Figure 1b) but possess similar gene content and genome size^29,42^. Despite the large differences in genome architecture, many of the same loci show signatures of natural selection during salinity transitions across the different clades^4,13,14,44^. Most notably, our prior study revealed that the 4-chromosome genome (Atlantic clade of the *E. affinis* complex, *E. carolleeae*) arose from multiple chromosomal fusions of the ancestral 15-chromosome genome (Europe clade, *E. affinis* proper), bringing together coadapted alleles at the fusion sites (Figure 1c)^29^. As such, this contrast of pre- and post-fusion genomes provides an unprecedented natural model to test how genome architecture impacts selection response in populations with these genomes.

**Figure 1.**
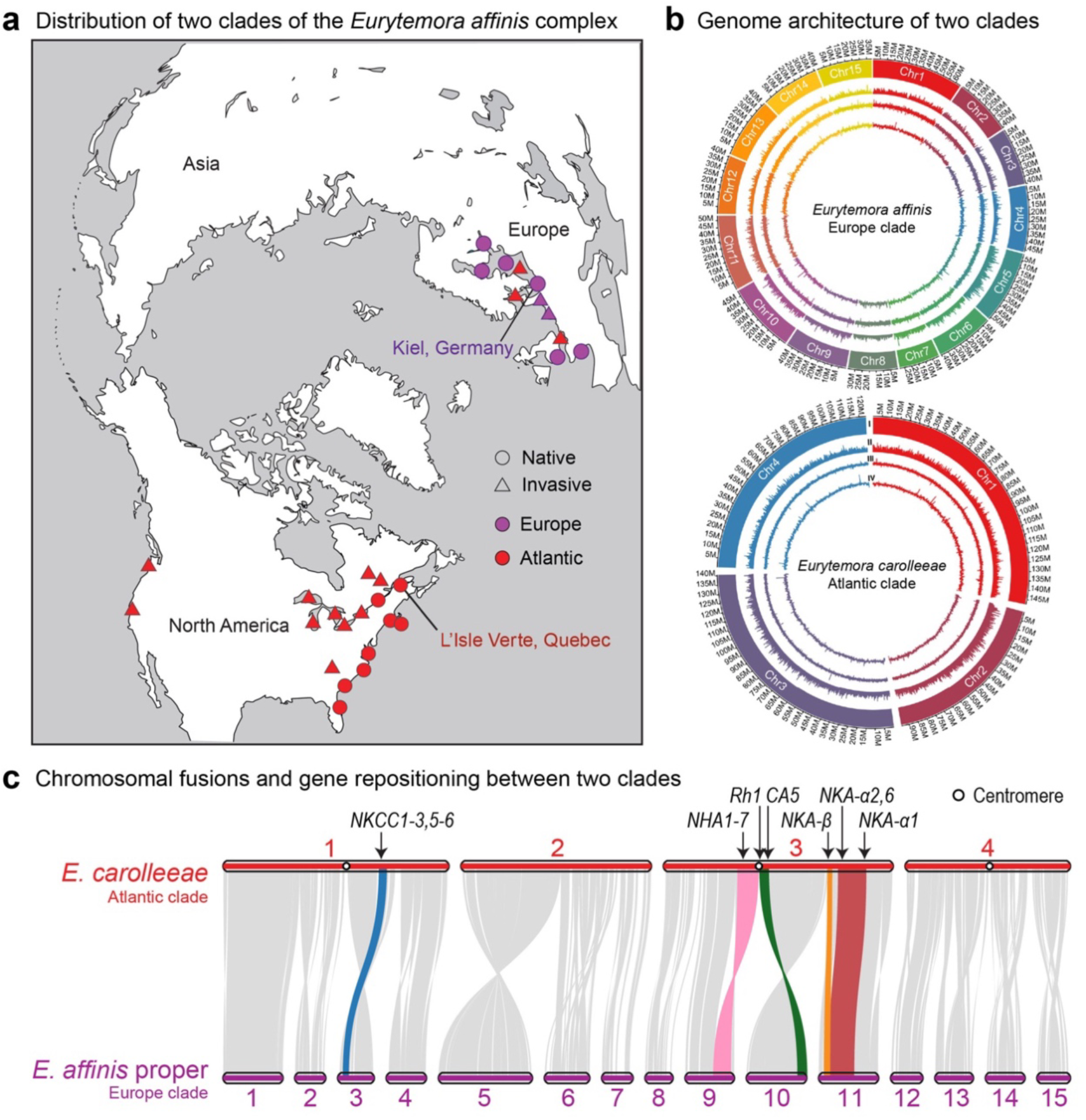
Geographic distribution, genome architecture, and chromosomal fusions in two clades of the *Eurytemora affinis* species complex. **(a)** Geographic distribution of two clades (sibling species) of the *E. affinis* species complex^43,45^, showing the Europe clade (*E. affinis* proper; haploid chromosome number = 15, purple) and the Atlantic clade (*E. carolleeae*; haploid chromosome number = 4, red). Native (circles) and invaded (triangles) locations are shown for populations from each clade. **(b)** Genome architecture of the two clades, illustrating the dramatic difference in chromosome number (15 versus 4) due to chromosomal fusions in the Atlantic clade^29^. **(c)** Schematic representation of chromosomal fusions and associated gene repositioning between the two clades. Fifteen ancestral chromosomes in the Europe clade correspond to four fused chromosomes in the Atlantic clade, with several key ion transporter genes (e.g., *NKCC*, *NHA*, and *NKA* gene families) repositioned near fusion sites^29^. Centromere positions are indicated by open circles.

Our prior results suggest that different mechanisms of adaptation might be operating depending on chromosome number. For the Europe clade, with a high number of chromosomes (15), in our prior Evolve-and-Resequence (E&R) study, we had found strikingly high levels of parallelism among selection lines adapting to salinity decline in the laboratory^4^. Based on theoretical simulations, this unexpected high level of parallelism was found to be consistent with the action of positive synergistic epistasis among adaptive alleles, bringing coadapted alleles together within genomes^4^. We also observed an exceptionally high degree of parallel evolution during saline-to-freshwater invasions by wild populations from the Atlantic (4 chromosomes) and Gulf (7 chromosomes) clades, with selection often favoring the same SNPs during independent invasions^13^. In this prior study, we found that a large proportion of SNPs under directional selection during invasions were at intermediate frequency in the native populations, suggesting the maintenance of genetic variation in the native range through balancing selection^13^. This balanced polymorphism in the native range could then be favored by natural selection during invasions and could help promote parallel evolution^12,32^.

These prior results above suggest that positive synergistic epistasis and selection on standing genetic variation could both play important roles in driving parallel evolution^1,4,13,46^. However, we do not fully understand the conditions under which one mechanism might prevail over the other. Might genome architecture, particularly chromosome number, impact the selection response of a population? And might the selection regime of a population influence the particular genome architecture that might be favorable in facilitating adaptation to environmental change?

Thus, to address these questions, we determined the impacts of genome architecture, particularly chromosome number, on the repeatability of selection response using Evolve-and-Resequence (E&R) experiments^5,47^ under rapid salinity decline in two clades that differ in chromosome number (15 versus 4). We also performed simulations to explore the selective mechanisms that might be involved in driving parallel evolution. Our specific goals were to (1) determine how genome architecture affects the trajectories of beneficial alleles and the number and effect sizes of alleles contributing to adaptation, (2) examine the impacts of genome architecture on the extent of parallelism among replicate selection lines in the E&R experiments, and (3) explore the selective mechanisms underlying the parallel responses, such as the potential roles of positive epistasis, linkage of beneficial alleles, and selection on standing genetic variation.

To achieve these goals, we imposed laboratory selection on replicate selection lines from the two clades with different karyotypes (15 versus 4 chromosomes; Figure 1b) under declining salinity for 10–20 generations (Figure 2a, b). We then performed time-resolved whole-genome pooled sequencing (E&R) to monitor beneficial allele frequency changes in the replicate selection lines across generations (Figure 2a, b). In addition, we employed forward genetic simulations tailored to the genomic parameters of each clade (effective population size, number of selected loci, and chromosome number) to evaluate which evolutionary processes best explained the observed extent of parallelism. In particular, we examined how chromosome number influenced the relative importance of positive synergistic epistasis versus linkage among beneficial alleles in shaping parallel adaptation.

**Figure 2.**
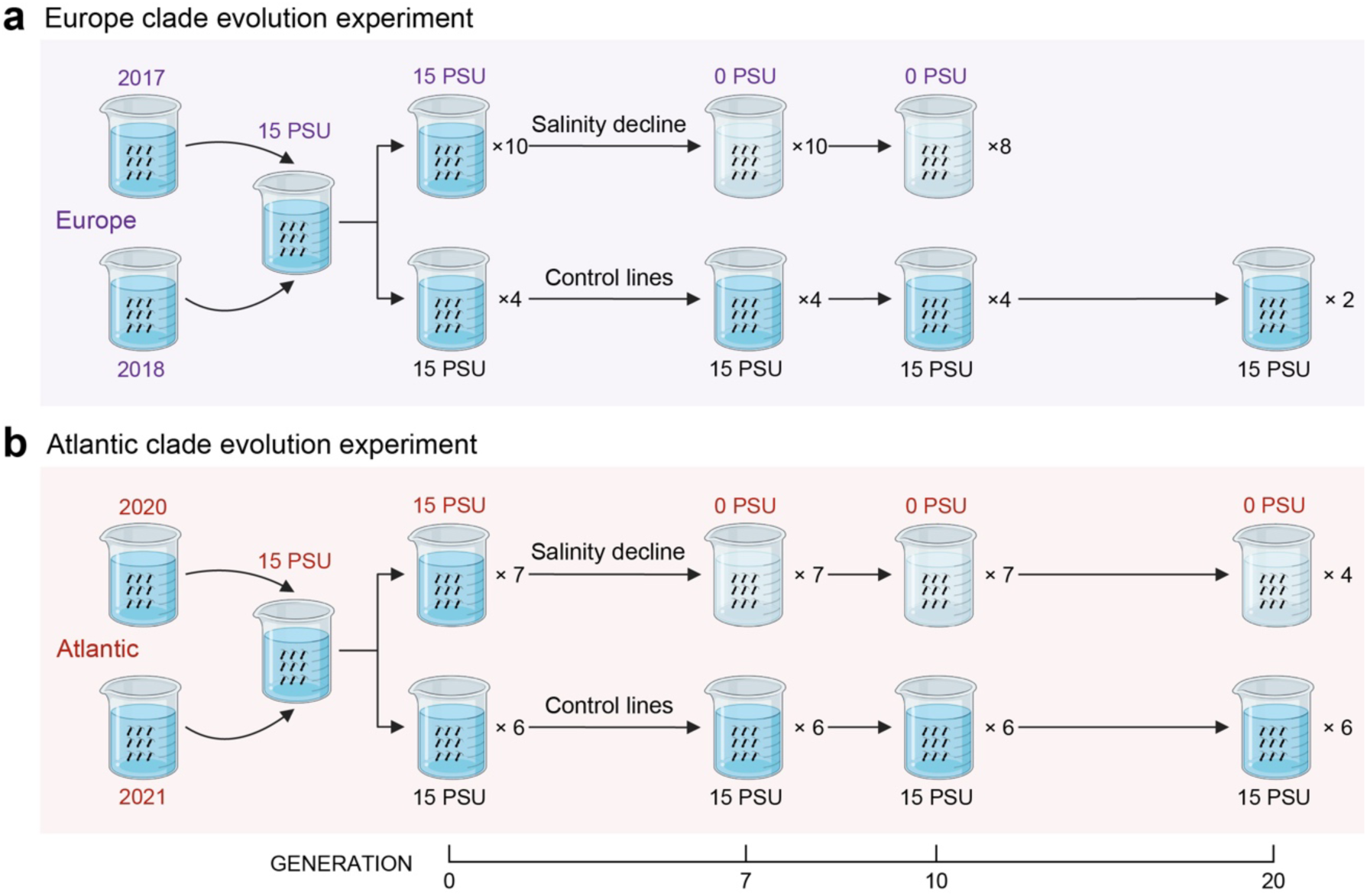
Experimental design examining selection responses to declining salinity in the two clades (sibling species) of the *Eurytemora affinis* complex. **(a)** Evolve-and-Resequence experimental design for the Europe clade population from Kiel, Germany. **(b)** Evolve-and-Resequence experimental design for the Atlantic clade population from the St. Lawrence estuarine marsh (Baie de L’Isle Verte), Quebec, Canada. For each clade, replicate lines were exposed to a stepwise salinity decline (15 → 10 → 5 → 1 → 0.1 → 0.01 → 0 PSU over six generations), and then maintained at 0 PSU until generation 10 (for the Europe clade) or generation 20 (for the Atlantic clade). Control lines for each clade were maintained at 15 PSU throughout the experiment. Selection and control lines were sampled via pooled sequencing (25 males and 25 females per pool) at generations 0 (start), 7, 10, and 20. Numbers on the right side of beakers (e.g., “×10”) indicate the number of replicate lines in each category. In total, 10 selection lines and 4 control lines were initiated in the Europe clade (a), while 7 selection lines and 6 control lines were initiated in the Atlantic clade (b). Due to occasional extinctions of beakers, eight replicate selection lines remained by generation 10 in the Europe clade (a), and four selection lines remained by generation 20 in the Atlantic clade (b). All surviving selection and control lines were included in the genomic analyses.

Given the sharp difference in genome architecture between the Europe and Atlantic clades and the fact that the four chromosomes of the Atlantic clade arose from fusions of the fifteen Europe clade chromosomes, we hypothesized that the two clades would exhibit divergent selection responses. Specifically, we hypothesized that chromosome number, the resulting recombination rate, and the positions of adaptive alleles on the chromosomes would result in varying responses to selection and differing degrees of parallel evolution. We predicted that greater chromosome number and higher recombination rate, such as those of the Europe clade, would enable positive epistatic interactions among many unlinked loci to drive highly parallel responses across replicate lines, as we observed in our previous study^4^. In contrast, we hypothesized that selection on the Atlantic clade lines would act on large haplotypes with linked loci, often brought together by chromosomal fusions at regions of reduced recombination^29^. In addition, given our prior results of signatures of balancing selection in ancestral saline populations^13^, we predicted that haplotypes with linked beneficial alleles would already be present in the native range. These pre-existing haplotypes on different genomic backgrounds could be favored during habitat change, consistent with soft selective sweeps that produce weaker selection signals and lower levels of parallelism. This effect would be magnified in genomes with longer chromosomes due to greater linkage with neutral polymorphic sites^12,32^.

Parallel evolution has long been a central topic of fundamental importance in evolutionary biology^6,48,49^. The degree of parallel evolution during replicate events provides insights into the extent to which evolutionary pathways are labile or constrained. This study is noteworthy for examining the impacts of genome architecture on the extent of parallel evolution and exploring the mechanisms of selection involved. By combining laboratory experimental evolution with whole-genome time-series data, as well as forward simulations of our data, our study directly links chromosome number to the mechanisms and predictability of polygenic adaptation. Previous experimental evolution studies have focused on species or populations with similar genome architectures, especially *Drosophila*^16–18,50,51^. In this study, the sharp contrast in karyotypes between pre- and post-fusion genomes of closely related sibling species provides a unique opportunity to dissect and gain novel insights into how genome architecture shapes evolutionary trajectories. Moreover, dissecting the genomic mechanisms underlying rapid freshwater adaptation is highly relevant in light of the sharp declines in coastal salinity at high latitudes, driven by global climate change^52,53^.

## Results and Discussion

### Rapid polygenic selection response to declining salinity in two copepod clades

We subjected laboratory selection lines from two *E. affinis* complex clades that differ in haploid chromosome number (15 in *E. affinis* proper versus 4 in *E. carolleeae*) to rapid salinity decline across 10–20 generations (Figure 2). Our experimental design introduced salinity reductions by gradually transitioning individuals from brackish water (15 practical salinity units, PSU) to fresh water (0 PSU, Lake Michigan water, ∼300 µS/cm). Multiple replicate lines (7–10 per clade; Figure 2a, b) were established from outbred wild populations, from Kiel, Germany, for the Europe clade (*E. affinis* proper) and from St. Lawrence marsh (Baie de L’Isle Verte), Quebec, Canada, for the Atlantic clade (*E. carolleeae*) (Figure 1a). Salinity was gradually reduced each generation from 15 PSU to 0 PSU over six generations (Figure 2a, b; see Methods), imposing strong selection for freshwater tolerance while avoiding immediate extinction. Thereafter, populations were maintained under freshwater conditions for up to 10 generations in the Europe clade and 20 generations in the Atlantic clade selection lines. Replicate control lines for each clade were maintained at constant salinity (15 PSU) to account for potential laboratory drift or beaker effects.

Selection lines from both the Europe and Atlantic clades showed rapid genome-wide selection responses to declining salinity (Figure 2). We identified allele frequency changes in replicate selection lines by sequencing pooled genomic DNA from each replicate at multiple time points. To identify candidate SNPs under selection, we applied Cochran–Mantel–Haenszel (CMH) and Chi-square tests, as well as linear mixed models (LMMs), to detect SNPs that deviated from neutral expectations in the selection lines and SNPs that diverged significantly between selection and control lines. These methods yielded 43,792 single-nucleotide polymorphisms (SNPs) (out of 474,041 SNPs) in the Europe clade and 9,417 SNPs (out of 195,136 SNPs) in the Atlantic clade (Supplementary Table S1).

To account for genetic linkage among the SNPs, we grouped candidate SNPs with signatures of selection into “haplotype blocks”^17,54^. These haplotype blocks under selection, treated as “selected alleles” in this study, consisted of proximate SNPs with correlated allele frequency shifts. We tracked the trajectory of each selected allele (haplotype block) via the median SNP frequency within that allele. In total, the Europe clade selection lines showed significant allele frequency changes at 10,224 SNPs, clustered into 261 selected alleles (Supplementary Table S2; Supplementary Figures 1–15). On the other hand, the Atlantic clade lines showed significant allele frequency changes at 2,383 SNPs, clustered into 66 selected alleles (Supplementary Table S3; Supplementary Figures 16–19). Control lines remained essentially unchanged at these loci, with the mean allele frequency changes not deviating from the neutral drift simulations (pink shaded areas in Figure 3a, b).

**Figure 3.**
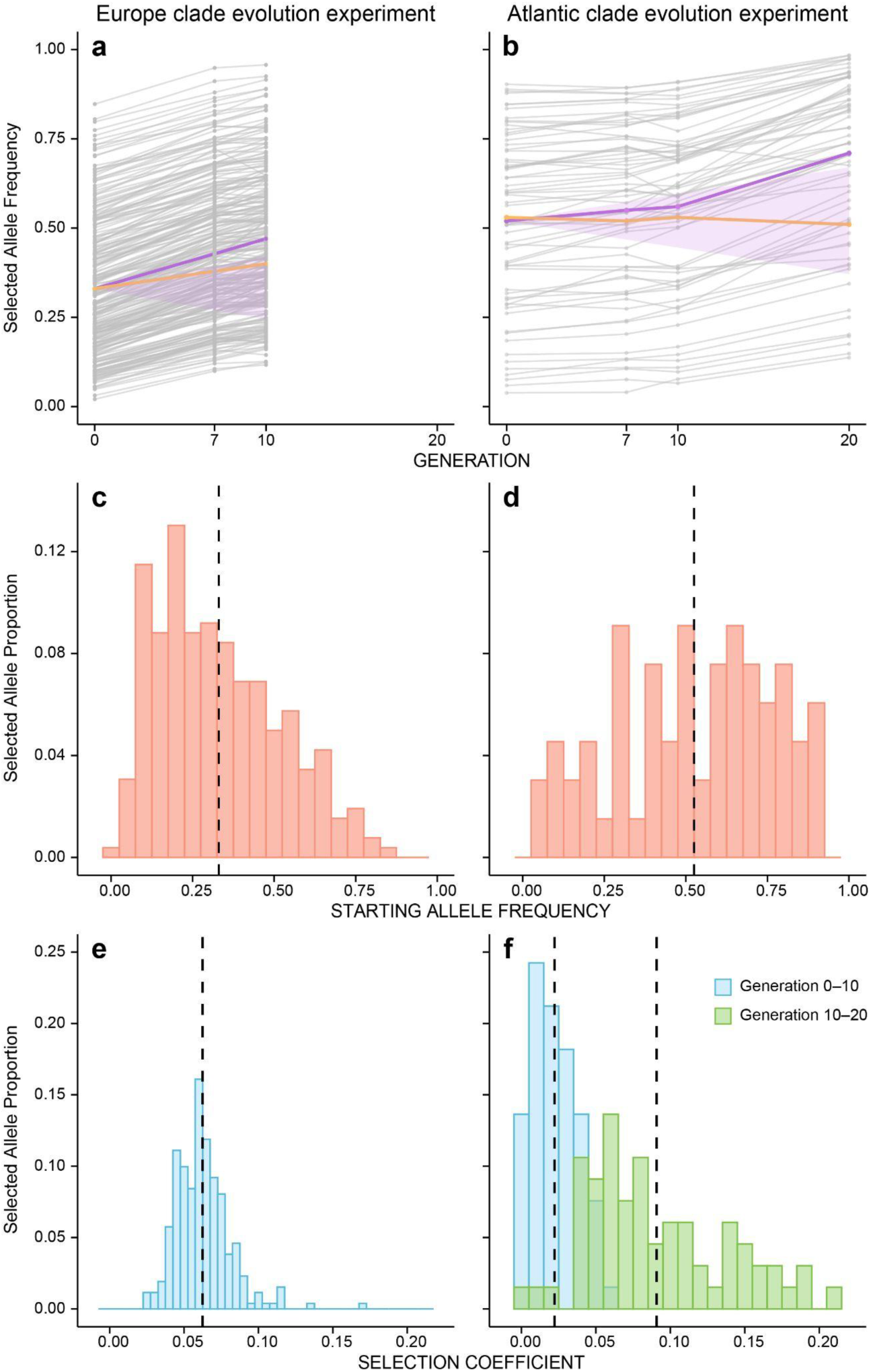
Laboratory selection responses of clades (sibling species) that differ in chromosome number from the *Eurytemora affinis* complex. The figure shows selection responses under salinity decline for the Europe clade (left graphs, 15 chromosomes) and Atlantic clade (right graphs, 4 chromosomes). **(a, b)** Allele frequency trajectories of selected alleles (haplotype blocks) during laboratory selection. The Europe clade (a) contains 261 selected alleles, whereas the Atlantic clade (b) contains 66 selected alleles. The gray lines represent mean allele frequencies for each selected allele, averaged across the replicate selection lines. The purple lines indicate the mean frequency of all selected alleles in the selection lines, whereas the yellow lines show the corresponding mean frequency in the control lines. Pink shaded areas represent the 1st and 99th percentile ranges of allele frequencies from 10,000 neutral simulations, initialized at the empirical starting frequency. **(c, d)** Starting frequencies of selected alleles in Europe (c) and Atlantic (d) clades, polarized to the rising allele in the selection lines. **(e, f)** Estimated selection coefficients (*s*) of selected alleles in the Europe (e) and Atlantic (f) clades. Selection coefficients were calculated from allele frequency changes over the first ten generations in both clades (blue) and additionally over Generation 10 to Generation 20 in the Atlantic clade (green). Dashed vertical lines indicate the mean selection coefficient within each distribution.

We performed Gene Ontology (GO) enrichment analyses for genes containing SNPs with signatures of selection (10,224 in the Europe clade and 2,383 in the Atlantic clade) during the evolution experiments. The enrichment analysis revealed that the selection response to declining salinity in both clades included genes related to ion-regulatory functions. In the Europe clade selection lines, enriched GO categories included several ion-binding or channel-related functions, such as ‘calcium-dependent protein binding’ (GO:0048306), ‘anion binding’ (GO:0043168), ‘ion binding’ (GO:0043167), and ‘chloride channel activity’ (GO:0005254) (Supplementary Table S4). Enriched categories in the Atlantic clade lines also included transmembrane ion transport functions, as well as regulatory aspects of transport physiology, such as ‘transepithelial transport’ (GO:0070633), ‘response to calcium ion’ (GO:0051592), ‘epithelial fluid transport’ (GO:0042045), ‘fluid transport’ (GO:0042044), and ‘regulation of transmembrane transport’ (GO:0034762) (Supplementary Table S5).

The number and size of selected alleles (haplotype blocks) indicate that the Atlantic clade genome contained much larger haplotypes of linked beneficial alleles than the Europe clade genome. Notably, the density of selected alleles per chromosome was similar between the clades, with the large Atlantic clade chromosomes and the smaller Europe clade chromosomes each containing similar numbers of selected alleles (with an average of 17 alleles per chromosome in both clades). However, the total number of selected alleles differed by about fourfold between them (261 for Europe versus 66 for Atlantic), mirroring the roughly fourfold difference in chromosome number between the clades (15 versus 4), despite their similarity in total genome size (671 Mb for Europe versus 529 Mb for Atlantic). Consequently, the Atlantic clade’s 66 selected alleles (haplotype blocks) were individually much larger (ranging 0.68–4.05 Mb, mean ∼1.72 Mb) than the 261 selected alleles of the Europe clade (0.14–1.76 Mb, mean ∼0.63 Mb) (Mann–Whitney test, *U* = 724, two-tailed *P* = 1.4 × 10^−30^). This result indicated that many adaptive SNPs in the Atlantic clade genome were physically linked within fewer large haplogroups on the same chromosome. In contrast, these haplogroups were partitioned into many smaller blocks on separate chromosomes in the Europe clade genome.

The larger haplogroups under selection in the Atlantic clade selection lines were consistent with our previous observations, which had found that adaptive loci were significantly clustered in the Atlantic clade genome^29,42^. In our prior study, we had found that the distribution of ion transport-related genes on the chromosomes deviated significantly from a uniform distribution and tended to be more clustered than expected^42^. Our comparison of the genome architecture of three clades within the *E. affinis* complex had revealed that chromosomal fusions repositioned key ion transport-related genes under selection from regions near the telomeres in the Europe clade genome into regions near the chromosome centers in the Atlantic clade genome^29^. Notably, population genomic signatures of selection associated with salinity change and linkage disequilibrium (LD) in wild populations of the Atlantic clade were also enriched at these chromosomal fusion sites, especially at the centromeres^29^. Such clustering of adaptive loci within chromosomes into potential coadapted complexes, as is the case for the Atlantic clade, would facilitate co-expression and co-inheritance of these loci as a unit^25,29^ (see next sections).

### Contrasting selection responses of clades with high- versus low-chromosome number

While both the Europe and Atlantic clade selection lines exhibited rapid evolutionary responses to laboratory salinity decline, the evolutionary trajectories underlying their selection responses were strikingly different (Figure 3). In the Europe clade selection lines, we observed an immediate and pronounced rise in the frequencies of beneficial alleles (haplotype blocks) (Figure 3a), which typically started at low frequency (Figure 3c). In contrast, the selection response of the Atlantic clade lines appeared muted at the early generations in the experiment (Figure 3b), acting on standing genetic variation of intermediate frequency alleles (Figure 3d). These contrasting selection responses were influenced by differences in the starting frequency of the beneficial alleles (Figure 3c, d).

The profoundly different selection responses between the clades suggest that differences in genome architecture between the clades led to contrasting mechanisms of selection. Specifically, the selection lines of the pre-fusion 15-chromosome Europe clade exhibited a rapid and strong selection response with a high mean selection coefficient of *s* = 0.06 ± 0.001 SE (with individual loci reaching up to *s* = 0.17 during Generations 0–10; Figure 3e). The selected alleles (haplotype blocks) had low starting frequencies at Generation 0 (mean = 0.33 ± 0.01 SE across the base experimental lines; Figure 3a, c) and then increased dramatically in frequency by Generation 10 (Figure 3a). Across the 261 selected alleles (haplotype blocks) in these selection lines, allele frequencies rose by an average of 28.7% (mean increase in absolute frequency of 0.13 ± 0.002 SE) in the selection lines by Generation 10, indicating a robust selection response.

In contrast, the selection lines of the post-fusion 4-chromosome Atlantic clade showed a minimal selection response up to Generation 10 but then ultimately exhibited a substantial rise in beneficial allele frequency by Generation 20. Specifically, within the first 10 generations of selection, the selection response of the 66 selected alleles (haplotype blocks) was minimal, showing only a 6.3% average frequency increase by Generation 10 (mean increase in absolute frequency of 0.035 ± 0.005 SE; Figure 3b). This result was consistent with the significantly lower mean selection coefficient up to Generation 10 in the Atlantic clade selection lines (mean *s* = 0.021 ± 0.002 SE; Figure 3f; blue histogram bars) relative to that of the Europe clade lines (*s* = 0.06 ± 0.001 SE) (Mann–Whitney test, *U* = 16,560, two-tailed *P* = 5.2 × 10^−31^). While the selection response appeared muted, the Atlantic clade’s selected alleles were already at intermediate to high frequencies in the replicate selection lines at Generation 0 (mean = 0.52 ± 0.03 SE in the base experimental lines; Figure 3d), relative to the lower mean starting allele frequency in the Europe clade selection lines (mean = 0.33 ± 0.01 SE) (Mann–Whitney test, *U* = 4,641, two-tailed *P* = 7.1 × 10^−9^). After Generation 10, however, the selection responses in the Atlantic clade selection lines intensified, with the adaptive alleles rising in frequency by an average of 24% from Generations 10 to 20 (mean increase in absolute frequency of 0.166 ± 0.008 SE; Figure 3b). This sharp increase in beneficial allele frequency corresponded to a significantly higher mean selection coefficient of *s* = 0.09 ± 0.006 SE (Figure 3f; green histogram bars) during the later phase of the experiment (Generations 10–20), relative to the early phase of the experiment (Generations 0–10) (paired *t*-test, *t* = 10.53, df = 65, two-tailed *P* = 1.1 × 10^−15^).

This intriguing delayed or weak selection response in the Atlantic clade during the first 10 generations of selection (Figure 3b) might be attributed to selection acting on the standing variation of beneficial alleles, consistent with the relatively high starting frequencies of beneficial alleles at Generation 0 (Figure 3d). Many of the Atlantic clade selected alleles remained at intermediate frequencies throughout Generations 0 to 10, rather than fixing. This phenomenon might be attributed to “soft selective sweeps” of beneficial alleles, in which the same beneficial variant (or functionally equivalent variants) segregates on multiple genetic backgrounds^32^. As a consequence, as a given beneficial allele rises in frequency, multiple distinct haplotypes carrying that allele can increase simultaneously, damping the shift in allele frequencies across the genome. This effect would be magnified in genomes with fewer longer chromosomes, as longer stretches of polymorphic SNPs (and linked neutral SNPs) would be associated with the beneficial alleles.

In contrast, the selection response in the Atlantic clade lines from Generations 10 to 20 (at 0 PSU) was markedly greater than the response of the previous 10 generations (see previous paragraph). By Generation 20, 22.7% (15 out of 66 alleles) of the beneficial alleles approached fixation (with a frequency > 0.9). The rapid increase in selected allele frequencies from Generations 10 to 20 in the Atlantic clade lines (24% rise in mean frequency), following the minimal response during the first 10 generations (only 6.3% rise in mean frequency), was comparable to the selection response from Generation 0 to 10 in the Europe clade (28.5% rise in mean frequency). This rapid response in the Atlantic clade lines during Generations 10 to 20 might have been facilitated by recombination and the aggregation of beneficial alleles within the genome. In the Atlantic clade genome, with fewer and longer chromosomes, recombination is expected to break down linkage disequilibrium more slowly, delaying the consolidation of beneficial alleles to form the highest-fitness haplotype.

### Adaptive architecture of different clades: genomic distribution of selected loci

The greater difficulty and more time required for recombination to break up linkages between alleles in the Atlantic clade versus the Europe clade is consistent with differences in the genomic distribution of loci under selection. Here, we refer to “adaptive architecture” as the genetic composition and organization underlying an adaptive trait^8^.

As discussed in the previous section, chromosomal fusion events might have important consequences for how populations from different clades respond to natural selection. A notable feature of these genomes is that the 15 chromosomes in the Europe clade genome fused to form 4 chromosomes in the Atlantic clade genome^29,42^. Such fusion events have reshaped genome architecture by bringing together key ion transport-related genes from telomeric regions to the fusion sites^29^. Thus, we performed location-based permutation tests to determine whether the loci favored by selection during our laboratory evolution experiment were spatially biased toward telomeric regions in the 15-chromosome genome and toward chromosomal fusion sites in the 4-chromosome genome. Our goal was to evaluate whether selected SNPs were non-randomly distributed along chromosomes, with a tendency to occur near chromosome ends (telomeres) or fusion sites, as we had previously found that selected SNPs tended to be concentrated near the fusion sites in wild populations with fused chromosomes^29^. Based on our prior results, we predicted stronger enrichment of selected SNPs near chromosome ends in the Europe clade, whereas we expected that the Atlantic clade would show enrichment near chromosome centers, particularly at fusion sites.

We tested for enrichment of SNPs in the telomeric regions of genomes of both clades and around chromosomal fusion sites in the Atlantic clade. To test for enrichment in the telomeric regions, chromosomes were partitioned into non-overlapping genomic bins of 1 Mb or 2 Mb. Telomeric regions were defined as the terminal bins at each end of a chromosome. The observed number of selected SNPs in each focal region was compared with chromosome-specific null distributions generated by random sampling of bins from the same chromosome. This procedure was repeated 10^6^ times to obtain empirical right-tailed *P* values. We applied the same framework to test for enrichment around chromosomal fusion sites in the Atlantic clade using bins centered on fusion midpoints and comparing the empirical distribution to the chromosome-specific permutation null distributions (see Methods).

In the Europe clade, selected SNPs were significantly enriched in telomere-proximal regions at both 1 Mb and 2 Mb intervals, in line with our prediction. Overall, the genome-wide telomeric SNP count was far above the null permutation and highly significant under both 1 Mb (right-tailed *P* = 1.0 × 10^−5^) and 2 Mb binning (right-tailed *P* = 1.0 × 10^−6^). Under 1 Mb binning, considering both ends jointly per chromosome, telomere enrichment was significant in 5 out of 15 chromosomes (Chr1, Chr4, Chr6, Chr8, Chr10; Supplementary Table S6; Supplementary Figure 20). Under 2 Mb binning, telomere enrichment was significant in 7 out of 15 chromosomes (Chr3, Chr4, Chr6, Chr8, Chr10, Chr12, Chr14; Supplementary Table S7; Supplementary Figure 21). These results supported our hypothesis that loci under selection in the pre-fusion, high-chromosome Europe clade were biased toward the chromosome ends, where recombination rates are typically elevated and can facilitate the assembly of favorable allele combinations across chromosomes^23,25^.

In contrast, in the Atlantic clade, we observed weaker and less consistent enrichment of selected SNPs toward the telomeres and no signal of enrichment at the chromosomal fusion sites. A direct comparison between clades, based on the proportion of selected SNPs located in telomeric bins, showed that the Europe clade consistently exhibited higher telomeric enrichment than the Atlantic clade. Under 1 Mb binning, 7.1% of selected SNPs in the Europe clade occurred in telomeric bins compared with 3.5% in the Atlantic clade (right-tailed *P* = 0.20). On the other hand, under 2 Mb binning, 14.6% of selected SNPs in the Europe clade were in telomeric bins compared with 5.1% in the Atlantic clade (right-tailed *P* = 3.15 × 10^−4^). Consistent with these patterns, genome-wide enrichment near the telomeres of the Atlantic clade showed modest but significant signals under 1 Mb binning (permutation test: right-tailed *P* = 0.00537) and marginal significance at 2 Mb binning (permutation test: right-tailed *P* = 0.01299). Telomere enrichment was significant in 2 out of 4 chromosomes under 1 Mb binning (Chr1 and Chr4; Supplementary Table S6; Supplementary Figure 22) and 1 out of 4 chromosomes under 2 Mb binning (Chr2; Supplementary Table S7; Supplementary Figure 23).

In terms of enrichment of selected SNPs at the fusion sites in the Atlantic clade, across all 11 fusion sites tested, neither individual fusion-site bins nor chromosomes showed significant enrichment at either 1 Mb or 2 Mb intervals (genome-wide fusion-site permutation test: 1 Mb right-tailed *P* = 0.710; 2 Mb right-tailed *P* = 0.604; Supplementary Table S8; Supplementary Figures 24–25). Thus, unlike the long-term selection patterns observed in natural Atlantic clade populations^29^, the chromosomal fusion sites did not constitute detectable hotspots of selected SNP enrichment during the short-term laboratory evolution in this study. This result is consistent with the limited time for recombination and selection to repeatedly target the same fusion-centered genomic regions across replicates in the short-term laboratory evolution, relative to the 300–400 generations of selection in the wild invasive populations^13,29^.

Although we did not find significant enrichment at the fusion sites in particular, the signals of selection were still enriched toward the chromosome centers in the Atlantic clade relative to the Europe clade. Using a centrality metric that quantifies how close each SNP lies to the chromosome center relative to chromosome length, we found that selected SNPs in the Atlantic clade were on average about 1.5–1.7% closer to chromosome centers than those in the Europe clade (permutation test: right-tailed *P* = 1.0 × 10^−6^). Given the markedly different chromosome length (Atlantic: mean 132 Mb per chromosome; Europe: mean 45 Mb per chromosome), this relative shift corresponds to an average distance to the chromosome centers of 2.0–2.2 Mb on the Atlantic clade chromosomes versus 0.67–0.76 Mb on the Europe clade chromosomes. These results indicate more central localization of selected SNPs toward regions of low recombination in the Atlantic genome relative to the Europe clade genome.

The differences in the chromosomal distribution of selected SNPs between the two clades suggest that these clades likely achieved rapid adaptation through different genomic mechanisms with their very different genome architectures. In the pre-fusion Europe clade genome, where selected loci are biased toward telomeres and distributed across many chromosomes, higher effective recombination can promote the repeated assembly of coadapted alleles across replicates. In such a genome, positive synergistic epistasis may have played an important role in bringing together functionally related alleles across unlinked loci distributed across many different chromosomes^4^ (see next sections; Figure 4). With the greater number of chromosomes and higher recombination rate in the Europe clade relative to the Atlantic clade, positive epistasis could drive the assembly of multiple beneficial freshwater-adapted alleles that were originally on different genetic backgrounds. Thus, positive synergistic epistasis, driving the assembly of the same sets of beneficial alleles in independent selection lines, could promote greater parallelism among replicate selection lines and drive rapid adaptation (see next sections).

**Figure 4.**
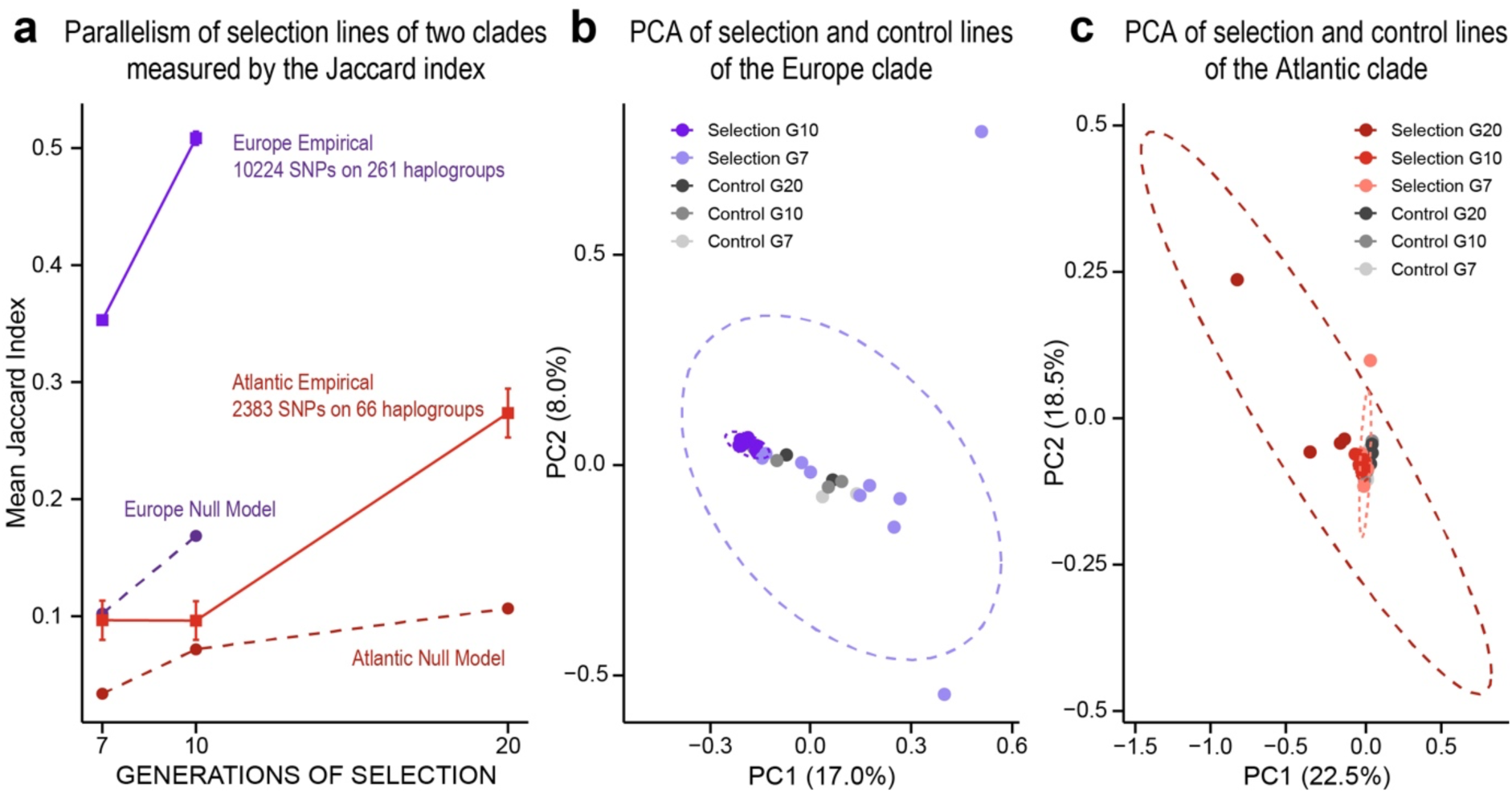
Parallel selection response of replicate selection lines from the Europe (15 chromosomes) and Atlantic (4 chromosomes) clades of the *Eurytemora affinis* complex. **(a)** Extent of parallelism indicated by the Jaccard Index for empirical data from replicate selection lines of the Europe clade (purple solid line) and Atlantic clade (red solid line) versus null expectations under a null additive model (α = 0) (dashed lines), at Generations 7, 10, and 20. Error bars represent ± SE. Parallelism in the Europe clade rose sharply by Generation 10. In contrast, the Atlantic clade showed a delayed or weak response until Generation 10, followed by a relatively rapid increase in parallelism over the next 10 generations of selection. **(b)** Principal component analysis (PCA) of the Europe clade selection response, showing trajectories of the selection lines (purple dots) and control lines (grey dots) at Generations 7 (light dots) and 10 (dark dots). Replicate selection lines at Generation 7 were dispersed, but by Generation 10, they clustered tightly and separated clearly from the control lines, indicating that parallel adaptation emerged rapidly and strongly. **(c)** Principal component analysis (PCA) of the Atlantic clade selection response, showing trajectories of the selection lines (red dots) and control lines (grey dots) at Generations 7 (light dots), 10 (medium dots), and 20 (dark dots). Selection lines at Generations 7 and 10 clustered relatively closely with each other and with the control lines, indicating shared SNP frequencies and a minimal selection response. By Generation 20, however, selection lines became more dispersed, reflecting the emergence of strong selection responses. At Generation 20, three out of four selection lines clustered closely in the PCA (c), consistent with the rise in the Jaccard index (a).

In contrast, in the post-fusion Atlantic genome with fewer chromosomes (4 large chromosomes), many of the selected loci are physically linked in large haplotype blocks^29,42^. In such a genome, lower effective recombination would be relatively ineffective at breaking apart or reshuffling those selected loci. As a result, selection would tend to act on whole blocks of coadapted alleles, linked together on chromosomes. In such a case, if one advantageous allele is present on a particular chromosome segment, that entire segment (and any other beneficial alleles on it) would undergo selection together as a unit. In other words, the Atlantic clade’s adaptation may have proceeded via selection on a few large, multi-gene haplotypes that each conferred freshwater tolerance.

In the Atlantic clade, these large selected haplotype blocks are likely pre-existing in the ancestral saline source populations of the St. Lawrence estuary (Figure 1a; red circle), given the high starting frequencies of beneficial (selected) alleles found in the starting population (Figure 3d). In addition, our prior study had found that the wild saline populations of the Atlantic and Gulf clades residing in the ancestral native range showed genomic signatures of balancing selection for a large proportion of SNPs under positive directional selection during saline-to-freshwater invasions^13^. These selected alleles might have been preserved in the native saline population through long-term balancing selection, maintaining both saline-adapted (higher salinity) and freshwater-adapted (low salinity) alleles through seasonal fluctuations in salinity (e.g., from 5 PSU to 40 PSU in some salt marsh habitats)^1,13,46^ across copepod generations (with a short generation time of ∼20 days)^55,56^. Maintaining variation in large linked sets of adaptive alleles could be advantageous in response to rapid environmental change. In effect, the Atlantic clade’s low chromosome number, combined with long-term balancing selection in heterogeneous estuarine environments, may have predisposed it to retain large blocks of coadapted alleles that enable rapid responses to habitat change, even though these blocks have been slowly formed by recombination and selection.

### Contrasting extent of parallel selection response in different clades

The repeatability of genomic responses to selection (parallel evolution) differed dramatically between the selection lines of the Europe and Atlantic clades, despite both clades showing a highly polygenic response to salinity decline (Figure 3). The pre-fusion 15-chromosome Europe clade showed a highly parallel genomic response to salinity decline across independent replicates, with many of the same alleles consistently favored across the selection lines (Figure 4a, see details below). In contrast, the post-fusion 4-chromosome Atlantic clade showed a much lower extent of parallelism among the selection lines than the Europe clade, particularly during the first 10 generations of selection (Figure 4a), consistent with the muted selection response in the first 10 generations (Figure 3b, previous section). Here, we define “parallel selection” as the case in which the same allele significantly increases in frequency across multiple replicate selection lines. Greater “parallelism” would refer to the same allele increasing in frequency in a greater number of selection lines.

To quantify the extent of parallelism of selection response between selection lines from the two clades, we applied three complementary statistical approaches: (1) the Jaccard index, to compare pairwise overlap of selected alleles between replicate lines relative to null simulations of our data, (2) linear model-based slope analysis, to test whether allele frequency changes were consistent in direction and magnitude across replicates, and (3) principal component analysis (PCA), to assess the clustering of replicate lines based on their genome-wide allele frequency changes.

To determine the degree of parallelism among replicate selection lines, we first compared the empirical pairwise Jaccard index to the distribution of Jaccard indices generated under a null expectation of independent allele-frequency changes^57^. For both clades, we found that the empirical Jaccard index at a given generation (Figure 4a, solid lines) was significantly greater than the null expectation (Figure 4a, dashed lines) (Mann–Whitney test, all pairwise contrasts two-tailed *P* < 0.05; Supplementary Table S9). Thus, even the Atlantic clade selection lines exhibited significantly greater parallel evolution than the null expectation, despite much lower parallelism than the Europe clade (see next paragraph) (Supplementary Table S9).

Using the Jaccard index, we found pronounced differences in the extent of parallelism between selection lines of the two clades. We considered a selected allele to be putatively adaptive in a given selection line if its allele-frequency increase exceeded the 99.9th percentile of neutral drift simulations (evaluated at Generations 7 and 10 in the Europe clade, and at Generations 7, 10, and 20 in the Atlantic clade). In the Europe clade, by Generation 10, an average of 64.5% of adaptive alleles that rose in frequency in one replicate also increased in at least one other replicate, corresponding to a mean Jaccard index of 0.508 ± 0.006 SE across the eight replicate lines (Figure 4a; Supplementary Table S9). In stark contrast, only 9.1% of adaptive alleles were shared between Atlantic clade selection lines at Generation 10 (mean Jaccard index = 0.0962 ± 0.02 SE; Supplementary Table S9). By Generation 20 in the Atlantic clade, parallelism increased to 37.1% pairwise overlap between lines (mean Jaccard index = 0.274 ± 0.02 SE; Supplementary Table S9). While this level of parallelism was significantly greater than the null expectation (Figure 4a; Supplementary Table S9), this value (37.1% overlap) remained far below pairwise overlap of the Europe clade selection lines at Generation 10 (64.5% overlap), or even the Europe clade at Generation 7 (42.6% overlap; mean Jaccard index = 0.353 ± 0.004 SE; Figure 4a; Supplementary Table S9).

To further quantify the degree of parallelism among replicate selection lines, we applied a linear-model approach to compare the evolutionary trajectories of selected alleles across replicate lines^58^. Specifically, for each allele in each replicate selection line, we fitted a linear model of allele frequency as a function of generation, and measured the regression slope as the rate of allele frequency change over time (see Methods). To account for neutral sampling noise and genetic drift, we compared each observed slope to the distribution of slopes obtained from neutral drift simulations (up to Generation 10 in the Europe clade and up to Generation 20 in the Atlantic clade). Alleles for which slopes were significantly greater than the neutral expectation (*P* < 0.05) in all replicate selection lines were classified as putatively adaptive alleles. To evaluate whether replicate trajectories were fully parallel across replicate selection lines, we then tested for slope heterogeneity across replicate lines using a Chi-square test, with Bonferroni-adjusted confidence intervals as a conservative alternative criterion. Alleles showing significant among-replicate slope heterogeneity were classified as non-parallel. The proportion of fully parallel alleles was calculated as the fraction of loci that were (i) putatively adaptive in all replicates and (ii) not classified as non-parallel, divided by the total number of loci tested.

Consistent with the Jaccard index results, the linear-model analysis indicated a much higher degree of parallelism in the Europe clade than in the Atlantic clade. In the Europe clade, 50.2% of alleles were putatively adaptive across all replicates by Generation 10, while only 6.9% (Chi-square) and 5.0% (Bonferroni corrected) were classified as non-parallel. Thus, 43.3–45.2% of alleles exhibited fully parallel frequency shifts across replicate selection lines. By contrast, in the Atlantic clade, only 21.2% of alleles were putatively adaptive across all replicate lines by Generation 20, and none were classified as non-parallel by either the Chi-square or Bonferroni criteria.

The PCA analysis further supported the contrasting patterns of parallelism between the clades and revealed the temporal dynamics of parallelism among selection lines in both clades (Figure 4b, c). The trajectories of the Europe clade selection lines across generations in the PCA plots revealed early dispersion of the selection lines at Generation 7, followed by increasing convergence by Generation 10 (Figure 4b). By Generation 10, the eight selection lines clustered tightly together, clearly diverging both from the control lines (MANOVA, Wilks’ λ = 0.17, F(6, 22) = 5.28, *P* = 0.0017) and from the Generation 7 selection lines (MANOVA, Wilks’ λ = 0.33, F(2, 9) = 6.89, *P* = 0.0025) (Figure 4b). This result indicates that parallel adaptation emerged rapidly and strongly after only a few generations (Figure 3a). This convergence among Europe clade selection lines in the PCA was consistent with the high Jaccard index (Figure 4a) and similarity in slope among selection line trajectories (see previous paragraph).

In contrast, the PCA plot of the Atlantic clade selection lines exhibited a highly divergent selection response from the Europe clade selection lines. For the Atlantic clade, the selection lines at Generations 7 and 10 clustered tightly together in the PCA plot (Figure 4c), consistent with their shared initial genetic composition. This result was concordant with the minimal selection response, in terms of allele frequency shifts, in the Atlantic clade selection lines from Generations 0 to 10 (Figure 3b; Supplementary Figure 26) and selection on standing genetic variation (Figure 3d). In addition, the minimal separation between Generations 7 and 10 (MANOVA, Wilks’ λ = 0.64, F(2, 11) = 3.16, *P* = 0.083) was consistent with the low levels of parallelism as indicated by the Jaccard index (Figure 4a), presumably due to the minimal selection response. In contrast, by Generation 20, the selection lines diverged from both Generation 7 (MANOVA, Wilks’ λ = 0.42, F(2, 8) = 5.43, *P* = 0.032) and Generation 10 (MANOVA, Wilks’ λ = 0.42, F(2, 8) = 5.43, *P* = 0.032), and no longer overlapped in distribution with the control lines (MANOVA, Wilks’ λ = 0.30, F(6, 34) = 4.68, *P* = 0.0014), reflecting strong selection responses (Figure 4c). While the selection lines at Generation 20 appeared dispersed in the PCA (Figure 4c), three out of four selection lines clustered closely together, consistent with the rise in the Jaccard index and increased parallel response (Figure 4a).

Notably, the Europe clade achieved a very high degree of parallelism (Figure 4a, b) despite the selection response involving a greater number of alleles (261 for Europe versus 66 for Atlantic clade). Both theoretical expectations and empirical evidence in *Drosophila* suggested that parallelism should decline as greater numbers of loci contribute to adaptation of a polygenic trait^8,17,59^. As such, the Europe clade’s unexpectedly strong parallelism suggests that some other mechanisms contribute to the greater parallelism across replicate lines than expected (see next section). By contrast, the Atlantic clade’s genome architecture with fewer large chromosomes might have favored selection on alternative linked loci, promoting replicate-specific trajectories and reducing parallelism (see next paragraph).

Furthermore, differences between the clades in the mode of selection and linkage structure likely contributed to the contrasting levels of parallelism between the clades. The Europe clade’s strongly parallel genomic response was consistent with the increase of initially rare or low-frequency beneficial alleles, producing relatively uniform selection signals across replicates. In contrast, adaptation in the Atlantic clade likely proceeded through soft selective sweeps arising from standing genetic variation, where pre-existing beneficial alleles are associated with multiple alternative genomic backgrounds. As such, selection favoring the beneficial alleles could drive selection to favor multiple linked haplotypes^32,60,61^. Consequently, selection from standing variation inherently reduces both the strength of selection signals and the degree of parallelism, as replicate lines can fix different haplotypes carrying alternative beneficial haplotypes with different sets of linked neutral alleles. Thus, with standing variation, beneficial alleles are often in linkage disequilibrium with polymorphic neutral sites. In species with large chromosomes and extended haplotype blocks, these linked neutral variants are co-inherited, further diminishing the consistency of allele frequency changes across the replicate selection events.

Over longer evolutionary timescales, however, recombination is expected to erode linkage disequilibrium and decouple adaptive alleles from their neutral backgrounds even in the Atlantic clade populations, potentially restoring parallelism as selection repeatedly targets the same loci. This process may explain the minimal selection response in the Atlantic clade selection lines for 10 generations of laboratory selection, followed by the rise in beneficial allele frequency and increased parallelism thereafter (Figures 3b and 4). In contrast, the opportunities for recombination over many generations across several decades might explain why natural populations from the Atlantic clade show strong parallel genomic signatures of selection across independent saline-to-freshwater invasions^13^.

### Role of positive epistasis in driving parallel evolution

The divergent extent of parallel evolution between the clades (previous section) raises questions regarding the mechanisms that drive parallel evolution in these clades. As mentioned above, our prior experimental results^4^ had revealed that the high degree of parallelism among selection lines in the Europe clade was consistent with positive epistasis among the alleles favored by selection^62,63^. In contrast, in the Atlantic clade, low recombination and large linkage blocks would make it less likely that the repeated assembly of the same beneficial allelic combinations across replicates would drive parallelism among the selection lines (see previous section). To explore the potential role of positive epistasis in promoting parallel selection within each clade, we performed theoretical simulations of our data under a model of positive synergistic epistasis with varying strengths of epistasis^4,64^.

We performed forward genetic simulations in SLiM 3^65^ to test whether positive synergistic epistasis could account for the observed patterns of parallelism in our selection lines. We parameterized the simulations to match our experimental design, with 261 loci distributed across 15 chromosomes for the Europe clade and 66 loci across 4 chromosomes for the Atlantic clade, each with 10 and 7 replicate populations, respectively. After each simulation run, we calculated the Jaccard index of selected alleles across replicates to quantify the extent of parallelism under alternative genetic models. We varied the strength of epistasis using the epistasis coefficient (α), which indicates the extent to which the combined effects of beneficial alleles confer fitness advantages that exceed additive expectations. That is, a larger α indicates a greater positive synergistic epistasis effect. We used Approximate Bayesian Computation (ABC) to identify the α values that best reproduced the empirical mean Jaccard indices observed among replicate selection lines at the corresponding generations.

In the Europe clade, the model with a large epistasis coefficient (α = 53) was able to closely reproduce the degree of parallelism in our empirical data (mean Jaccard index of 0.508 ± 0.006 SE; Supplementary Table S9), including the sharp rise in parallelism at Generation 10 (Figure 5a). In contrast, simulations under the additive null model (α = 0, no epistasis) consistently underestimated the levels of empirical parallelism, with Jaccard indices falling far below the observed values at Generations 7 and 10 (Figure 5a). The degree of parallelism of the model with high positive epistasis (α = 53) was significantly greater than the model with no epistasis (α = 0), both at Generation 7 (Mann–Whitney test, *U* = 1,000,000, two-tailed *P* < 0.0001; Supplementary Table S10) and Generation 10 (Mann–Whitney test, *U* = 1,000,000, two-tailed *P* < 0.0001; Supplementary Table S10) (Figure 5a). These results suggest that selection in the Europe clade lines occurred in a regime in which groups of alleles interacted synergistically, conferring large collective benefits and driving the same sets of alleles to high frequency across replicates, promoting parallelism.

**Figure 5.**
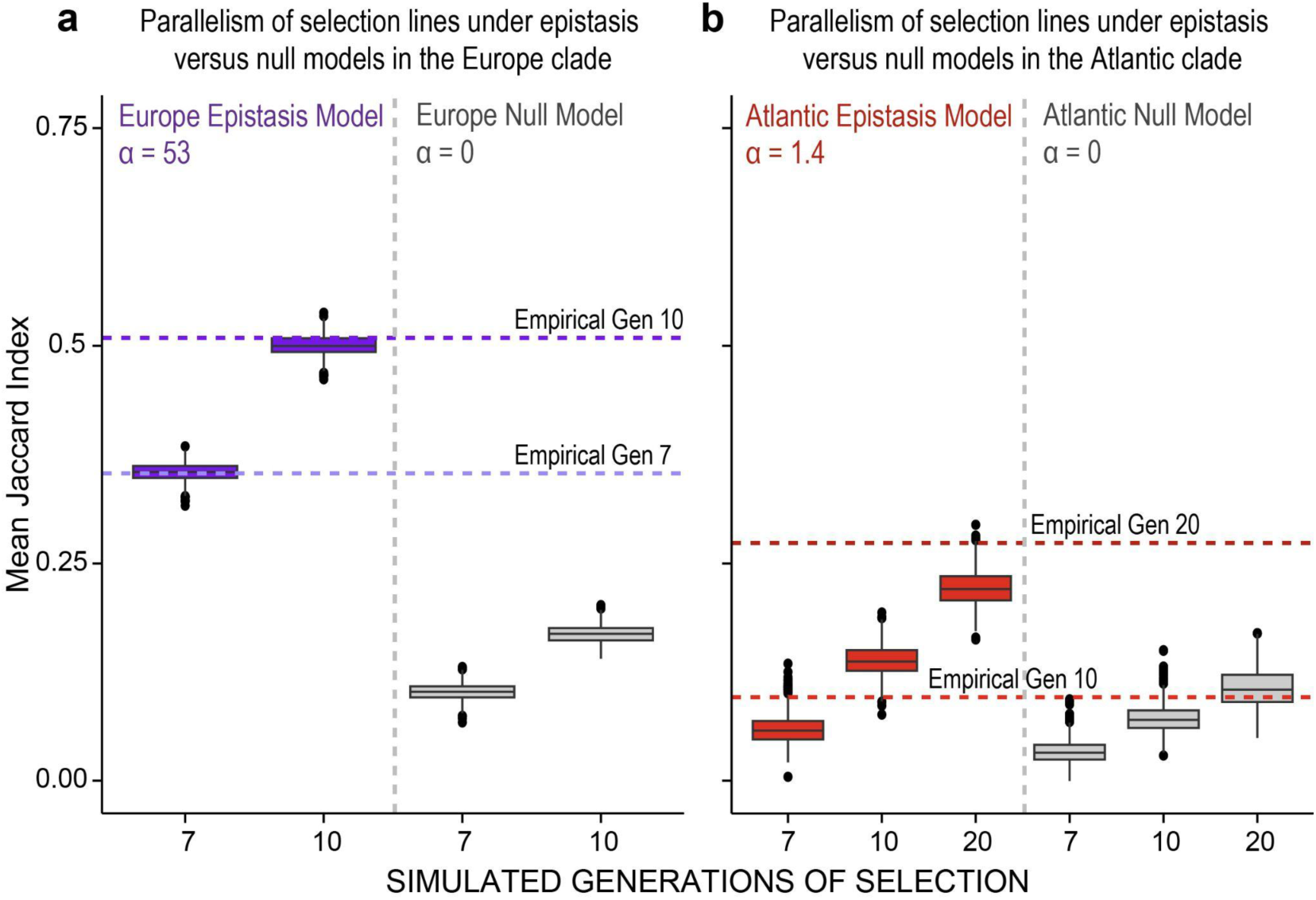
Role of positive epistasis in driving parallel evolution in selection lines of the Europe and Atlantic clades of the *Eurytemora affinis* complex. **(a)** Simulation results for the Europe clade selection lines under a positive synergistic epistasis model (α = 53; left) and a null additive model (α = 0; right). Only a high epistasis coefficient reproduced the high Jaccard indices in the model (purple boxes) that matched the empirical results of the selection lines (horizontal purple dashed lines), particularly at Generation 10. **(b)** Simulation results for the Atlantic clade under a low epistasis coefficient (α = 1.4; left) and a null additive model (α = 0; right). The low epistasis coefficient values at Generation 10 and 20 resulted in relatively low Jaccard indices in the model (red boxes), resembling the low values of the selection lines (horizontal red dashed lines). Simulations were parameterized to match the experimental design, with 1,000 replicate runs per scenario. Within each clade, all contrasts showed that epistasis models and null models differed significantly in their Jaccard index distributions at a given generation (Mann–Whitney tests, two-tailed *P* < 0.0001). Detailed pairwise comparisons and statistical results are provided in Supplementary Table S10.

By contrast, in the Atlantic clade lines, the epistasis model with a low epistasis coefficient (α = 1.4) was sufficient to match the level of parallelism in the empirical data at Generation 10 (mean Jaccard index of 0.0962 ± 0.02 SE; Supplementary Table S9) and Generation 20 (mean Jaccard index of 0.274 ± 0.02 SE; Supplementary Table S9) (Figure 5b). The best-fit α value (α = 1.4) for the Atlantic clade was therefore much lower than that for the Europe clade (α = 53), suggesting that synergistic interactions among alleles contributed little to the selection response. Instead, in this low-epistasis regime, the observed modest parallelism was more parsimoniously explained by selection acting on standing variation of alternative linked haplotypes, whose frequencies change largely as blocks rather than as recombined multi-locus genotypes.

### Effects of chromosome number and epistasis on the repeatability of adaptation

Our empirical results indicate that genome architecture, in particular chromosome number, can affect the parallelism of selection responses during replicate events (Figure 5a, b). Given the large difference in chromosome number between the Atlantic and Europe clades, we sought to quantify how chromosome number, along with epistasis, impacts the repeatability of adaptation. As such, we performed additional simulations that varied both chromosome number and the strength of positive synergistic epistasis, and examined their effects on the Jaccard index of replicate selection lines. These simulations revealed that chromosome number impacted the repeatability of polygenic adaptation in a strongly epistasis-dependent manner, as predicted by theory on multilocus selection and linkage^25,66,67^.

We simulated genomes containing 640 loci, of which 160 were designated as loci under selection and 480 as neutral. The number of selected loci (N = 160) was intermediate between the empirically identified haplotype blocks in the Atlantic (N = 66) and Europe (N = 261) clades. This ratio of 0.25 between selected and total loci was derived from the fraction of the genome length containing the selected haplotype blocks out of the total genome length for the two clades (Atlantic clade: 113.4 Mb out of 529 Mb; Europe clade: 165.2 Mb out of 671 Mb). Selection coefficients of beneficial alleles were drawn from an exponential distribution with mean 0.1 (rate parameter λ = 10), and initial allele frequencies were drawn from a uniform distribution between 0 and 1. We examined four genome architectures with 1, 4, 7, or 15 chromosomes. These values corresponded to the extreme case of complete linkage (one chromosome) to karyotypes observed in the *E. affinis* complex, with 4 chromosomes in the Atlantic clade, 7 in the Gulf clade, and 15 in the Europe clade^29,42^. We further tested six epistasis coefficients of varying strengths (α = 0, 1.4, 4, 16, 53, and 64), including the values inferred from the empirical data (α = 1.4 for Atlantic and α = 53 for Europe). After each simulation run, we calculated the mean Jaccard index of selected alleles across replicate lines to quantify the degree of parallelism at Generations 10 and 20.

To evaluate whether the effect of epistasis on parallelism depends on genome architecture, we analyzed the simulation results using a linear model (*Jaccard ∼ Chromosome number × α + Generation*), treating chromosome number as a categorical factor, the epistasis coefficient α as a continuous predictor, and Jaccard Index as the response variable. Type II ANOVA tests of this linear model showed that both chromosome number (*F* = 859.3, *P* < 0.0001) and the strength of positive synergistic epistasis (α) (*F* = 308885.2, *P* < 0.0001) had significant effects on parallelism (Jaccard Index) among the selection lines. In addition, this analysis revealed a highly significant interaction effect between chromosome number and the epistasis coefficient (α) (*F* = 734.6, *P* < 0.0001). This result indicated that chromosome number and epistasis influenced each other’s impact on parallelism, independent of generation effects. This interaction effect was also highly significant when the analysis was performed separately for each generation (Type II ANOVA; Generation 10: *F* = 404.7, *P* < 0.0001; Generation 20: *F* = 380.2, *P* < 0.0001). Thus, both chromosome number and positive epistasis can significantly enhance the extent of parallelism among replicate lines. Notably, chromosome number and the extent of epistasis have non-additive positive effects on each other, with the effect of epistasis (higher α) on parallelism being magnified with higher chromosome number, and vice versa (Figure 6). This result emphasizes that greater numbers of chromosomes (and greater recombination) and greater epistasis can together have non-additive amplifying effects in promoting parallelism, potentially by repeatedly facilitating the assembly of the same sets of coadapted allelic combinations.

**Figure 6.**
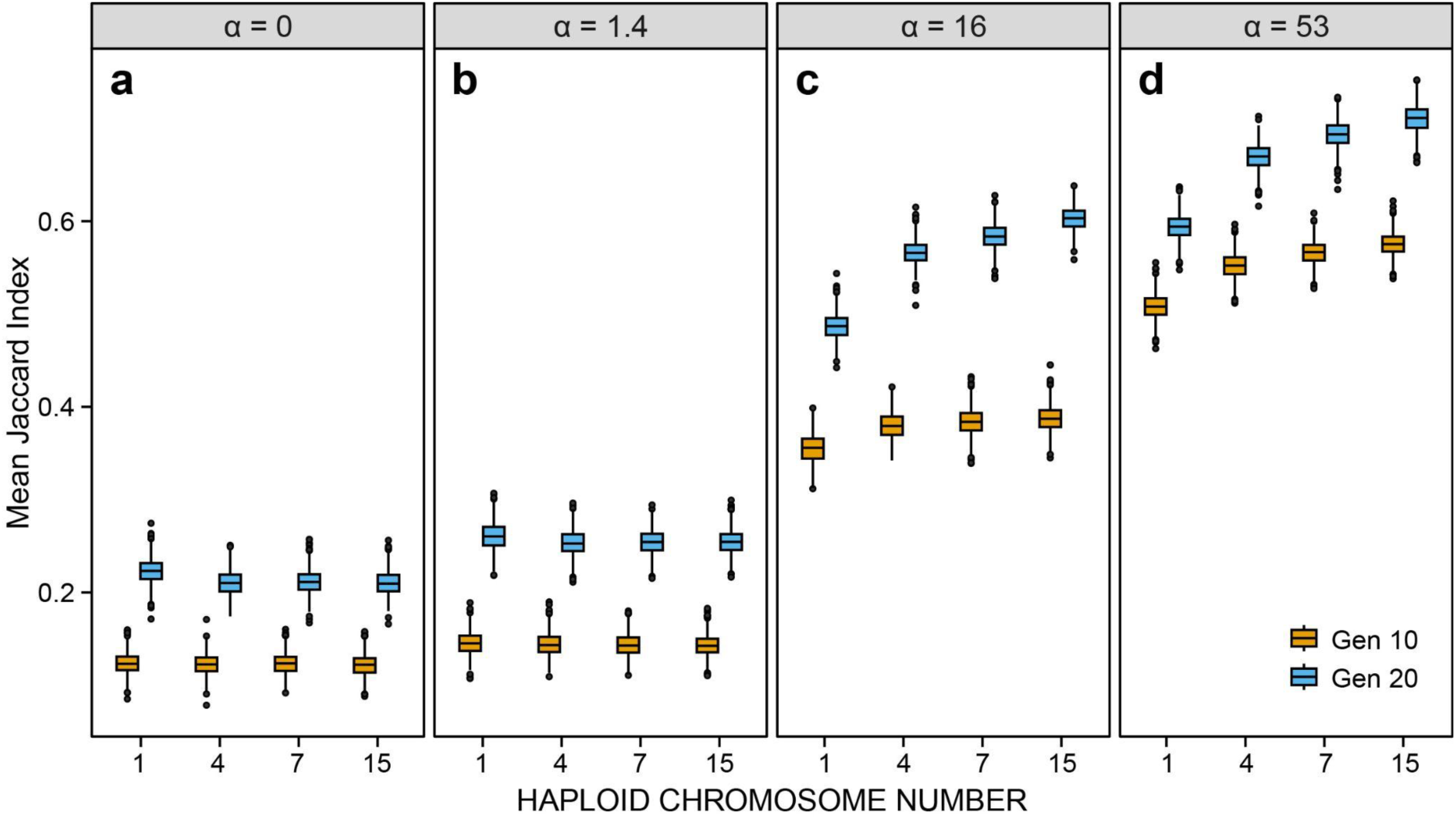
Effects of haploid chromosome number and positive synergistic epistasis on the repeatability of polygenic selection response (parallelism). Simulation results show the combined effects of chromosome number and epistasis strength on parallelism in selection response, measured as the mean Jaccard index across replicate populations (pairwise similarity between selection lines). The four panels correspond to different levels of positive epistasis: **(a)** α = 0 (additive, no epistasis), **(b)** α = 1.4 (weak epistasis, estimated for the Atlantic clade), **(c)** α = 16 (moderate epistasis), and **(d)** α = 53 (strong epistasis, estimated for the Europe clade). Within each panel, boxplots show Jaccard indices at Generation 10 (yellow) and Generation 20 (blue) across genome architectures with 1, 4, 7, or 15 chromosomes. At low or no epistasis (α = 0 or 1.4), chromosome number had only minor effects on parallelism, although extremely low chromosome counts (one chromosome) produced slightly higher values. At higher levels of epistasis (α = 16 or 53), parallelism increased markedly with higher chromosome number. When α = 16 and 53, all chromosome numbers differed significantly from one another in their Jaccard index distributions (Mann–Whitney tests, two-tailed *P* < 0.0001). Detailed pairwise comparisons and statistical results are provided in Supplementary Table S11. Additional results for α = 4 and α = 64 are also shown in Supplementary Figure 27.

Across all simulation scenarios, parallelism increased from Generation 10 to Generation 20 (Figure 6; Supplementary Table S11), consistent with the expected convergence of replicate lines under continued selection^17,59^. However, the direction and magnitude of chromosome number effects differed strikingly depending on the strength of epistasis. Under no epistasis (α = 0) or very weak epistasis (α = 1.4), differences in the Jaccard index among chromosome numbers were limited. Significant differences were apparent only between a fully linked genome (one chromosome) and the multi-chromosome genomes (Mann–Whitney test, two-tailed *P* < 0.0001; Supplementary Table S11) (Figure 6a, b). Thus, under no or low epistasis, increasing chromosome number did not increase parallelism. These results indicate that when selection is effectively additive, recombination provides few additional benefits for generating parallel responses^66,67^.

In contrast, under moderate to strong epistasis (e.g., α ≥ 16), chromosome number became a significant determinant of repeatability. Under these scenarios, the Jaccard index increased monotonically with increasing chromosome number, such that 15-chromosome genomes consistently exhibited the highest Jaccard indices (Mann–Whitney test, two-tailed *P* < 0.0001; Supplementary Table S11) (Figure 6c, d; Supplementary Figure 27). These results suggest that when synergistic positive epistasis is strong, recombination among multiple chromosomes enables replicate lines to repeatedly assemble similar coadapted multi-locus genotypes, thereby enhancing parallelism among replicate lines.

Thus, the significant chromosome number × epistasis interaction provides a mechanistic explanation for the contrasting empirical patterns between clades. In the Europe clade selection lines, the amplifying effects of high chromosome number with strong inferred epistasis (α = 53) likely generated a selection regime in which higher recombination and positive epistasis promoted the repeated assembly of beneficial allelic combinations and highly parallel genomic responses. By contrast, in the Atlantic clade selection lines, fewer chromosomes and weak inferred epistasis (α = 1.4) generated a selection regime in which large linked haplogroups persisted, leading to alternative evolutionary pathways, particularly in the early generations. More broadly, these results indicate that the predictability of adaptation cannot be inferred solely from chromosome number or epistasis alone, but also from their interaction.

### Conclusions

Our study demonstrates that rapid selection responses to a shared environmental stress can proceed through radically different genomic mechanisms due to different adaptive architectures, with contrasting consequences for evolutionary trajectories and repeatability. In particular, in the pre-fusion 15-chromosome Europe clade, the higher recombination, resulting from greater numbers of chromosomes, allowed synergistic positive epistasis among genomically dispersed adaptive alleles to generate highly parallel genomic responses across replicate lines. These adaptive alleles tended to be concentrated closer to the edges of the chromosomes in the Europe clade, relative to the Atlantic clade, further enhancing the effects of recombination. In contrast, in the post-fusion 4-chromosome Atlantic clade, adaptation occurred primarily through selection on alternative pre-existing haplotypes of linked adaptive alleles, producing less parallel outcomes in the short run.

In selection lines from both clades, laboratory selection acted on standing genetic variation rather than *de novo* mutations during the short timeframe of this study. However, the extent to which standing variation contributed to the selection response depended jointly on the starting frequency of beneficial alleles and the extent of linkage among the alleles within each genome. In the Europe clade, beneficial alleles generally started at lower frequencies and were distributed across many chromosomes with weaker linkage among loci. In such a case, higher recombination allowed alleles from different genomic backgrounds to repeatedly assemble into the same sets of allelic combinations across replicate lines, promoting the consistent rise of the same alleles and leading to strong parallelism. By contrast, in the Atlantic clade, beneficial alleles were already at intermediate to high frequencies and were embedded within large haplotype blocks due to extensive linkage. Consequently, selection acted largely on a multitude of pre-existing haplotypes. Thus, for the Atlantic clade lines, this soft selective sweep favored different linked allelic combinations in different lines and reduced the likelihood that the same alleles would respond to selection in parallel.

Together, these findings provide direct empirical evidence that genome architecture, particularly chromosome number and the recombination landscape, can shape the mode and mechanism of polygenic adaptation. Our study bridges population genomics and evolutionary theory by supporting a central role for synergistic epistasis and its interaction with recombination in driving parallel adaptation. Moreover, our results reveal how chromosomal fusions can constrain recombination and promote soft selective sweeps, thereby redirecting evolutionary trajectories. Given the accelerating pace of climate-driven environmental change, genome architecture may represent a key determinant of both the capacity for adaptation and its predictability. More broadly, this work emphasizes that the repeatability of evolution depends not only on the selective environment but also on the genetic and structural genomic substrate on which selection acts.

## Methods

### Experimental populations and Evolve-and-Resequence design

Evolve-and-Resequence (E&R) experiments were performed using populations from two clades of the *Eurytemora affinis* species complex, from the Europe clade (*E. affinis* proper, 15 chromosomes) and the Atlantic clade (*E. carolleeae,* 4 chromosomes). Wild copepods from the Europe clade were collected from Kiel, Germany (54°20′N, 10°09′E), from a salinity of 15 PSU, in 2017 and 2018^4^. Copepod samples from the Atlantic clade were collected from a brackish marsh at Baie de L’Isle Verte in the St. Lawrence estuary, Quebec, Canada (48°00′16″N, 69°25′01″W), where salinity fluctuates between 5 and 30 PSU, in 2020 and 2021. For each clade, approximately 1,000 individuals were combined to maximize genetic diversity and subsequently maintained in the laboratory incubators at 15 PSU (practical salinity units) and 12 °C for 3–4 generations to acclimate to laboratory conditions and promote random mating and recombination.

After acclimation, each clade’s population was subdivided into replicate lines in separate containers (beakers). For the Europe clade, two Generation 0 pooled samples (25 males and 25 females per pool) were collected before subdivision to characterize starting allele frequencies. The mixed culture was then divided into 14 beakers, with 10 designated as selection lines and 4 as control lines^4^. For the Atlantic clade, 13 beakers were subdivided, comprising 7 selection lines and 6 control lines. One generation after this subdivision (to minimize potential founder effects), Generation 0 samples were collected from each line by pooling 50 adults (25 males and 25 females per pool). For both clades, each beaker initially contained ∼500 mixed-stage copepods derived from the acclimated population.

Selection lines in each clade were subjected to a stepwise salinity decline to impose strong natural selection for freshwater tolerance (Figure 2). Starting from 15 PSU, salinity was gradually reduced in each generation until reaching freshwater conditions (0 PSU) over six generations. Specifically, in the selection lines of both Europe and Atlantic clades, salinity was lowered from 15 PSU to 10 PSU in Generation 1, then to 5, 1, 0.1, 0.01, and finally to 0 PSU by Generation 6. This gradual protocol prevented acute osmotic shock, allowing the populations to adapt over multiple generations. After reaching 0 PSU, selection lines were maintained in fresh water for additional generations of selection.

Specifically, the Europe clade lines were maintained until Generation 10, whereas the Atlantic clade lines were continued until Generation 20. Throughout the experiment, replicate control lines for each clade were kept at a constant 15 PSU to serve as controls for laboratory drift and beaker effects.

In this study, generations were not propagated by discrete transfers. Instead, individuals were kept in the same beaker and allowed to reproduce freely under the ambient salinity (i.e., overlapping generations). A generation time of 3 weeks (∼20 days) was used for timing the salinity changes and sample collection^55,56^. All lines were maintained on a diet of cryptophyte algae, fed three times per week. The marine alga *Rhodomonas salina* was fed to the lines at moderate to high salinities (5–15 PSU), whereas the freshwater alga *Rhodomonas minuta* was used at low salinities (≤ 1 PSU), to ensure adequate nutrition without osmotic shock to the algae^68,69^. Culture saline water (5–15 PSU) was prepared with artificial sea salt (Instant Ocean, Blacksburg, VA, USA) and filtered deionized water (supplemented with 20 mg/L Primaxin to reduce microbial contamination). Natural Lake Michigan water (∼300 µS/cm conductivity) was used for 0 PSU and low salinity cultures (≤ 1 PSU). Water in all beakers was changed weekly to maintain optimal water hygiene.

To track evolutionary genomic change, time-resolved pooled sampling of each replicate line was performed over the course of the experiment (Figure 2). From each selection line and control line, a pool of 50 adults (25 males and 25 females) was collected at multiple time points for whole-genome sequencing. Specifically, for the Atlantic clade, pooled samples were collected at Generation 0 (start), in every generation from Generation 2 through Generation 7 (during the salinity transition phase and until one generation after reaching 0 PSU), and at Generation 10 and Generation 20 (after prolonged freshwater selection). For the Europe clade, pooled samples were collected at Generation 0, Generation 7 (one generation after reaching 0 PSU), and Generation 10 (four generations in fresh water)^4^. A few selection lines went extinct during the course of selection. Consequently, 8 out of 10 Europe clade selection lines survived to Generation 10, and 4 out of 7 Atlantic clade selection lines survived to Generation 20. All surviving selection lines were included in the genomic analyses. Control lines remained viable throughout the experiment and were sampled at matching time points in both clades, enabling comparisons with the selection lines (Figure 2).

### DNA extraction, library preparation, and sequencing

Genomic DNA was extracted for each pooled sample. Atlantic clade pooled samples were processed with a standard CTAB-based DNA extraction protocol^42^, while Europe clade pooled samples were extracted using a DNeasy Blood & Tissue Kit (Qiagen, Hilden, Germany)^4^. The integrity of genomic DNA was assessed by agarose gel electrophoresis, and concentrations were quantified using a Qubit 3.0 fluorometer (Thermo Fisher, Wilmington, DE, USA).

Sequencing libraries were prepared with an average insert size of ∼350 bp using TruSeq or Nextera DNA Library Prep Kits (Illumina, San Diego, CA, USA), following the manufacturer’s protocol. Libraries were paired-end sequenced (2×150 bp reads) on Illumina platforms. The Atlantic clade libraries were sequenced on an Illumina NovaSeq 6000 platform to a target read depth of ∼30× coverage per sample. The Europe clade libraries were sequenced on multiple Illumina platforms (including HiSeq 4000 and NovaSeq) with 2×100 bp reads, yielding on average ∼117 million read pairs (∼23× coverage) per pool^4^. In total, the time-series dataset comprised whole-genome pooled sequences from 114 samples in the Atlantic clade and 28 samples in the Europe clade across generations.

### Data processing and variant calling

Sequence reads from all samples were processed with a uniform bioinformatics pipeline. Adapters and low-quality bases were trimmed using Fastp v0.23.0^70^. Clean reads were mapped to the appropriate reference genome assembly for each clade using BWA-MEM v0.7.12-r1039^71^ with default parameters. The recently assembled chromosome-level genome of *E. affinis* proper (15 chromosomes, 671 Mb)^29^ was used as the reference for the Europe clade selection lines, and the chromosome-level genome of *E. carolleeae* (4 chromosomes, 529 Mb)^42^ for the Atlantic clade selection lines. Only confidently mapped reads and proper paired-end reads were retained, with duplicates removed using Picard v2.21.6 (https://github.com/broadinstitute/picard). The resulting genome mappings were sorted, converted to BAM format, and then transformed into pileup format with SAMtools v1.21^72^, discarding low-quality alignments and bases with a quality score less than 20. SNPs were called using VarScan v2.4.6^73^ and filtered with BCFtools v1.21^74^. The resulting VCF files were processed using the R package *poolfstat* v1.1.1^75^, retaining only high-quality biallelic SNPs with a minor allele frequency greater than 0.01, at least four reads for a base call, and a minimum of 10 and a maximum of 200 total read counts for all pools. In total, 474,041 SNPs were identified for the Europe clade and 195,136 SNPs for the Atlantic clade. For each high-quality SNP, allele frequencies were obtained at each time point for each selection line by counting reference (major) and alternate (minor) allele reads in the pooled sequencing data. These allele count (and frequency) data served as the basis for all downstream selection analyses.

### Selection analyses and identification of selected haplotypes

To identify genomic signatures of selection in response to declining salinity, individual SNPs showing significant allele frequency changes over generations were first identified in the selection lines. Multiple statistical tests were applied to the time-series data and to selection-versus-control comparisons to detect SNPs that deviated from neutral expectations. In particular, Cochran–Mantel–Haenszel (CMH) tests were performed using the R package^76^ to detect SNPs with frequencies that changed more than expected under random genetic drift across selection lines. Chi-square tests were also applied, using the same R package above^76^, to detect line-specific signatures of selection that deviated from genetic drift in each selection line. CMH and Chi-square test statistics were computed using estimated effective population sizes for each line (Supplementary Table S12) to calibrate *P*-values and account for genetic drift. *P*-values were then converted to *q*-values using the R package *qvalue*^77^ to correct for multiple testing.

The effective population size (*N*_e_) was estimated using the R package *poolSeq* v0.3.5^78,79^, based on changes in SNP frequencies over time for each line. This *poolSeq* framework was used to estimate *N_e_* under both the “Plan I” (dependent on census size) and “Plan II” (independent of census size) methods. Since the census size of copepods was uncertain in each beaker, a range of plausible values (500–5,000) was assumed for the Plan I calculations. The *N*_e_ estimates were computed in 1,000-SNP genomic windows, with the median value taken per line (shown in Supplementary Table S12).

To analyze divergence between selected and control lines, linear mixed models (LMMs) were used with the R package *lme4*^80^ to identify SNPs with significantly different frequency trajectories between the selection and control lines. *Generation* and *Treatment* (selection versus control lines) were designated as fixed factors, and *Line* (replicate line) as a random factor. SNP frequency change (*yi*) served as the response variable in the linear regression, with weights proportional to sequencing depth and the effective population size (*N*_e_)^81^. An arcsine square root transformation was applied to SNP frequencies to stabilize variance and reduce bias caused by varying initial frequencies^82–84^. Likelihood ratio tests (LRTs) evaluated the significance of the treatment effect on SNP frequency change from the experiment’s start, incorporating the fixed effects of generation and the random effects of line. Two models were compared: (1) *yi* ∼ *Generation* + (1|*Line*) + *εi* and (2) *yi* ∼ *Generation* + *Treatment* + *Generation*: *Treatment* + (1|*Line*) + *εi*, where *yi* represents the SNP frequency change from the initial population at the ith SNP, and *εi* is a Gaussian error accounting for coverage and sampled individual count. LRT statistics were compared to a Chi-square distribution with two degrees of freedom to determine *P*-values. These *P*-values were subsequently converted to *q*-values for multiple testing correction.

Since many neighboring SNPs are in linkage disequilibrium, nearby significant SNPs were combined into haplotype blocks to represent individual adaptive loci. A “haplotype block” was defined as a group of nearby SNPs that showed highly correlated allele frequency patterns across generations. To objectively define these haplotype blocks, the R package *haplovalidate*^54,85^ was used with data-driven parameters. The *haplovalidate* package groups SNPs by gradually increasing the correlation thresholds until SNPs on the same chromosome do not show significantly higher correlation than those on different chromosomes. This process identifies clusters of hitchhiking SNPs that change in frequency together as a single linkage unit. Each group was treated as a single haplotype block (allele) for analysis. The median allele frequency of all SNPs in a haplotype block was used to represent the haplotype block (allele) frequency at each time point.

A haplotype block (allele) was classified as “selected” (putatively adaptive) if its allele frequency changes exceeded neutral expectations. Specifically, at each time point, the haplotype (allele) frequency increase (from its ancestral frequency at Generation 0) was required to be greater than the 99.9th percentile of the null distribution of changes expected under genetic drift. This null distribution was obtained through 10,000 simulations using the R package *poolSeq*^78^. Given the haplotype’s starting frequency and an estimated *N*_e_, the distribution of allele frequency after *t* generations was computed under neutrality (Wright–Fisher drift combined with binomial sampling), and the 99.9th quantile was used as a threshold. Any haplotype block whose frequency rose above this cutoff (in at least one replicate line) by Generations 7 and 10 for the Europe clade or Generations 7, 10, and 20 for the Atlantic clade was flagged as under selection. Frequencies in the control lines were monitored in parallel to ensure that these changes were specific to the selection treatment lines.

For each selected haplotype block (allele), the selection coefficient (*s*), which represents the per-generation fitness advantage, was estimated from the allele frequency trajectory in the experiment. The allele frequency over time was modeled using a logistic regression of allele frequency change using the R package *lme4*^80^. A linear model was fitted to the logit-transformed frequency across generations for that allele, using data from all replicate lines (considered as parallel realizations). The slope provided an estimate of *s* under the assumption of approximately constant selection^78^.

Gene Ontology enrichment analyses were performed for genes containing SNPs under selection that were contained within the selected haplotype block (allele) using TBtools v1.112^86^.

### Location-based permutation tests of chromosomal localization

To test whether SNPs under selection were non-randomly distributed along chromosomes and spatially biased toward the telomeric regions in either the Europe and Atlantic clades, or towards chromosomal fusion sites in the Atlantic clade, location-based permutation tests were implemented using R. For each clade, the corresponding chromosome-level reference genome assembly and chromosome length were used^29,42^. Chromosomes were partitioned into non-overlapping genomic bins of both 1 Mb and 2 Mb, generating two independent spatial intervals. For telomere analyses, telomeric regions were defined as the terminal bins at each chromosome end in both clades. For fusion-site analyses in the Atlantic clade, bins were defined as the non-overlapping genomic bin containing the midpoint of each chromosomal fusion site (see Du et al.^29^).

For each telomeric or fusion-site bin, the observed test statistic was the number of selected SNPs located within that bin. To generate a null distribution under the hypothesis that selected loci are not spatially biased along chromosomes, chromosome-specific permutation tests were performed using R. Specifically, bins of the same size were randomly sampled from all bins on the same chromosome (including telomeric or fusion bins), and their SNP counts were recorded. This procedure was repeated 10^6^ times to generate empirical null distributions. *P*-values were calculated as the right-tail probability that a randomly sampled bin contained at least as many selected SNPs as the observed focal bin.

Both chromosome-level and genome-wide tests of telomeric or fusion-site enrichment were conducted using R. For chromosome-level tests, the observed test statistic was the sum of SNP counts across all focal bins on that chromosome, and null distributions were generated by repeatedly sampling the same number of bins from that chromosome 10^6^ times and summing their SNP counts. For genome-wide tests, null distributions were constructed using chromosome-stratified permutation, in which bins were sampled independently within each chromosome according to the number of focal bins on that chromosome and then summed across chromosomes 10^6^ times. This stratification preserves chromosome-specific differences in length and SNP density.

To directly compare the strength of telomeric enrichment between the two clades, we summarized for each clade the proportion of selected SNPs located in terminal bins relative to all SNPs included in the non-overlapping bins across the entire chromosome. Differences between clades were evaluated using a chromosome-stratified permutation framework that preserved the observed number of selected SNPs on each chromosome. In each permutation, two bins were randomly sampled with replacement from each chromosome to represent pseudo-telomeric regions, and the corresponding proportion was recalculated for both clades. The null distribution of the between-clade difference was obtained from 10^6^ permutations, and right-tailed *P*-values were calculated as the probability that the permuted difference was equal to or greater than the observed difference.

To evaluate overall spatial tendencies of selected SNPs independent of predefined focal regions, we assessed how close each selected SNP was localized toward the center of its chromosome relative to chromosome length. For each clade, a SNP-count–weighted average was calculated as the sum of (relative distance × SNP count) divided by the total SNP count, providing a measure of whether selected loci were preferentially located toward chromosome ends (0%) or centers (100%). Between-clade differences were tested using the same chromosome-stratified permutation approach described above in R, which preserved chromosome-specific SNP counts and length differences. This analysis explicitly accounted for the contrasting genome architectures of the two clades, with the Atlantic genome comprising 529 Mb across four chromosomes and the Europe genome comprising 671 Mb across fifteen chromosomes^29,42^. Based on the mean chromosome lengths of approximately 132 Mb in the Atlantic clade and 45 Mb in the Europe clade, the observed percentage shift toward chromosome centers in the Atlantic clade corresponds to an average displacement of (percentage shift × mean chromosome length), equivalent to ∼2 Mb in the Atlantic clade compared with ∼0.7 Mb in the Europe clade.

### Quantifying parallelism of the genome-wide selection response

To quantify the extent of parallel selection response between selection lines from the two clades, three complementary statistical approaches were applied. First, the Jaccard index^57^ was computed to compare the pairwise overlap of selected alleles between replicate lines relative to null simulations of our data. Second, a linear model-based slope analysis^58^ was used to test whether allele frequency changes were consistent in direction and magnitude across replicates. Third, a principal component analysis (PCA)^87^ was conducted to assess the clustering of replicate lines based on genome-wide allele frequency changes across generations.

For the Jaccard index, the pairwise overlap of selected haplotype blocks (alleles) was measured between every pair of replicate lines within a clade at a given generation (e.g., 0, 7, 10, 20). A haplotype block was considered “selected” in a replicate if its frequency increase exceeded the 99.9th percentile of the neutral drift distribution. For each pair of lines, the Jaccard index was calculated as the number of shared selected haplotype blocks divided by the total number of unique haplotype blocks detected in either line. This index ranges from 0 (no common selected alleles) to 1 (identical sets of selected alleles). The mean Jaccard index across all pairwise comparisons among replicates was reported as an overall summary of parallelism at each time point.

For the linear model-based slope analysis, allele frequency trajectories were modeled separately for each allele in each replicate using a linear regression of allele frequency across generations. The regression slope represented the rate of allele frequency change over time. To account for sampling noise and stochastic variation, slope variances were estimated from neutral drift simulations and were used to standardize the observed slopes. Alleles with slopes significantly greater than the neutral expectation (*P* < 0.05) were identified within each replicate line. To evaluate whether trajectories were parallel across replicates, a Chi-square test was used to assess slope heterogeneity across replicate lines. Bonferroni-adjusted confidence intervals were also computed as a conservative alternative criterion. Alleles showing significant among-replicate heterogeneity were classified as non-parallel. The proportion of fully parallel alleles was calculated as the fraction of loci that were putatively adaptive in all replicates and not classified as non-parallel, divided by the total number of loci tested.

Finally, PCA was conducted to assess the similarity of genomic responses among replicate selection lines. For each clade, VCF files containing SNPs underlying selected haplotype blocks (alleles) were generated. PCA was performed in PLINK v1.9^88^ using the allele frequency matrix derived from these selected SNPs. Specifically, allele counts were extracted from the filtered VCFs, with each replicate line at a given generation treated as a separate observation. PLINK computes eigenvectors based on the variance–covariance matrix of genotype frequencies. The resulting principal components were visualized in R, with selection line time points plotted in the plane of the first two principal components.

### Simulation of the role of positive epistasis on empirical parallelism

Forward-time Wright–Fisher simulations implemented in SLiM v3.7^65^ were used to test whether positive synergistic epistasis could reproduce the extent of parallelism observed in our empirical selection lines at a given generation. Simulations were designed to mimic the conditions of the E&R experiments for each clade, including the effective population size, number of selected loci, selection strength, and chromosome number.

For each clade, 10 and 7 replicate populations were simulated, analogous to the number of replicate selection lines in the Europe (10 replicate lines) and Atlantic (7 replicate lines) clades. The diploid population size per simulated line was set to the estimated *N*_e_ of the experimental populations (Atlantic clade *N*_e_ = 580 and Europe clade *N*_e_ = 740). Each simulation was run for the same number of generations as the experiment (10 generations for the Europe clade, 20 generations for the Atlantic clade), and simulated allele frequencies were sampled at intermediate points (Generations 7, 10, and 20 where applicable).

In the simulations, a defined set of beneficial loci corresponding to the number of haplotype blocks (alleles) under selection was introduced in each clade. The number of selected loci was set to match the empirically observed count of selected haplotype blocks (alleles), 261 loci in the Europe clade (across 15 chromosomes) and 66 loci in the Atlantic clade (across 4 chromosomes). These loci were positioned on a simplified genomic map with the same chromosome number and lengths as the real genomes, thereby preserving the distribution of adaptive loci per chromosome observed in each clade. Each selected locus was given an allele frequency equal to the empirical starting frequency of the corresponding empirical haplotype in Generation 0 (drawn from our pool-seq data). Loci were otherwise assumed to be neutral polymorphisms initially. Recombination was modeled by assigning a recombination rate of approximately two crossovers per chromosome per meiosis event. Thus, in the Atlantic clade simulations, 4 chromosomes each had two crossovers per meiosis on average. Likewise, the 15 chromosomes in the Europe clade simulations also had two crossovers each, yielding a greater total recombination rate per genome per generation in the Europe clade.

To explore the effect of positive synergistic epistatic interactions between selected loci on parallelism, a variable epistasis parameter (α) was introduced into the fitness model in our simulations. The framework of epistasis adapted by Stern et al.^4^ from Keightley and Otto^64^ was followed. In this model, an individual’s fitness 𝑤_i_ with respect to the selected loci is given by:

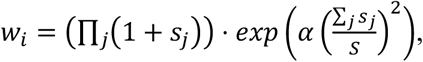

where 𝑠_j_ is the fitness advantage of the beneficial allele at locus *j*. The sum ∑ 𝑠_j_ and the product ∏(1 + 𝑠_j_)( are taken over all loci *j,* where individual *i* has the beneficial allele.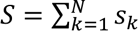 is the total of the fitness advantages across all *N* loci. The parameter α determines the strength of the epistatic effect.

Assuming that every beneficial allele has a fitness advantage of *s*, the formula for the fitness of an individual with *n* beneficial alleles simplifies to:

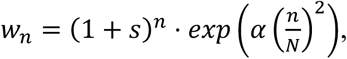

When α is set to 0, the fitness is multiplicative with respect to the number of beneficial alleles carried by an individual, and there is no epistasis^4^.

Each run was terminated after 10 generations when simulating the Europe clade and 20 generations for the Atlantic clade. Frequencies of the beneficial alleles were recorded after 7, 10, and 20 (Atlantic clade only) generations and separately for each subpopulation, by sampling 100 random individuals from each subpopulation (to mimic the experiment) and calculating the observed allele frequencies. To determine whether the change in beneficial allele frequency over a number *k* of generations within a subpopulation was significant, we calculated the 99th quantile 𝑞_k,0.99_ of the distribution of allele frequencies that one expects to observe after *k* generations if the locus were neutral (i.e., under neutrality given its initial frequency). If the frequency of an allele rose to a value exceeding 𝑞_k,0.99_ over *k* generations, the change was then regarded as significant.

For each run, an incidence matrix of significant allele frequency changes was obtained for all loci and all lines after 7, 10, and 20 (Atlantic clade only) generations. As in the experiment, the Jaccard index was calculated between all pairwise combinations of the 10 (Europe clade) or 7 (Atlantic clade) replicate lines. The final Jaccard index after 7, 10, and 20 generations was obtained by taking the mean of all Jaccard indices obtained for the given generation. This score was used to compare the amount of parallelism observed in the simulations to that observed in the experiments.

Using the SLiM setup as described above, an Approximate Bayesian computation (ABC) approach was employed to estimate the level of epistasis present during experimental evolution in the Atlantic and Europe clade selection lines. In particular, we determined the value of α that best explains the observed pattern in the two clades. The Lenormand algorithm^89^ implemented in the R package EasyABC^90^ was used to generate successive Jaccard indices obtained from SLiM runs for given proposal values 𝛼_i_, 𝑖 = 1, …, 𝑙 for the parameter α, using a uniform prior distribution on the interval [0,100]. This algorithm repeats the runs while updating the proposal values, until the Jaccard scores obtained cannot be brought closer to the target observations from the selection experiments, using a numerical stopping criterion (see Lenormand et al.^89^). The final 1,000 iterations of the algorithm were used to represent the posterior distribution of α, with the weighted mean taken as the point estimate.

### Simulations and analyses of the interaction of chromosome number and epistasis on parallelism

An additional set of forward simulations was performed to explore how differences in chromosome number (and recombination) and positive synergistic epistasis might influence the repeatability of polygenic adaptation, independently of other factors. The Europe and Atlantic clades differ fourfold in haploid chromosome count (15 versus 4), which alters the recombination rate and physical linkage relationships among loci.

Evolutionary scenarios with varying haploid chromosome counts (1, 4, 7, and 15) were simulated, covering the range from a single linkage group (one chromosome) to three different observed chromosome numbers in the *E. affinis* complex (4 for the Atlantic clade, 7 for the Gulf clade, and 15 for the Europe clade). In all other respects, simulations were set up identically to permit fair comparison. Each genome contained 160 selected loci underlying adaptation (intermediate between 66 and 261) and 480 neutral loci (threefold of the number of selected loci), yielding ∼25% of loci under selection. The 160 beneficial loci were distributed randomly across chromosomes in each scenario, with the remaining positions assigned to neutral loci. Recombination was modeled with an average of two crossovers per chromosome per meiosis; thus, total crossovers per genome increased with chromosome number (intentionally modeling greater independent assortment in high-chromosome karyotypes). All simulations used a population size of *N*_e_ = 600 and ran for 20 generations, with outputs recorded at Generations 10 and 20.

The above simulations were repeated under varying strengths of synergistic positive epistasis (α) to examine interactions between epistasis and chromosome number. Six values of α were tested (α = 0, 1.4, 4, 16, 53, and 64), spanning from purely additive (α = 0) to moderate (α = 1.4) and strong (α = 53) epistasis, corresponding to values estimated from the Atlantic and Europe clades, respectively. For each combination of chromosome number and α, 1,000 replicates were performed. To avoid bias from a particular random distribution of loci, the random genome architecture (locus-to-chromosome assignments and recombination maps) was regenerated periodically every 100 runs.

Parallelism in these simulations was evaluated using the same Jaccard index metric as mentioned in the previous sections. For each scenario, the mean Jaccard index of selected loci across the 10 simulated replicates was calculated at Generations 10 and 20, and the distribution of these indices across the 1,000 runs was compiled.

The effects of chromosome number and epistasis on the Jaccard index were analyzed using linear models in R with chromosome number as a categorical factor, α as a continuous predictor, and Jaccard index as the response variable (*Jaccard ∼ Chromosome number × α + Generation*). Significance of the effects of factors, *Chromosome number* and *α,* and *Chromosome number × α* interaction was evaluated by Type II ANOVA. Robustness of the results was confirmed by fitting the same model separately to Generations 10 and 20. Pairwise contrasts among chromosome numbers within each α level were additionally assessed using two-tailed Mann–Whitney tests.

## Acknowledgements

This work was funded by National Science Foundation grants IOS-2412790, OCE-1658517, and NSF DEB-2055356, and French National Research Agency ANR-19-MPGA-0004 (Macron’s “Make Our Planet Great Again” award) to Carol E. Lee.

## Author contributions

Conceptualization: C.E.L. and Z.D.; Methodology: Z.D., J.W., Q.L., and D.B.S.; Investigation: Z.D., J.W., and Q.L.; Resources: A.T., L.L., and S.L.; Visualization: Z.D. and C.E.L.; Funding acquisition: C.E.L.; Project administration: C.E.L.; Supervision: C.E.L. and Z.D.; Writing – original draft: Z.D. and C.E.L.; Writing – review & editing: Z.D., J.W., Q.L., A.T., L.L., S.L., D.B.S., and C.E.L.

## Data Availability

The Pool-seq data for the Atlantic clade (*Eurytemora carolleeae*) generated in this study have been deposited in the NCBI Sequence Read Archive (SRA) under BioProject PRJNA1088474. Our previously published Pool-seq data for the Europe clade (*E. affinis* proper) from BioProject PRJNA844002 (https://www.ncbi.nlm.nih.gov/bioproject/PRJNA844002) were reanalyzed in this study. Reference genomes of the Europe and Atlantic clades used in this study are available in NCBI under BioProject PRJNA1075304 (https://www.ncbi.nlm.nih.gov/bioproject/PRJNA1075304) and at figshare (https://doi.org/10.6084/m9.figshare.29104271). Allele frequency data (SNP and haplotype block) used in the statistical analysis are also available at figshare.

## Code Availability

Custom scripts for sequence data processing and selection analyses are available on Zenodo (https://doi.org/10.5281/zenodo.6615047). Scripts for location-based permutation test, linear model-based slope analysis, and forward genetic simulation are available at figshare.

