## Supplementary Figure for "Impacts of genome architecture on the repeatability of polygenic adaptation"

**Table of Contents**

**Supplementary Figure 1.** Manhattan plot of SNPs under selection in response to salinity decline on Chromosome 1 of the Europe clade genome.

**Supplementary Figure 2.** Manhattan plot of SNPs under selection in response to salinity decline on Chromosome 2 of the Europe clade genome.

**Supplementary Figure 3.** Manhattan plot of SNPs under selection in response to salinity decline on Chromosome 3 of the Europe clade genome.

**Supplementary Figure 4.** Manhattan plot of SNPs under selection in response to salinity decline on Chromosome 4 of the Europe clade genome.

**Supplementary Figure 5.** Manhattan plot of SNPs under selection in response to salinity decline on Chromosome 5 of the Europe clade genome.

**Supplementary Figure 6.** Manhattan plot of SNPs under selection in response to salinity decline on Chromosome 6 of the Europe clade genome.

**Supplementary Figure 7.** Manhattan plot of SNPs under selection in response to salinity decline on Chromosome 7 of the Europe clade genome.

**Supplementary Figure 8.** Manhattan plot of SNPs under selection in response to salinity decline on Chromosome 8 of the Europe clade genome.

**Supplementary Figure 9.** Manhattan plot of SNPs under selection in response to salinity decline on Chromosome 9 of the Europe clade genome.

**Supplementary Figure 10.** Manhattan plot of SNPs under selection in response to salinity decline on Chromosome 10 of the Europe clade genome.

**Supplementary Figure 11.** Manhattan plot of SNPs under selection in response to salinity decline on Chromosome 11 of the Europe clade genome.

**Supplementary Figure 12.** Manhattan plot of SNPs under selection in response to salinity decline on Chromosome 12 of the Europe clade genome.

**Supplementary Figure 13.** Manhattan plot of SNPs under selection in response to salinity decline on Chromosome 13 of the Europe clade genome.

**Supplementary Figure 14.** Manhattan plot of SNPs under selection in response to salinity decline on Chromosome 14 of the Europe clade genome.

**Supplementary Figure 15.** Manhattan plot of SNPs under selection in response to salinity decline on Chromosome 15 of the Europe clade genome.

**Supplementary Figure 16.** Manhattan plot of SNPs under selection in response to salinity decline on Chromosome 1 of the Atlantic clade genome.

**Supplementary Figure 17.** Manhattan plot of SNPs under selection in response to salinity decline on Chromosome 2 of the Atlantic clade genome.

**Supplementary Figure 18.** Manhattan plot of SNPs under selection in response to salinity decline on Chromosome 3 of the Atlantic clade genome.

**Supplementary Figure 19.** Manhattan plot of SNPs under selection in response to salinity decline on Chromosome 4 of the Atlantic clade genome.

**Supplementary Figure 20.** Location-based permutation test of telomeric enrichment of selected SNPs in the Europe clade (1-Mb bins).

**Supplementary Figure 21.** Location-based permutation test of telomeric enrichment of selected SNPs in the Europe clade (2-Mb bins).

**Supplementary Figure 22.** Location-based permutation test of telomeric enrichment of selected SNPs in the Atlantic clade (1-Mb bins).

**Supplementary Figure 23.** Location-based permutation test of telomeric enrichment of selected SNPs in the Atlantic clade (2-Mb bins).

**Supplementary Figure 24.** Location-based permutation test of fusion-site enrichment of selected SNPs in the Atlantic clade (1-Mb bins).

**Supplementary Figure 25.** Location-based permutation test of fusion-site enrichment of selected SNPs in the Atlantic clade (2-Mb bins).

**Supplementary Figure 26. Time-resolved allele-frequency trajectories of selected haplotype blocks in the Atlantic clade across all sampled generations.**

**Supplementary Figure 27.** Effects of epistasis (α) and chromosome number (n) on the repeatability of adaptation (Jaccard index) at Generations 10 and 20.

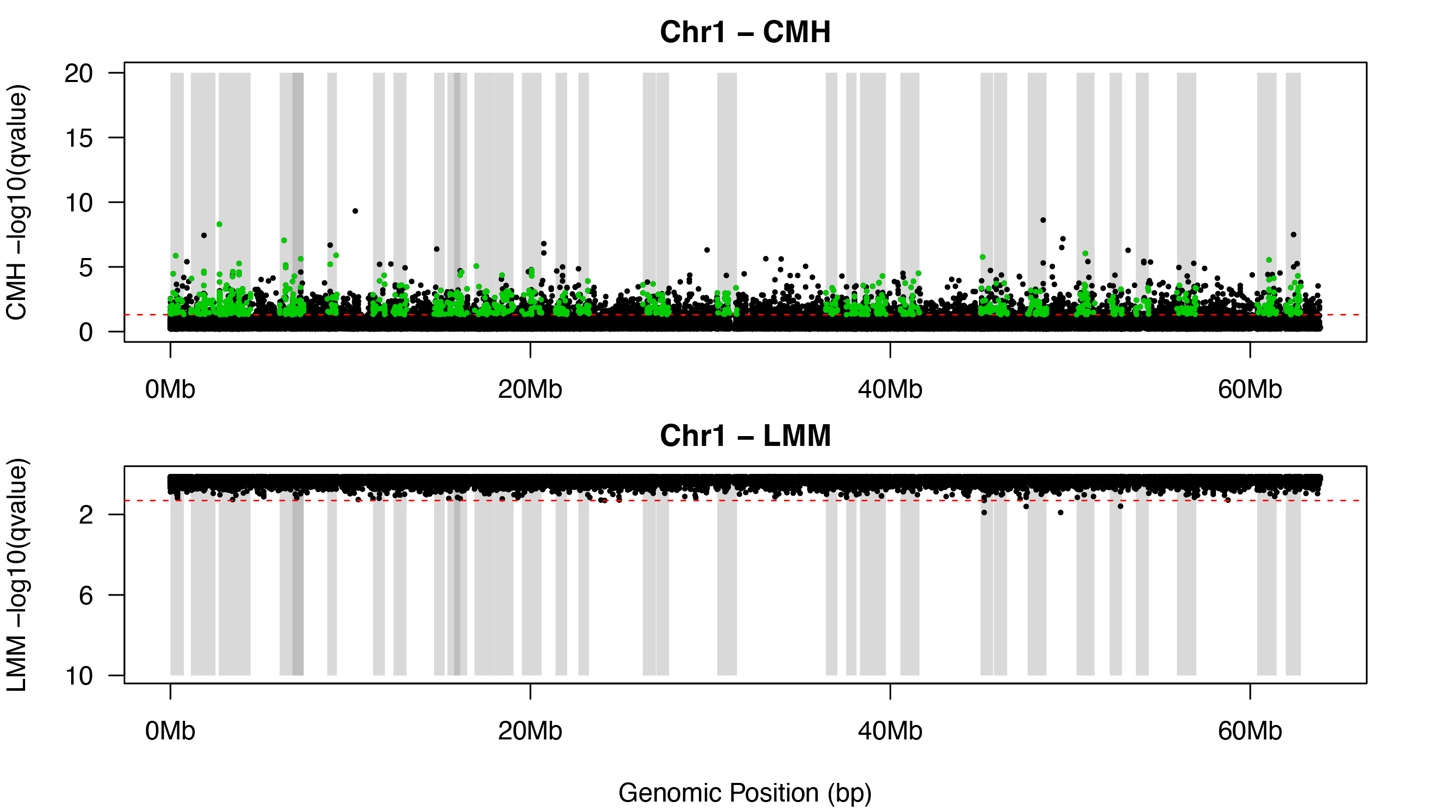

**Supplementary Figure 1.** **Manhattan plot of SNPs under selection in response to salinity decline on Chromosome 1 of the Europe clade genome.** SNPs above the dotted red line were deemed significant after correction for multiple testing (adjusted *P* < 0.05). Shaded gray bars delineate haplotype blocks identified as targets of selection on this chromosome. Top: results of the Cochran–Mantel–Haenszel (CMH) test, detecting significant allele frequency changes beyond expectations from genetic drift. Bottom: results of the linear mixed model (LMM) test, distinguishing allele frequency trajectories between selection and control lines.

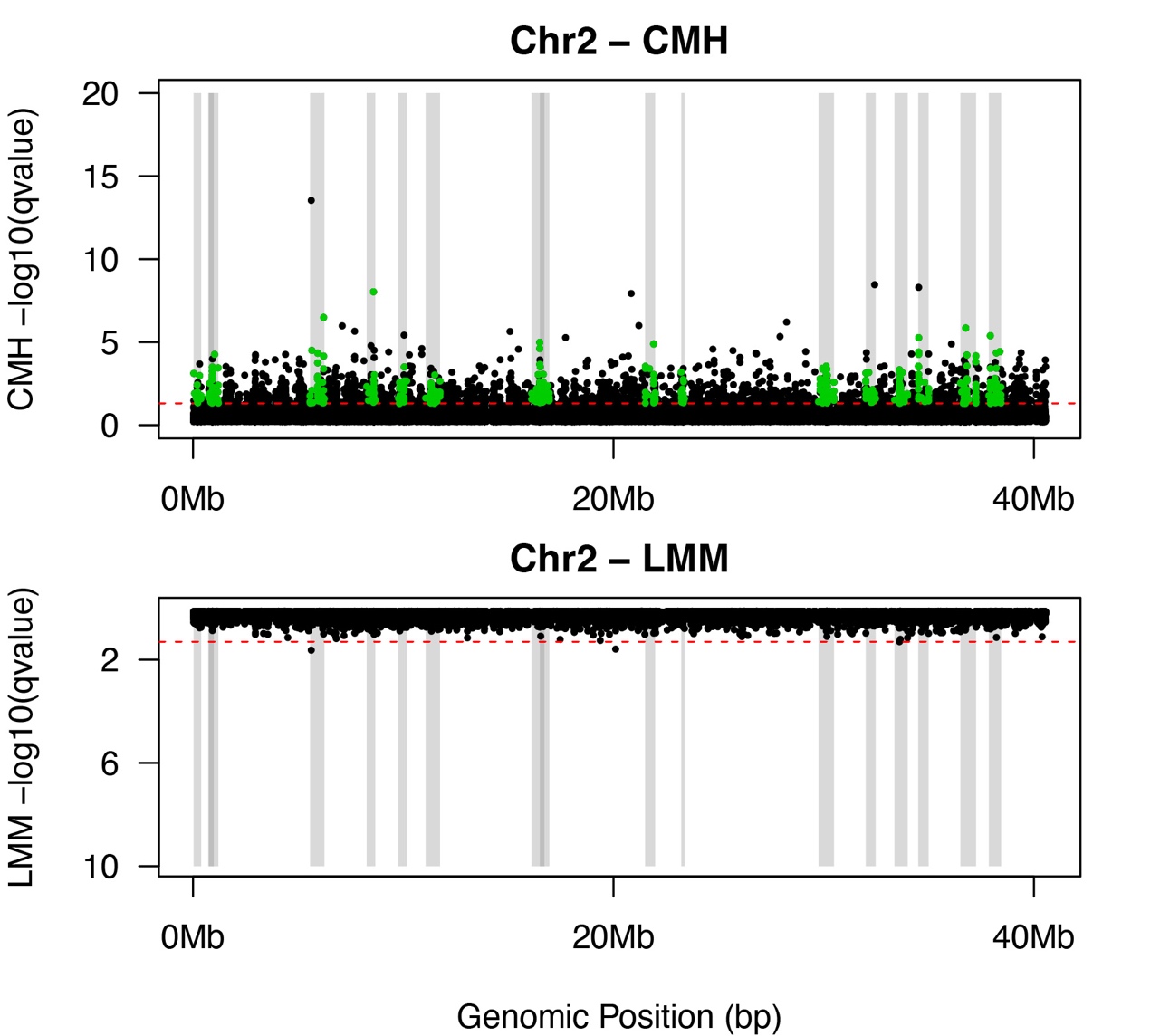

**Supplementary Figure 2.** **Manhattan plot of SNPs under selection in response to salinity decline on Chromosome 2 of the Europe clade genome.** SNPs above the dotted red line were deemed significant after correction for multiple testing (adjusted *P* < 0.05). Shaded gray bars delineate haplotype blocks identified as targets of selection on this chromosome. Top: results of the Cochran–Mantel–Haenszel (CMH) test, detecting significant allele frequency changes beyond expectations from genetic drift. Bottom: results of the linear mixed model (LMM) test, distinguishing allele frequency trajectories between selection and control lines.

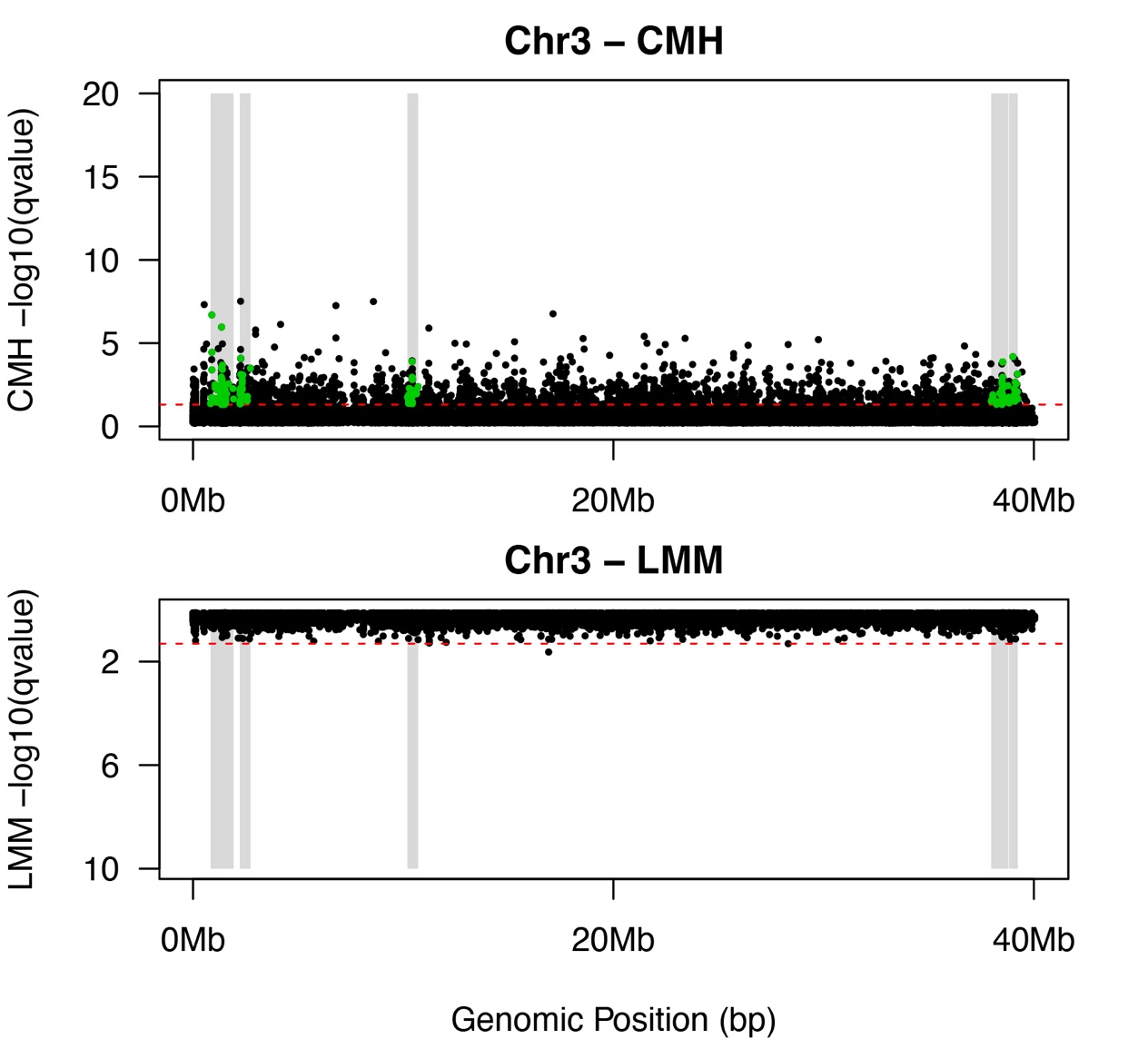

**Supplementary Figure 3.** **Manhattan plot of SNPs under selection in response to salinity decline on Chromosome 3 of the Europe clade genome.** SNPs above the dotted red line were deemed significant after correction for multiple testing (adjusted *P* < 0.05). Shaded gray bars delineate haplotype blocks identified as targets of selection on this chromosome. Top: results of the Cochran–Mantel–Haenszel (CMH) test, detecting significant allele frequency changes beyond expectations from genetic drift. Bottom: results of the linear mixed model (LMM) test, distinguishing allele frequency trajectories between selection and control lines.

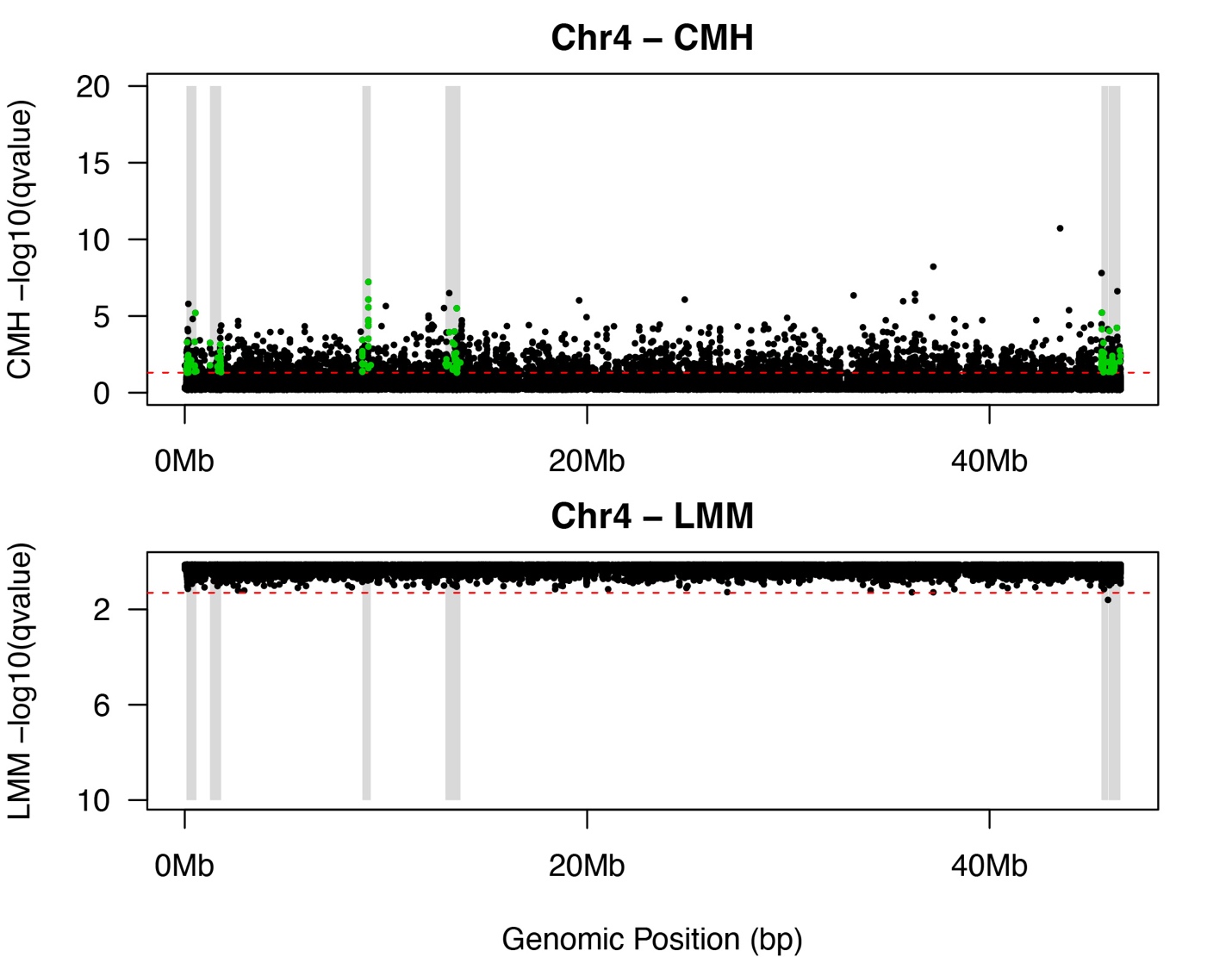

**Supplementary Figure 4.** **Manhattan plot of SNPs under selection in response to salinity decline on Chromosome 4 of the Europe clade genome.** SNPs above the dotted red line were deemed significant after correction for multiple testing (adjusted *P* < 0.05). Shaded gray bars delineate haplotype blocks identified as targets of selection on this chromosome. Top: results of the Cochran–Mantel–Haenszel (CMH) test, detecting significant allele frequency changes beyond expectations from genetic drift. Bottom: results of the linear mixed model (LMM) test, distinguishing allele frequency trajectories between selection and control lines.

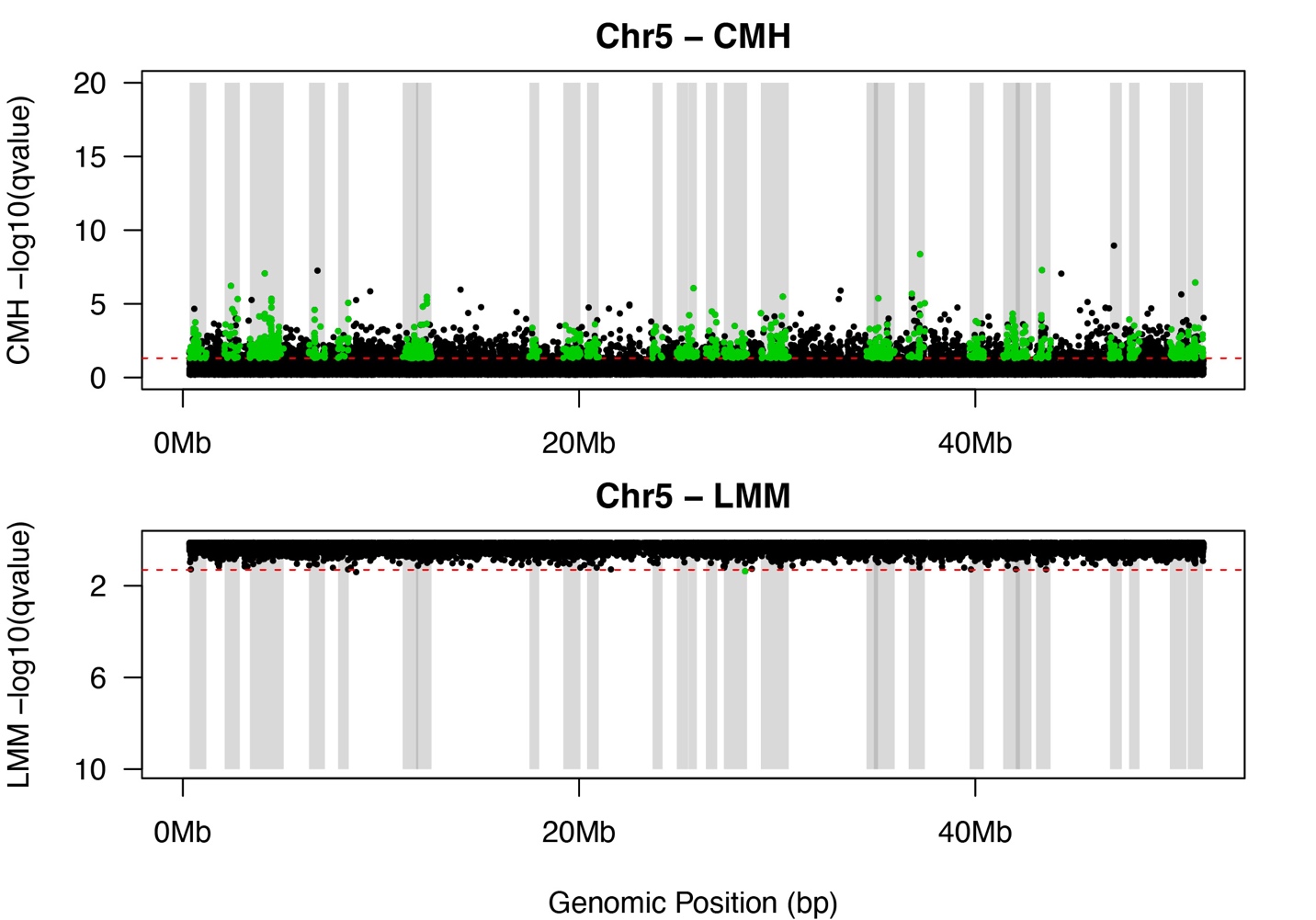

**Supplementary Figure 5.** **Manhattan plot of SNPs under selection in response to salinity decline on Chromosome 5 of the Europe clade genome.** SNPs above the dotted red line were deemed significant after correction for multiple testing (adjusted *P* < 0.05). Shaded gray bars delineate haplotype blocks identified as targets of selection on this chromosome. Top: results of the Cochran–Mantel–Haenszel (CMH) test, detecting significant allele frequency changes beyond expectations from genetic drift. Bottom: results of the linear mixed model (LMM) test, distinguishing allele frequency trajectories between selection and control lines.

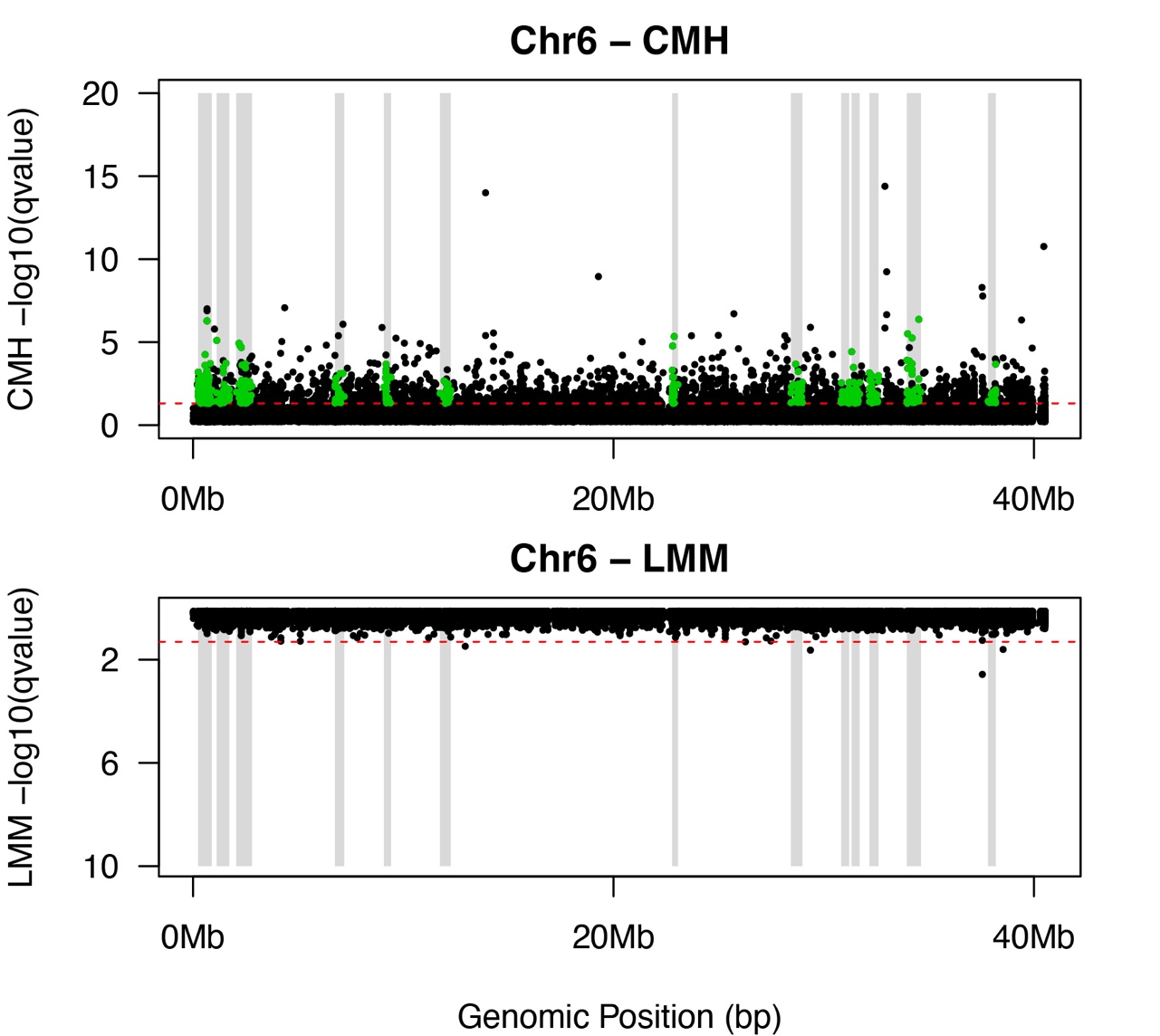

**Supplementary Figure 6.** **Manhattan plot of SNPs under selection in response to salinity decline on Chromosome 6 of the Europe clade genome.** SNPs above the dotted red line were deemed significant after correction for multiple testing (adjusted *P* < 0.05). Shaded gray bars delineate haplotype blocks identified as targets of selection on this chromosome. Top: results of the Cochran–Mantel–Haenszel (CMH) test, detecting significant allele frequency changes beyond expectations from genetic drift. Bottom: results of the linear mixed model (LMM) test, distinguishing allele frequency trajectories between selection and control lines.

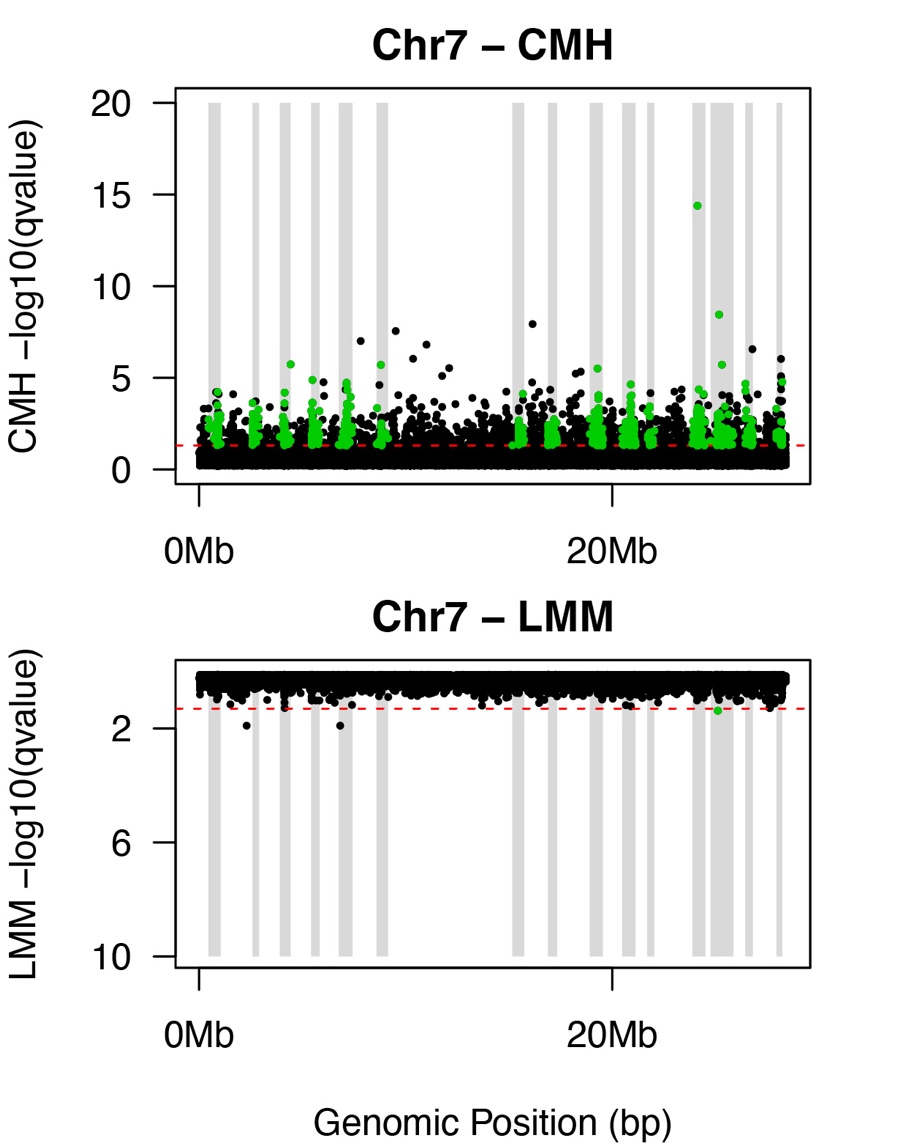

**Supplementary Figure 7.** **Manhattan plot of SNPs under selection in response to salinity decline on Chromosome 7 of the Europe clade genome.** SNPs above the dotted red line were deemed significant after correction for multiple testing (adjusted *P* < 0.05). Shaded gray bars delineate haplotype blocks identified as targets of selection on this chromosome. Top: results of the Cochran–Mantel–Haenszel (CMH) test, detecting significant allele frequency changes beyond expectations from genetic drift. Bottom: results of the linear mixed model (LMM) test, distinguishing allele frequency trajectories between selection and control lines.

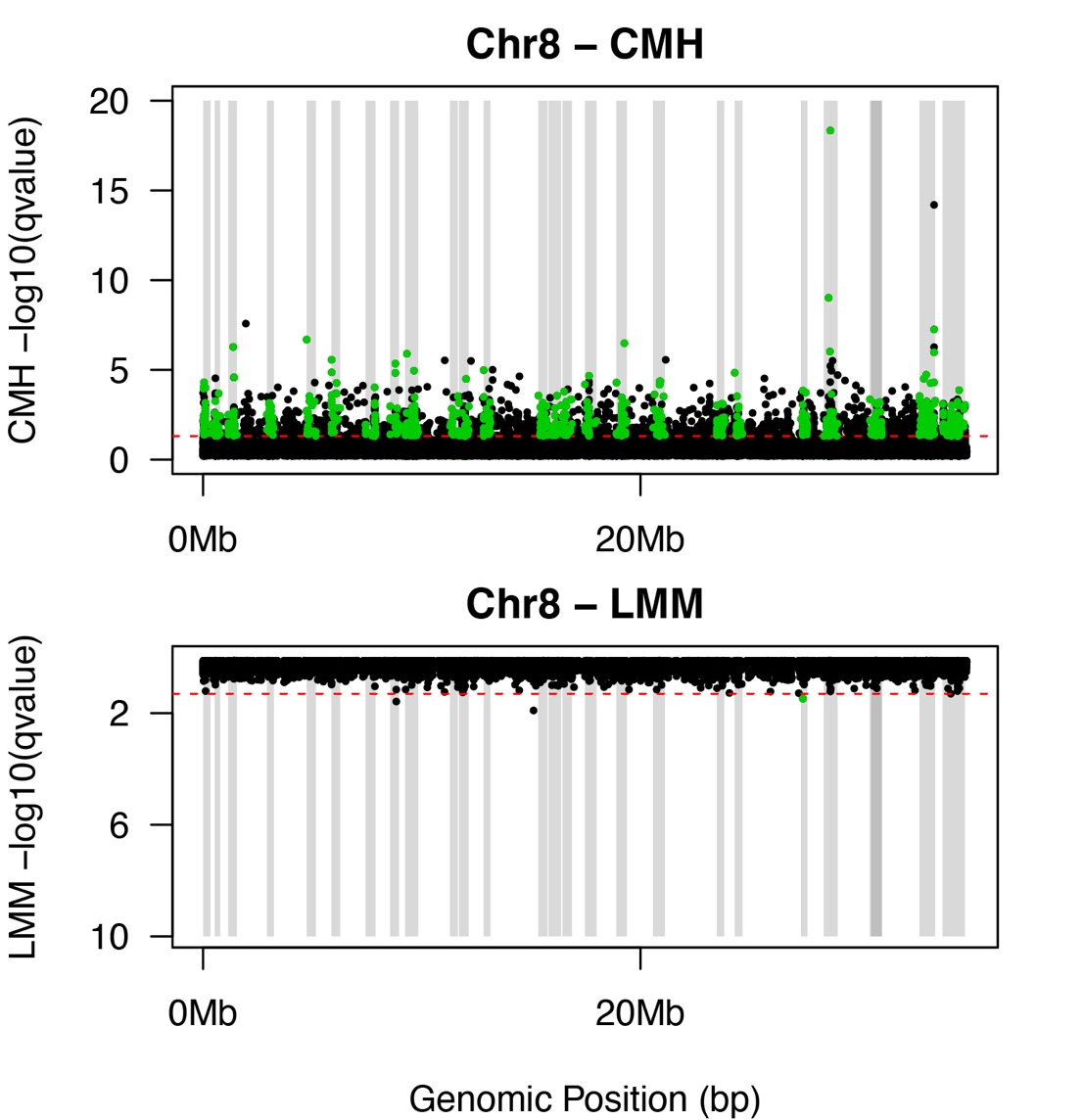

**Supplementary Figure 8.** **Manhattan plot of SNPs under selection in response to salinity decline on Chromosome 8 of the Europe clade genome.** SNPs above the dotted red line were deemed significant after correction for multiple testing (adjusted *P* < 0.05). Shaded gray bars delineate haplotype blocks identified as targets of selection on this chromosome. Top: results of the Cochran–Mantel–Haenszel (CMH) test, detecting significant allele frequency changes beyond expectations from genetic drift. Bottom: results of the linear mixed model (LMM) test, distinguishing allele frequency trajectories between selection and control lines.

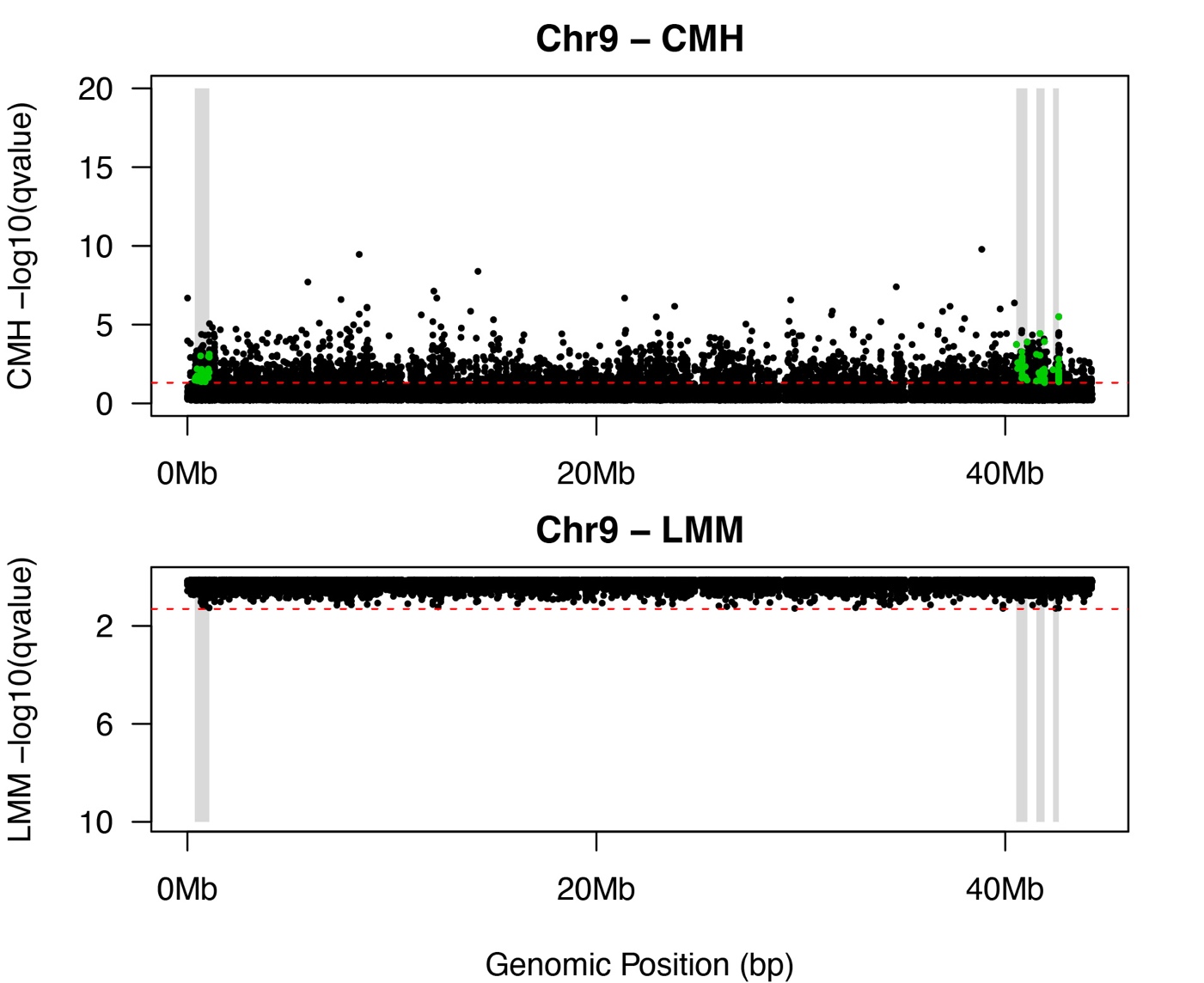

**Supplementary Figure 9.** **Manhattan plot of SNPs under selection in response to salinity decline on Chromosome 9 of the Europe clade genome.** SNPs above the dotted red line were deemed significant after correction for multiple testing (adjusted *P* < 0.05). Shaded gray bars delineate haplotype blocks identified as targets of selection on this chromosome. Top: results of the Cochran–Mantel–Haenszel (CMH) test, detecting significant allele frequency changes beyond expectations from genetic drift. Bottom: results of the linear mixed model (LMM) test, distinguishing allele frequency trajectories between selection and control lines.

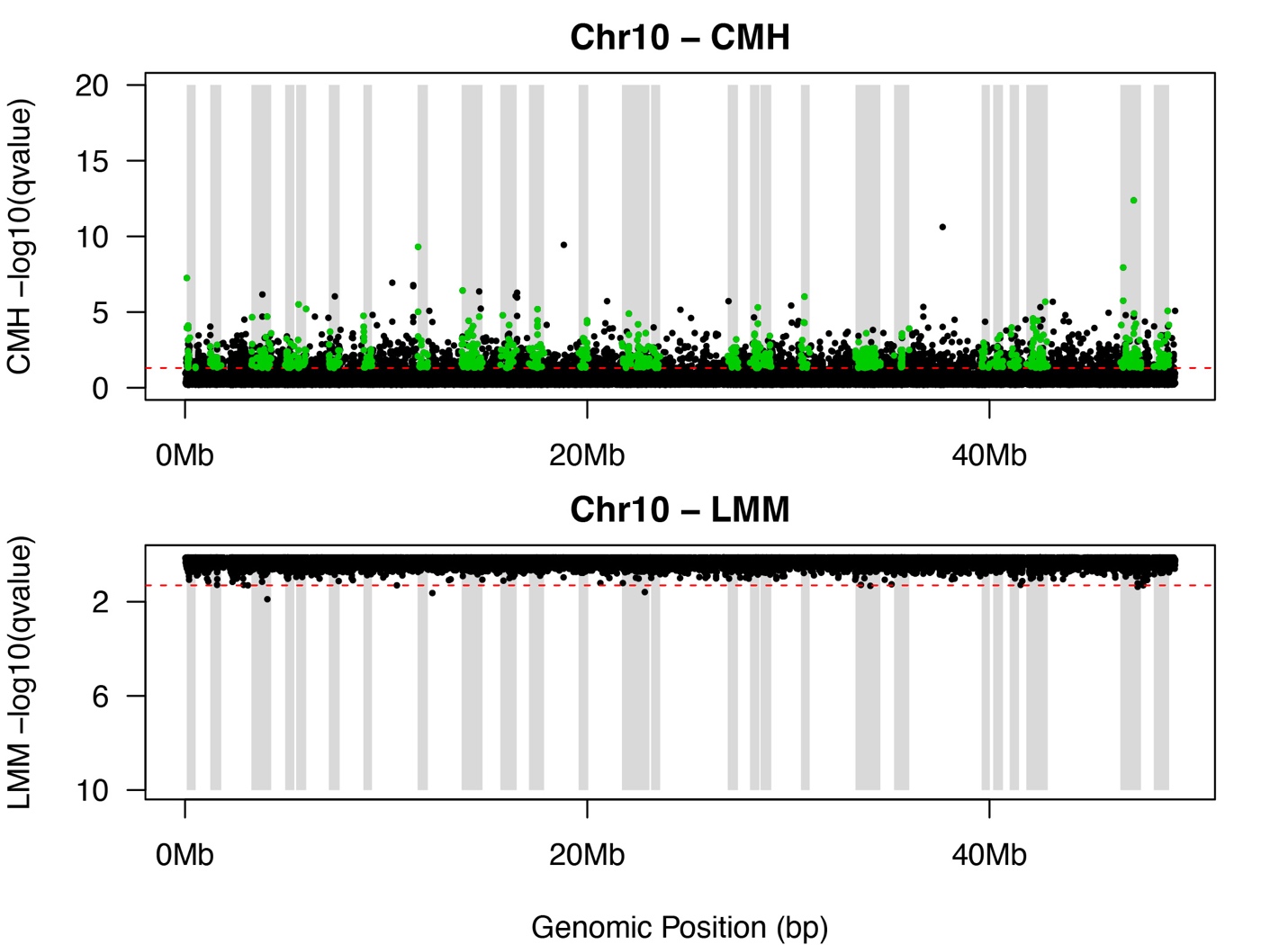

**Supplementary Figure 10.** **Manhattan plot of SNPs under selection in response to salinity decline on Chromosome 10 of the Europe clade genome.** SNPs above the dotted red line were deemed significant after correction for multiple testing (adjusted *P* < 0.05). Shaded gray bars delineate haplotype blocks identified as targets of selection on this chromosome. Top: results of the Cochran–Mantel–Haenszel (CMH) test, detecting significant allele frequency changes beyond expectations from genetic drift. Bottom: results of the linear mixed model (LMM) test, distinguishing allele frequency trajectories between selection and control lines.

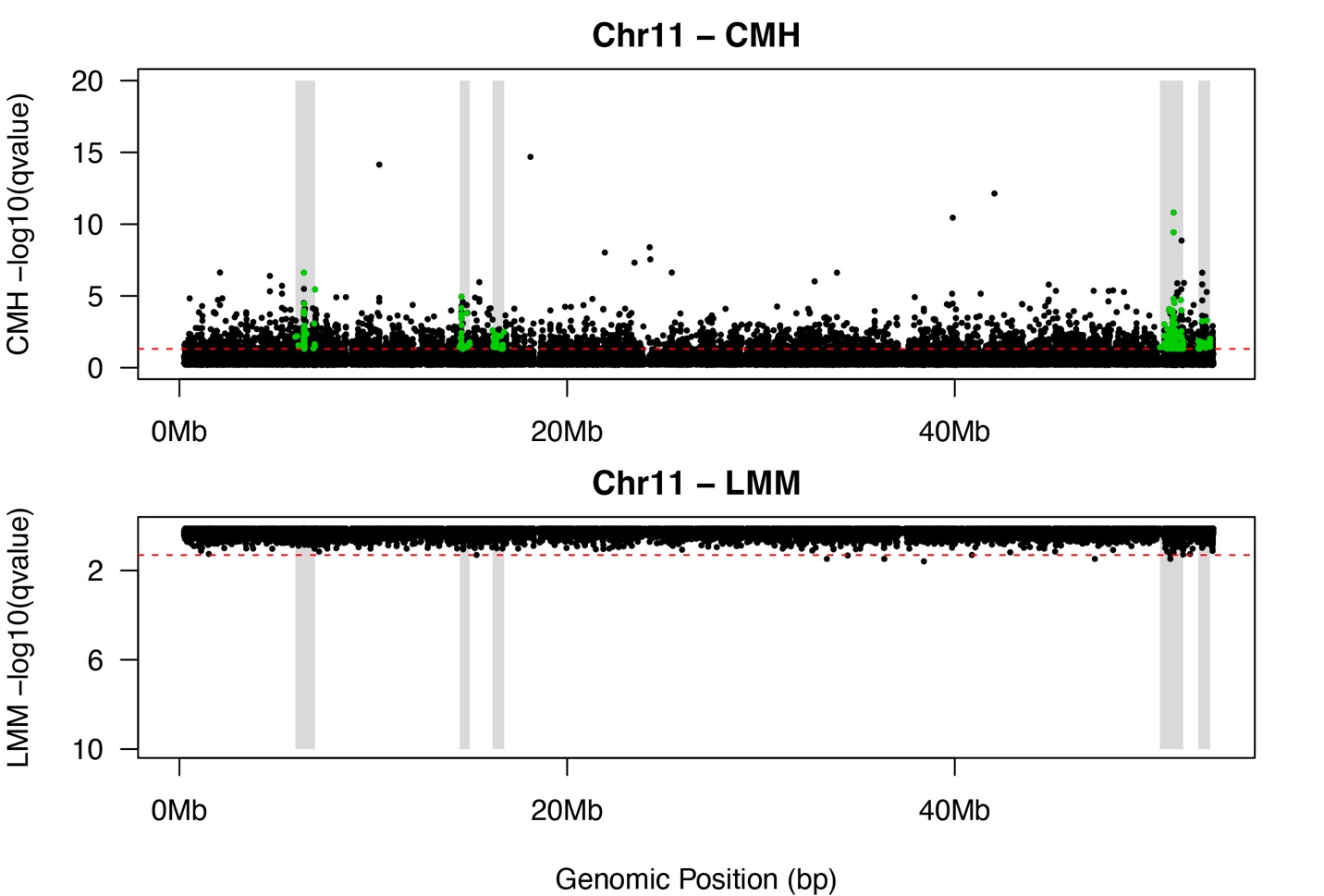

**Supplementary Figure 11.** **Manhattan plot of SNPs under selection in response to salinity decline on Chromosome 11 of the Europe clade genome.** SNPs above the dotted red line were deemed significant after correction for multiple testing (adjusted *P* < 0.05). Shaded gray bars delineate haplotype blocks identified as targets of selection on this chromosome. Top: results of the Cochran–Mantel–Haenszel (CMH) test, detecting significant allele frequency changes beyond expectations from genetic drift. Bottom: results of the linear mixed model (LMM) test, distinguishing allele frequency trajectories between selection and control lines.

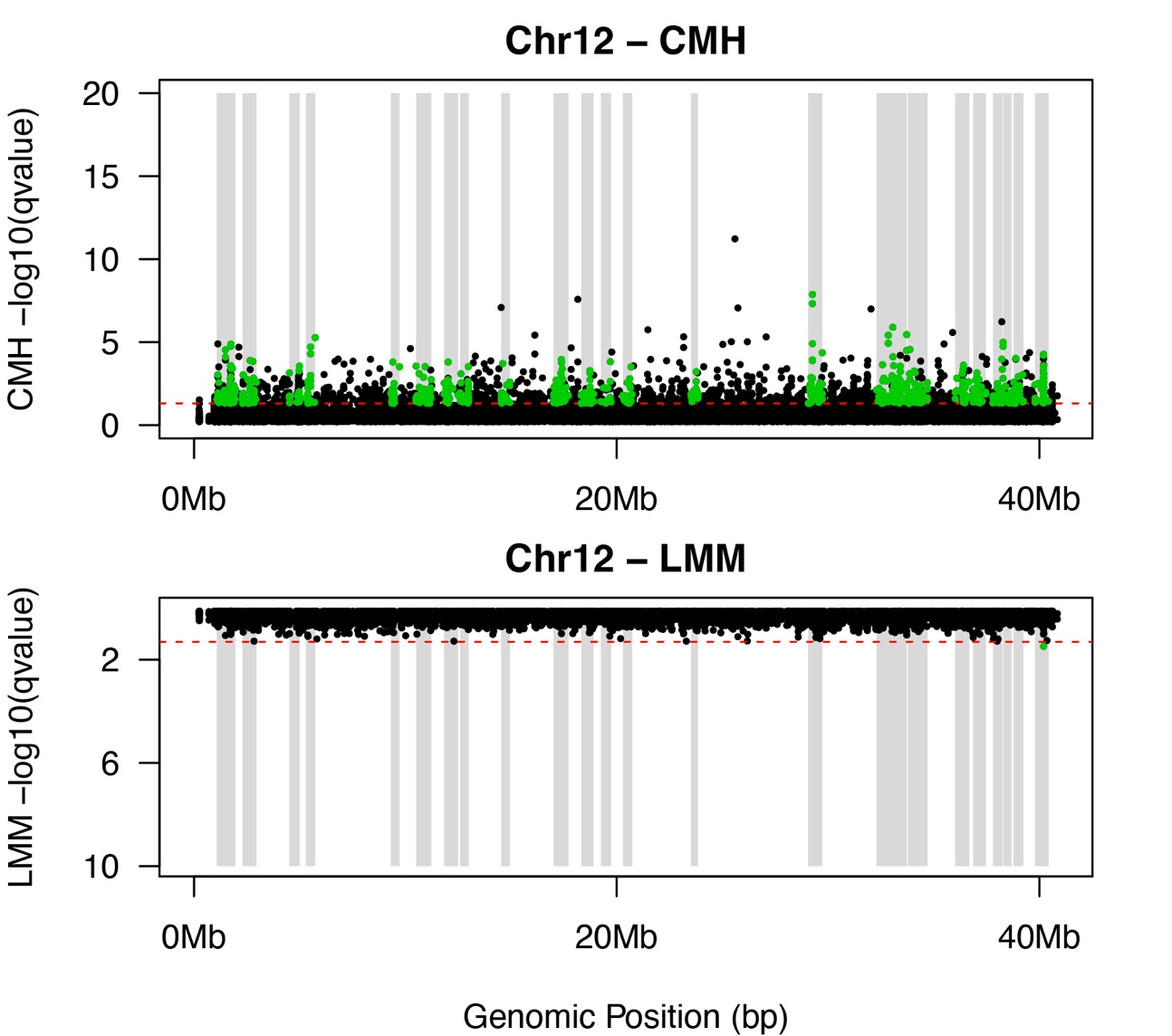

**Supplementary Figure 12.** **Manhattan plot of SNPs under selection in response to salinity decline on Chromosome 12 of the Europe clade genome.** SNPs above the dotted red line were deemed significant after correction for multiple testing (adjusted *P* < 0.05). Shaded gray bars delineate haplotype blocks identified as targets of selection on this chromosome. Top: results of the Cochran–Mantel–Haenszel (CMH) test, detecting significant allele frequency changes beyond expectations from genetic drift. Bottom: results of the linear mixed model (LMM) test, distinguishing allele frequency trajectories between selection and control lines.

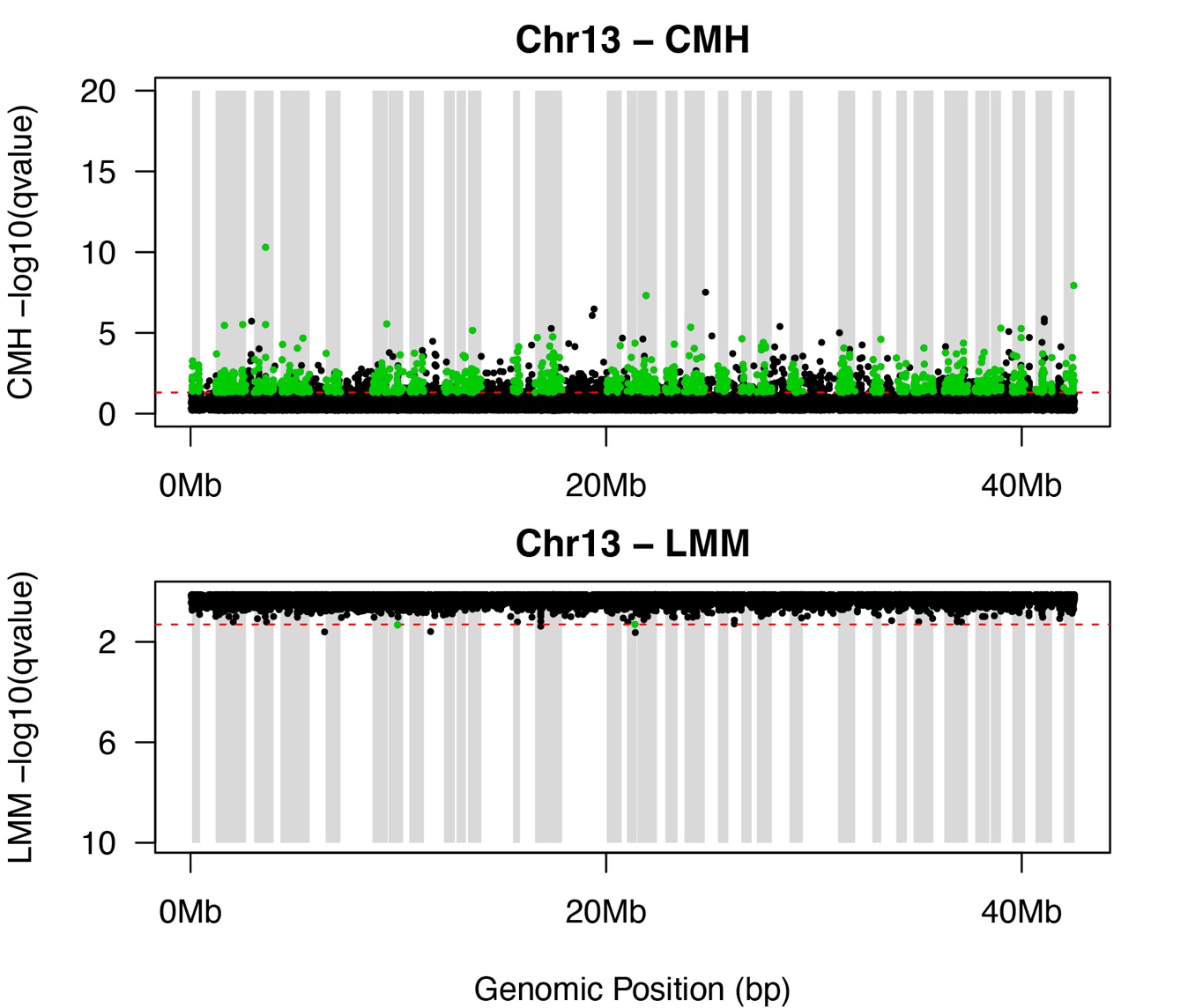

**Supplementary Figure 13.** **Manhattan plot of SNPs under selection in response to salinity decline on Chromosome 13 of the Europe clade genome.** SNPs above the dotted red line were deemed significant after correction for multiple testing (adjusted *P* < 0.05). Shaded gray bars delineate haplotype blocks identified as targets of selection on this chromosome. Top: results of the Cochran–Mantel–Haenszel (CMH) test, detecting significant allele frequency changes beyond expectations from genetic drift. Bottom: results of the linear mixed model (LMM) test, distinguishing allele frequency trajectories between selection and control lines.

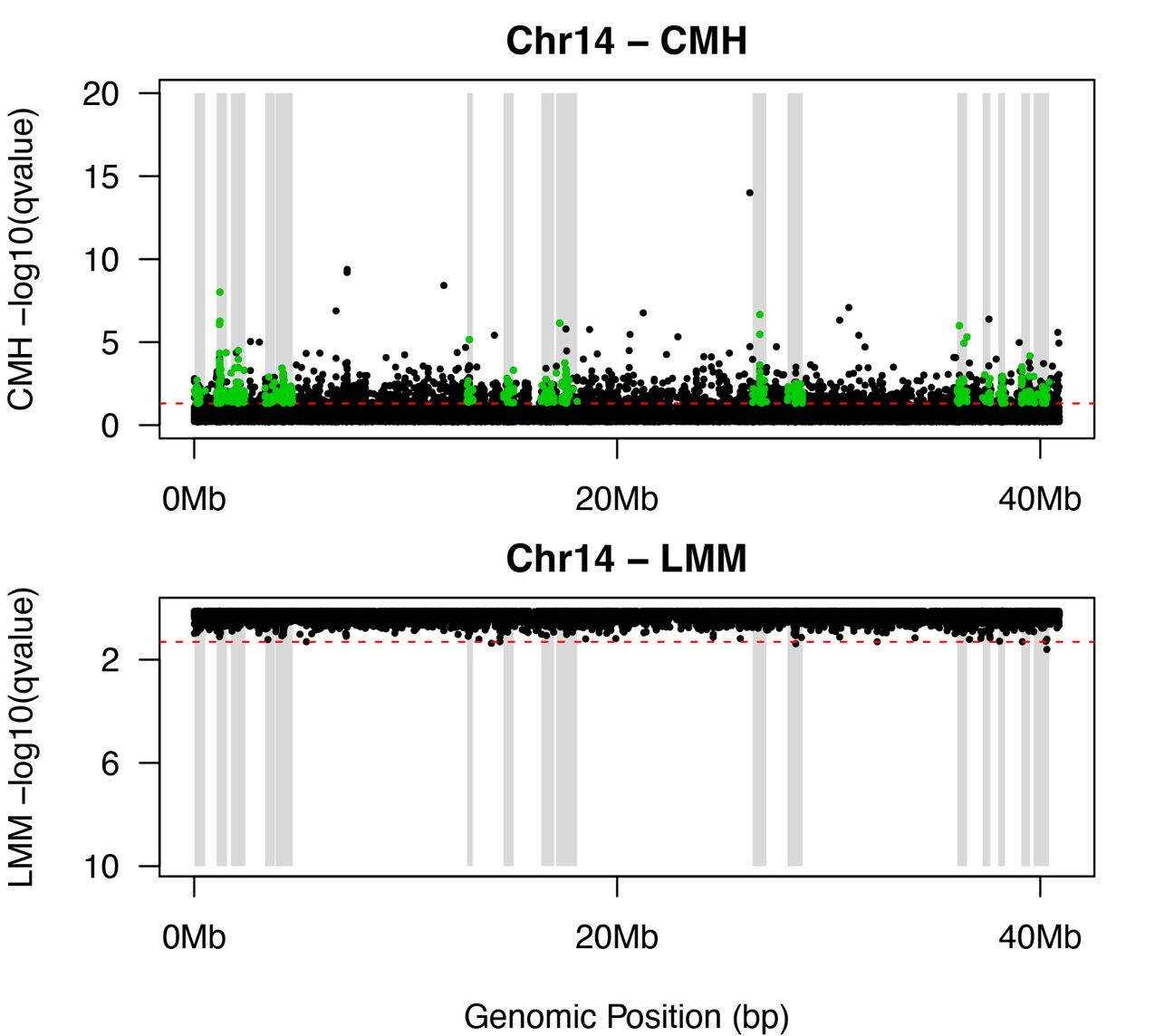

**Supplementary Figure 14.** **Manhattan plot of SNPs under selection in response to salinity decline on Chromosome 14 of the Europe clade genome.** SNPs above the dotted red line were deemed significant after correction for multiple testing (adjusted *P* < 0.05). Shaded gray bars delineate haplotype blocks identified as targets of selection on this chromosome. Top: results of the Cochran–Mantel–Haenszel (CMH) test, detecting significant allele frequency changes beyond expectations from genetic drift. Bottom: results of the linear mixed model (LMM) test, distinguishing allele frequency trajectories between selection and control lines.

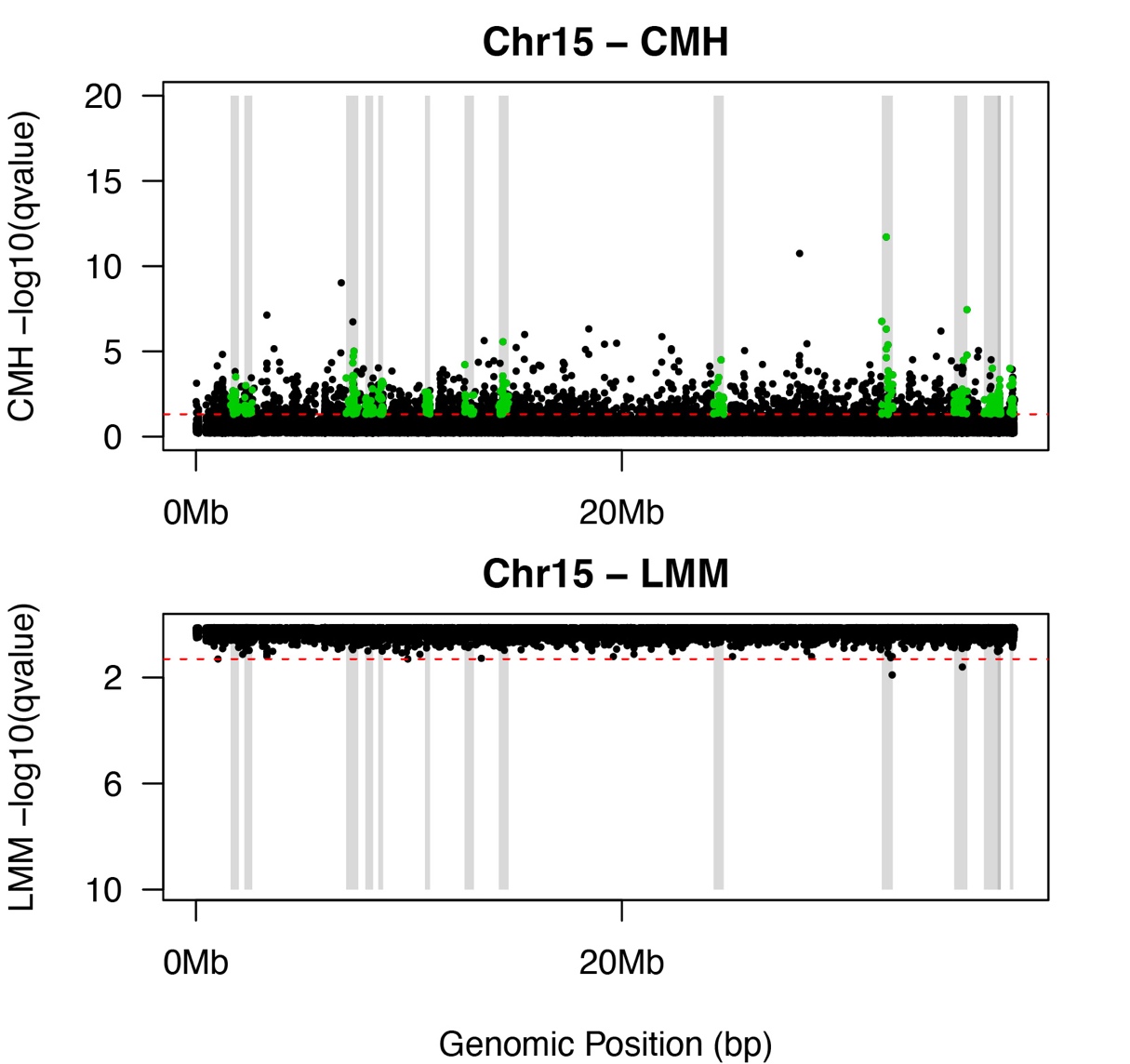

**Supplementary Figure 15.** **Manhattan plot of SNPs under selection in response to salinity decline on Chromosome 15 of the Europe clade genome.** SNPs above the dotted red line were deemed significant after correction for multiple testing (adjusted *P* < 0.05). Shaded gray bars delineate haplotype blocks identified as targets of selection on this chromosome. Top: results of the Cochran–Mantel–Haenszel (CMH) test, detecting significant allele frequency changes beyond expectations from genetic drift. Bottom: results of the linear mixed model (LMM) test, distinguishing allele frequency trajectories between selection and control lines.

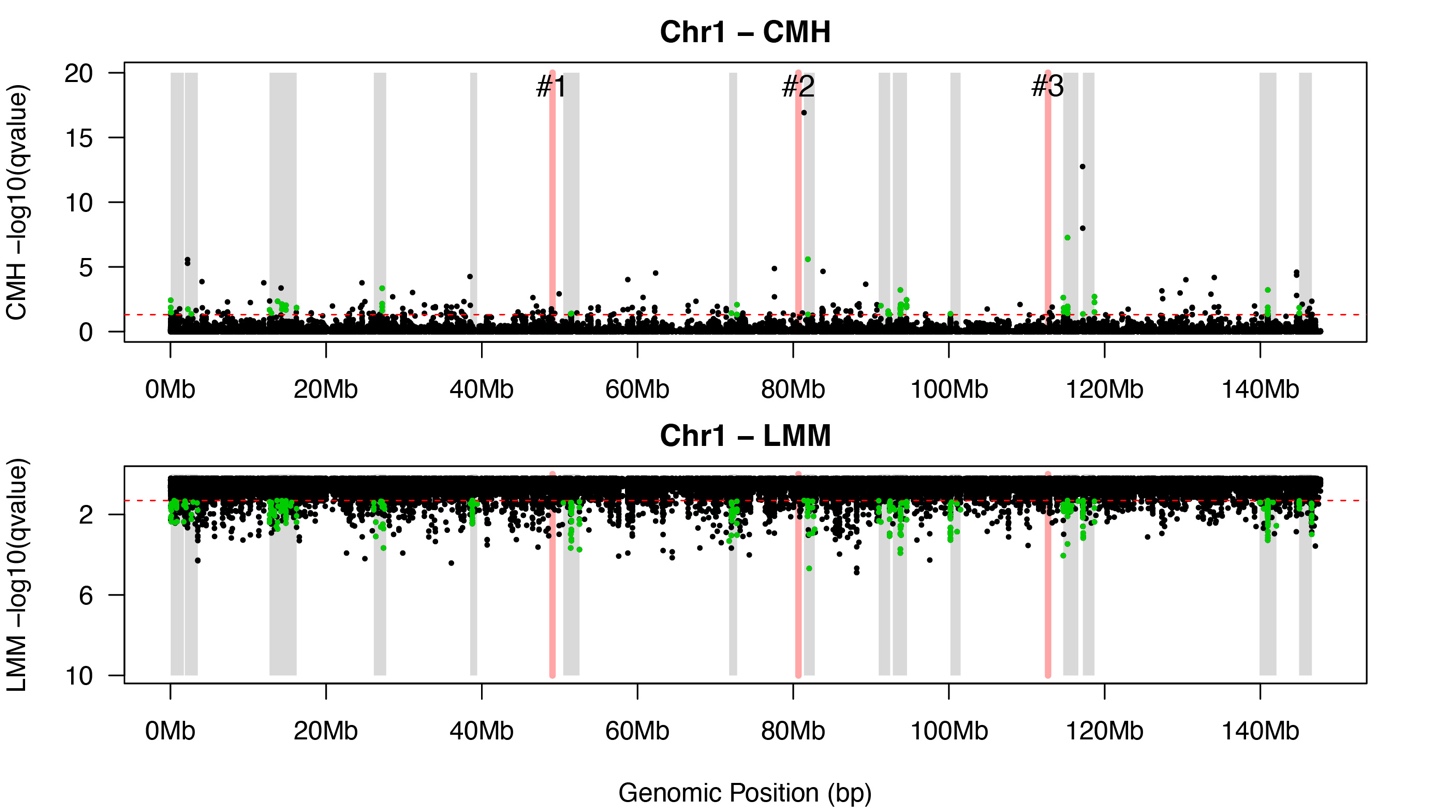

**Supplementary Figure 16.** **Manhattan plot of SNPs under selection in response to salinity decline on Chromosome 1 of the Atlantic clade genome.** SNPs above the dotted red line were deemed significant after correction for multiple testing (adjusted *P* < 0.05). Shaded gray bars delineate haplotype blocks identified as targets of selection on this chromosome. Shaded red bars delineate positions of chromosomal fusion sites. Top: results of the Cochran–Mantel–Haenszel (CMH) test, detecting significant allele frequency changes beyond expectations from genetic drift. Bottom: results of the linear mixed model (LMM) test, distinguishing allele frequency trajectories between selection and control lines.

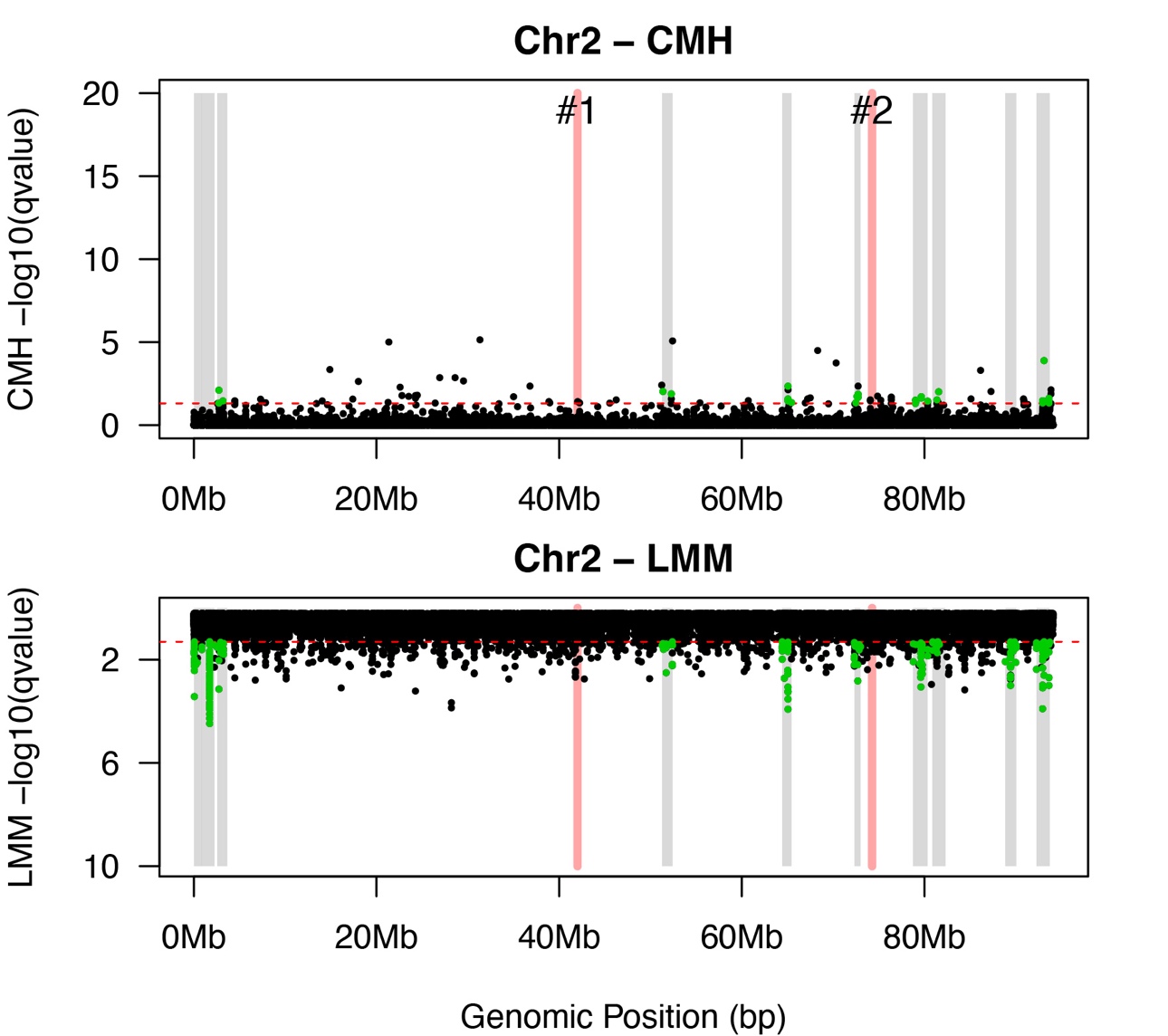

**Supplementary Figure 17.** **Manhattan plot of SNPs under selection in response to salinity decline on Chromosome 2 of the Atlantic clade genome.** SNPs above the dotted red line were deemed significant after correction for multiple testing (adjusted *P* < 0.05). Shaded gray bars delineate haplotype blocks identified as targets of selection on this chromosome. Shaded red bars delineate positions of chromosomal fusion sites. Top: results of the Cochran–Mantel–Haenszel (CMH) test, detecting significant allele frequency changes beyond expectations from genetic drift. Bottom: results of the linear mixed model (LMM) test, distinguishing allele frequency trajectories between selection and control lines.

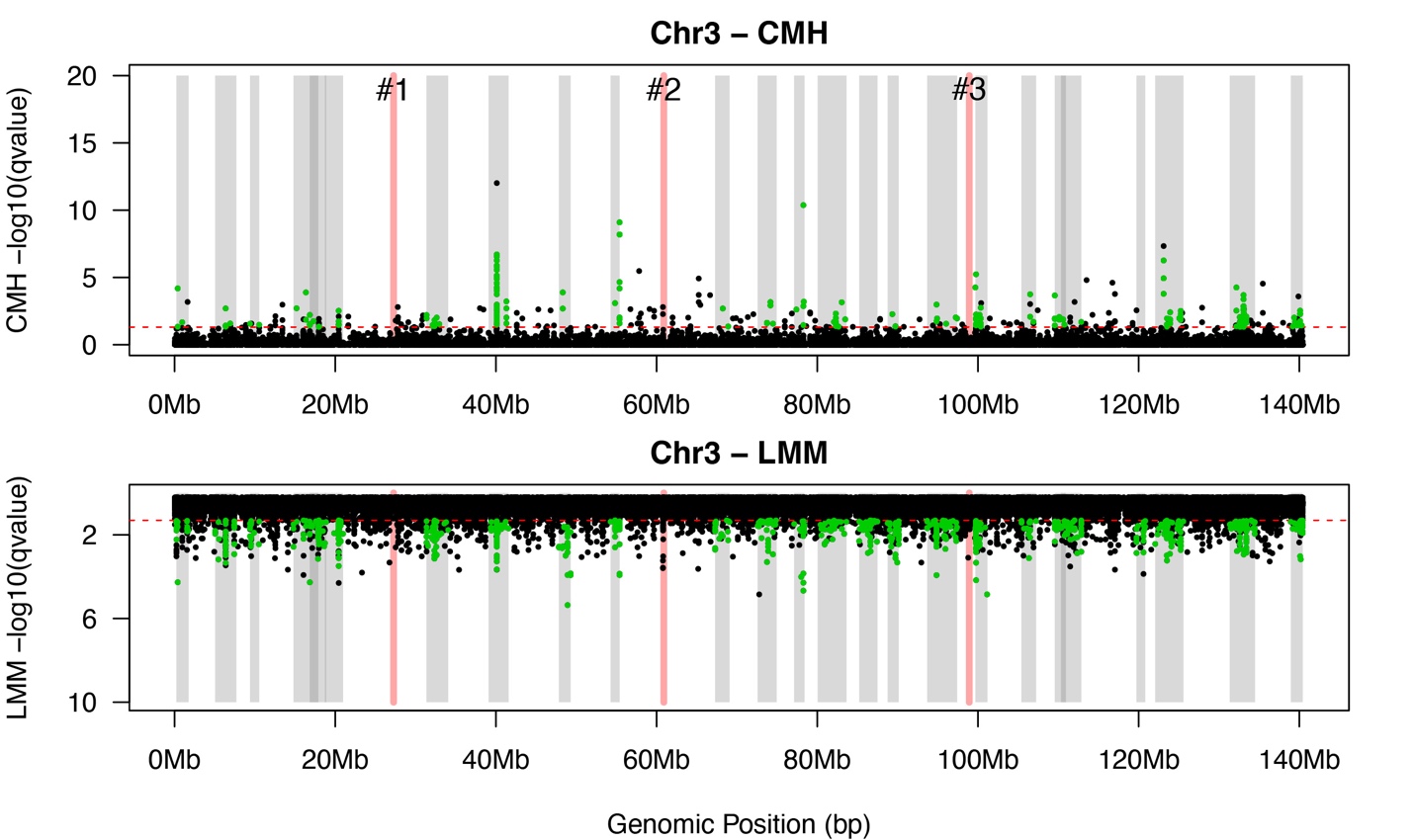

**Supplementary Figure 18.** **Manhattan plot of SNPs under selection in response to salinity decline on Chromosome 3 of the Atlantic clade genome.** SNPs above the dotted red line were deemed significant after correction for multiple testing (adjusted *P* < 0.05). Shaded gray bars delineate haplotype blocks identified as targets of selection on this chromosome. Shaded red bars delineate positions of chromosomal fusion sites. Top: results of the Cochran–Mantel–Haenszel (CMH) test, detecting significant allele frequency changes beyond expectations from genetic drift. Bottom: results of the linear mixed model (LMM) test, distinguishing allele frequency trajectories between selection and control lines.

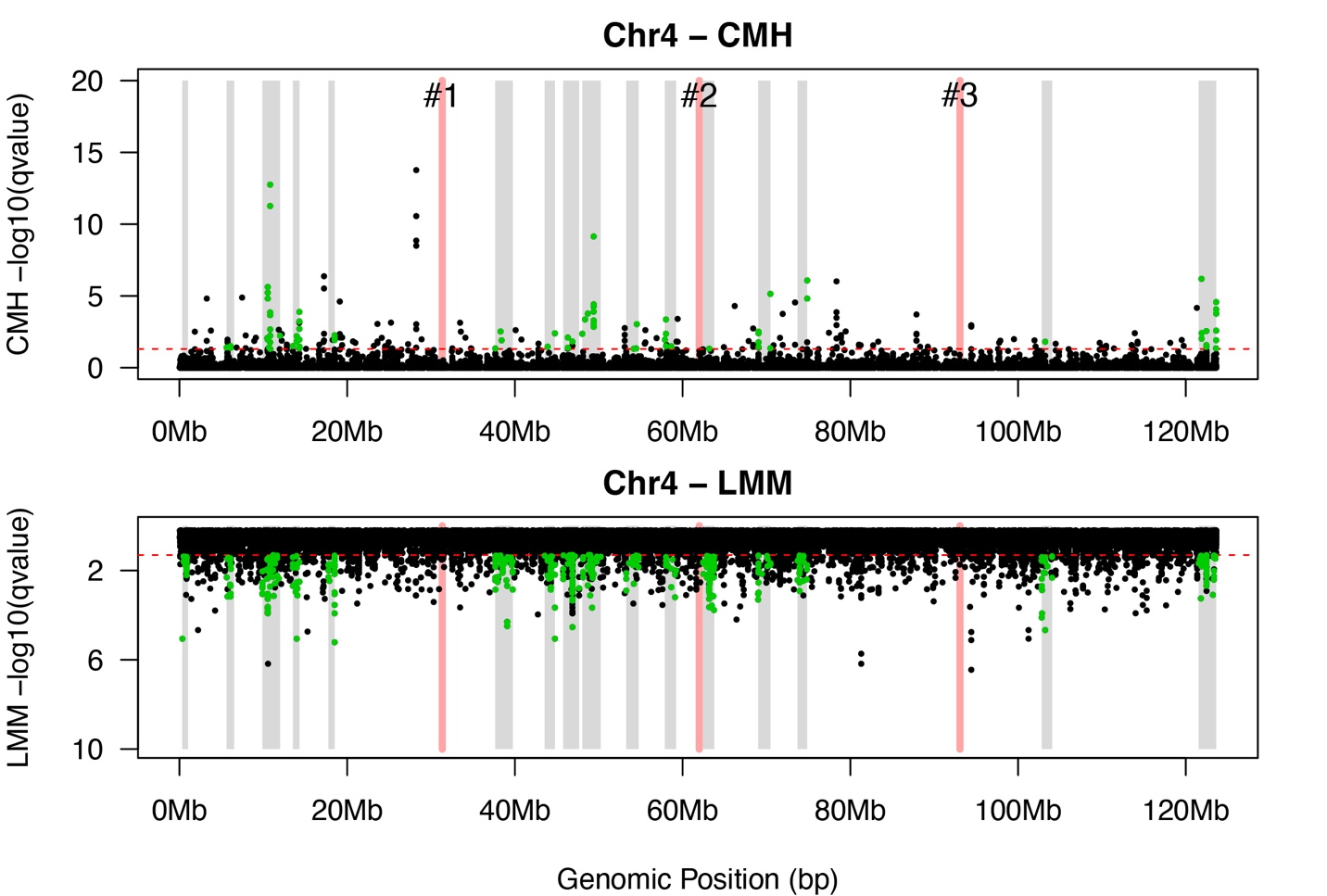

**Supplementary Figure 19.** **Manhattan plot of SNPs under selection in response to salinity decline on Chromosome 4 of the Atlantic clade genome.** SNPs above the dotted red line were deemed significant after correction for multiple testing (adjusted *P* < 0.05). Shaded gray bars delineate haplotype blocks identified as targets of selection on this chromosome. Shaded red bars delineate positions of chromosomal fusion sites. Top: results of the Cochran–Mantel–Haenszel (CMH) test, detecting significant allele frequency changes beyond expectations from genetic drift. Bottom: results of the linear mixed model (LMM) test, distinguishing allele frequency trajectories between selection and control lines.

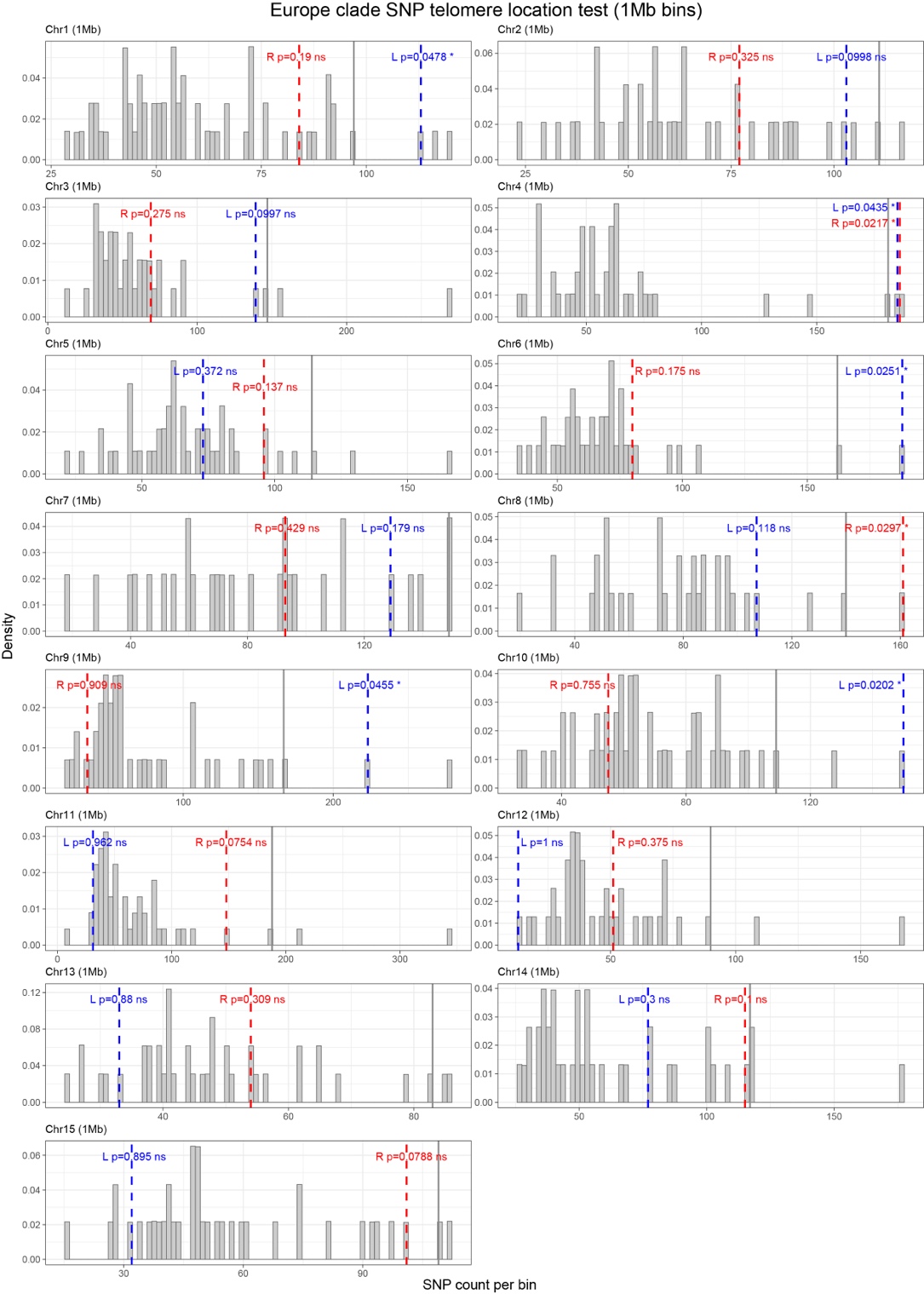

**Supplementary Figure 20. Location-based permutation test of telomeric enrichment of selected SNPs in the Europe clade (1-Mb bins).** Each panel corresponds to one chromosome (Chr1–Chr15), partitioned into non-overlapping 1-Mb bins. Grey bars represent the null distributions of selected SNP counts per 1-Mb bins on each chromosome, generated by randomly sampling bins from the same chromosome (including telomeric bins) 10^6^ times. The y-axis shows probability density and the x-axis shows the number of selected SNPs per bin. For each chromosome, the left telomere (blue dashed line) and right telomere (red dashed line) indicate the observed number of selected SNPs in the terminal 1-Mb bin at each chromosome end. The grey vertical line marks the 95th percentile of the null distribution. *P*-values next to each dashed line give the right-tail permutation probability that a randomly positioned 1-Mb bin on that chromosome would contain at least as many selected SNPs as the observed telomeric bin. ns = not significant and * = *P* < 0.05. Full statistical results are reported in Supplementary Table S6.

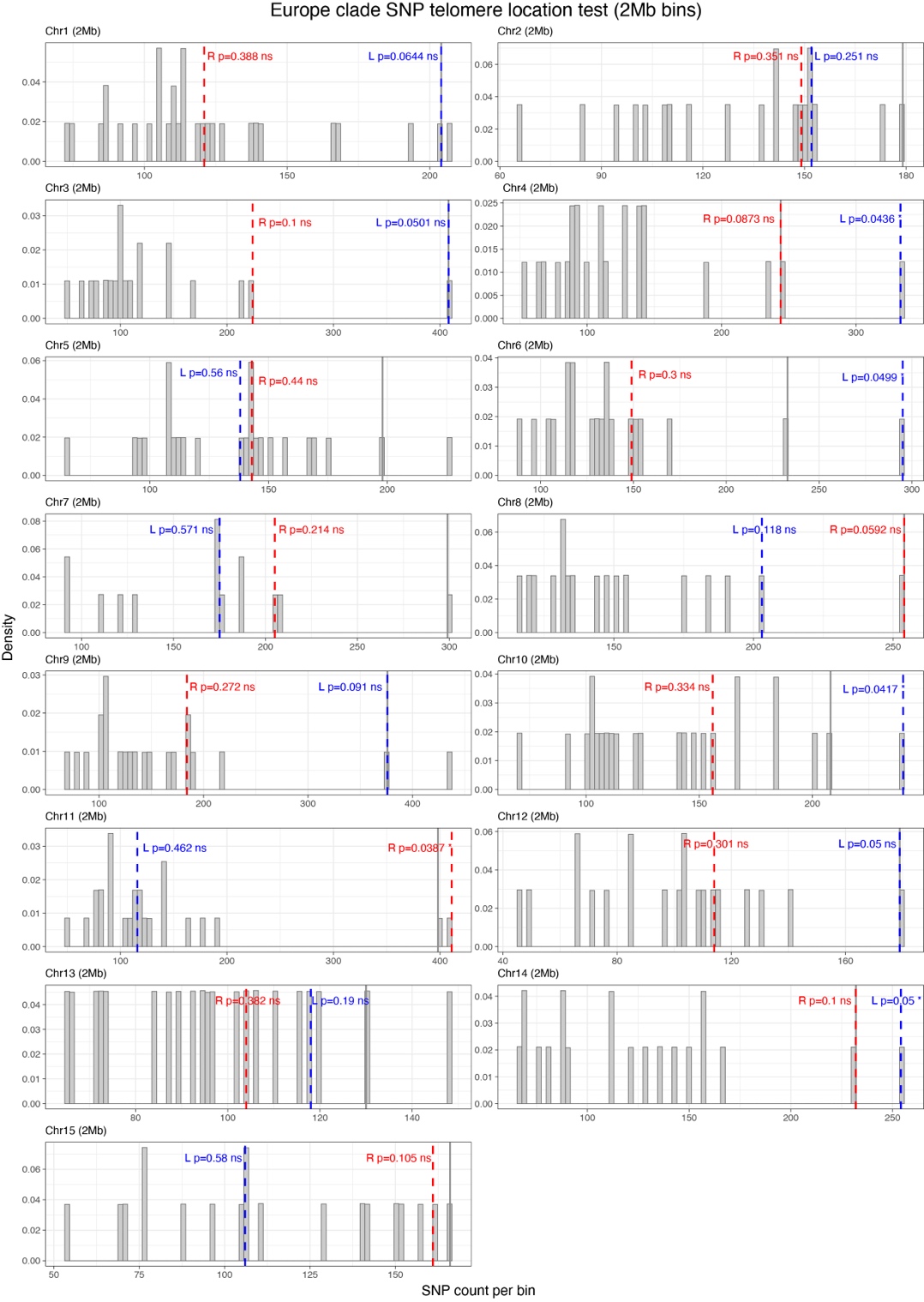

**Supplementary Figure 21. Location-based permutation test of telomeric enrichment of selected SNPs in the Europe clade (2-Mb bins).** Each panel corresponds to one chromosome (Chr1–Chr15), partitioned into non-overlapping 2-Mb bins. Grey bars represent the null distributions of selected SNP counts per 2-Mb bins on each chromosome, generated by randomly sampling bins from the same chromosome (including telomeric bins) 10^6^ times. The y-axis shows probability density and the x-axis shows the number of selected SNPs per bin. For each chromosome, the left telomere (blue dashed line) and right telomere (red dashed line) indicate the observed number of selected SNPs in the terminal 2-Mb bin at each chromosome end. The grey vertical line marks the 95th percentile of the null distribution. *P*-values next to each dashed line give the right-tail permutation probability that a randomly positioned 2-Mb bin on that chromosome would contain at least as many selected SNPs as the observed telomeric bin. ns = not significant and * = *P* < 0.05. Full statistical results are reported in Supplementary Table S7.
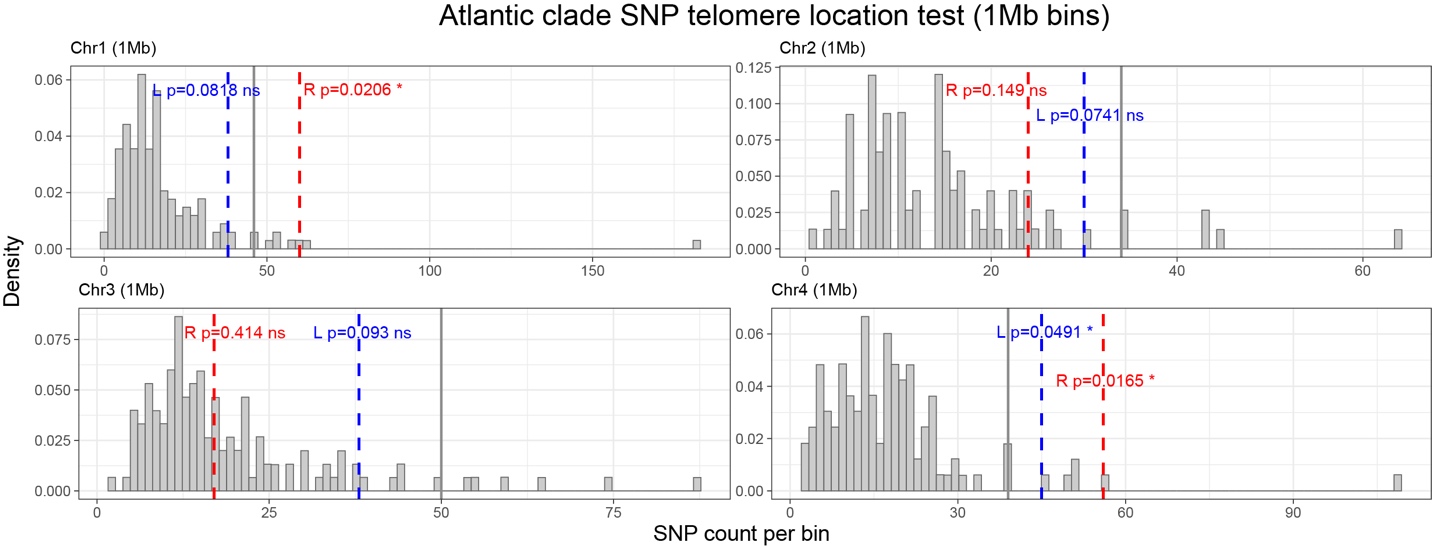

**Supplementary Figure 22. Location-based permutation test of telomeric enrichment of selected SNPs in the Atlantic clade (1-Mb bins).** Each panel corresponds to one chromosome (Chr1–Chr4), partitioned into non-overlapping 1-Mb bins. Grey bars represent the null distributions of selected SNP counts per 1-Mb bins on each chromosome, generated by randomly sampling bins from the same chromosome (including telomeric bins) 10^6^ times. The y-axis shows probability density and the x-axis shows the number of selected SNPs per bin. For each chromosome, the left telomere (blue dashed line) and right telomere (red dashed line) indicate the observed number of selected SNPs in the terminal 1-Mb bin at each chromosome end. The grey vertical line marks the 95th percentile of the null distribution. *P*-values next to each dashed line give the right-tail permutation probability that a randomly positioned 1-Mb bin on that chromosome would contain at least as many selected SNPs as the observed telomeric bin. ns = not significant and * = *P* < 0.05. Full statistical results are reported in Supplementary Table S6.

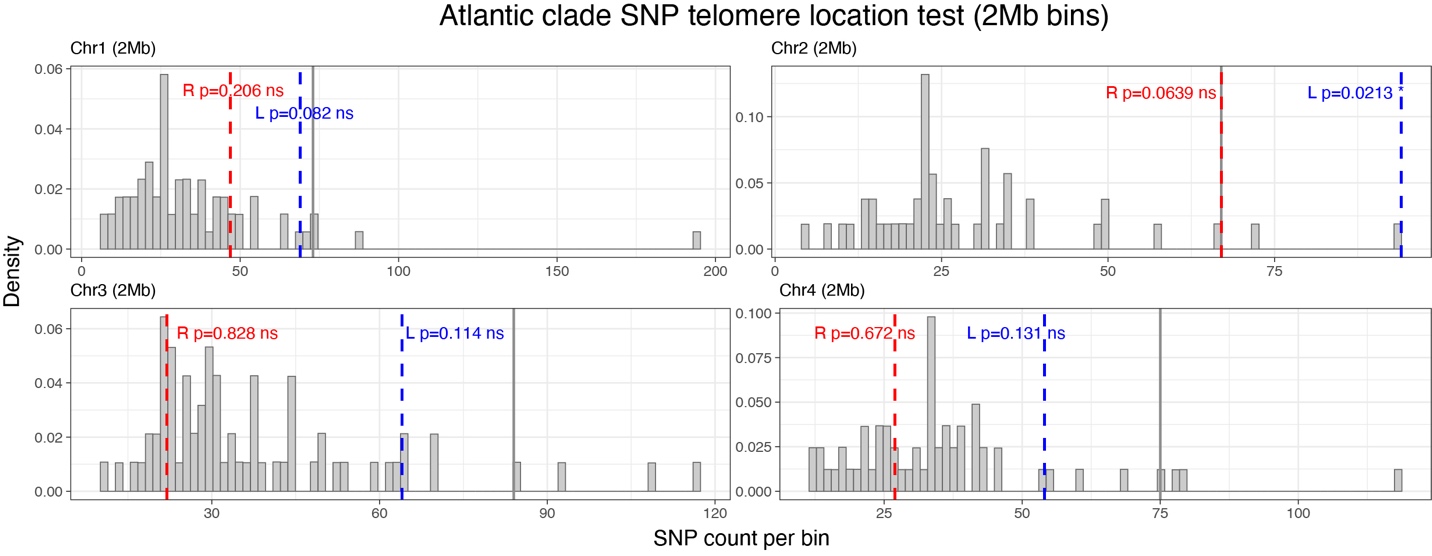

**Supplementary Figure 23. Location-based permutation test of telomeric enrichment of selected SNPs in the Atlantic clade (2-Mb bins).** Each panel corresponds to one chromosome (Chr1–Chr4), partitioned into non-overlapping 2-Mb bins. Grey bars represent the null distributions of selected SNP counts per 2-Mb bins on each chromosome, generated by randomly sampling bins from the same chromosome (including telomeric bins) 10^6^ times. The y-axis shows probability density and the x-axis shows the number of selected SNPs per bin. For each chromosome, the left telomere (blue dashed line) and right telomere (red dashed line) indicate the observed number of selected SNPs in the terminal 2-Mb bin at each chromosome end. The grey vertical line marks the 95th percentile of the null distribution. *P*-values next to each dashed line give the right-tail permutation probability that a randomly positioned 2-Mb bin on that chromosome would contain at least as many selected SNPs as the observed telomeric bin. ns = not significant and * = *P* < 0.05. Full statistical results are reported in Supplementary Table S7.

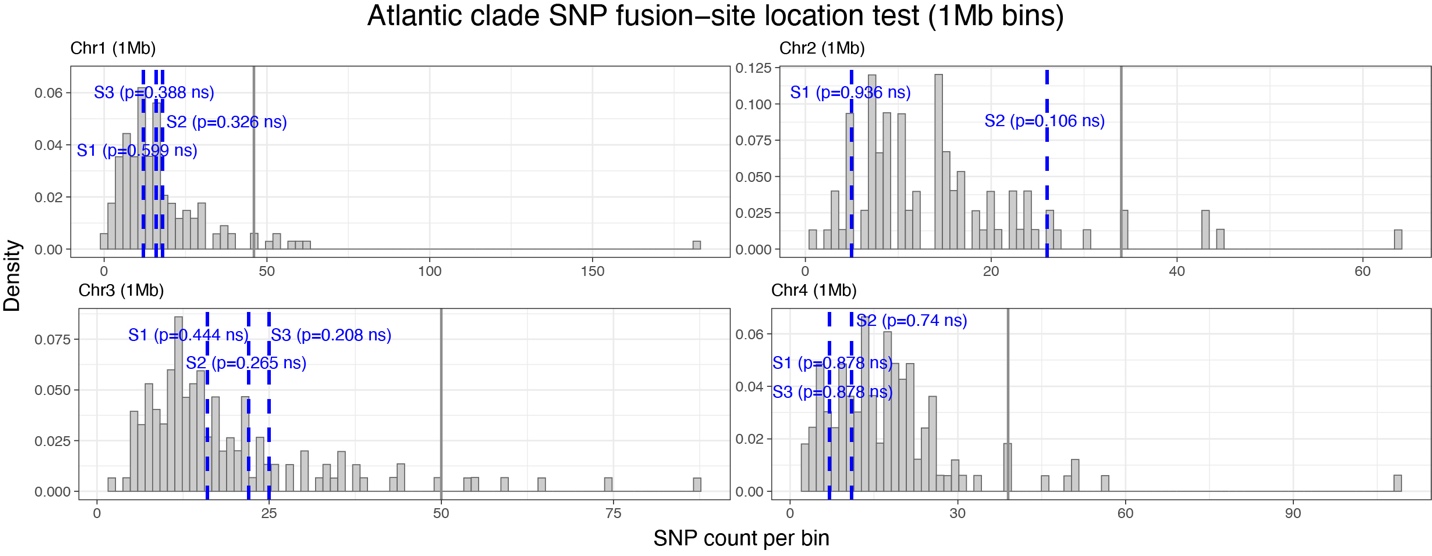

**Supplementary Figure 24. Location-based permutation test of fusion-site enrichment of selected SNPs in the Atlantic clade (1-Mb bins).** Each panel corresponds to one chromosome (Chr1–Chr4) of the Atlantic clade, partitioned into non-overlapping 1-Mb bins. Grey bars represent the null distributions of selected SNP counts per 1-Mb bins on each chromosome, generated by randomly sampling bins from the same chromosome (including telomeric bins) 10^6^ times. The y-axis shows probability density and the x-axis shows the number of selected SNPs per bin. For each chromosome, blue dashed vertical lines mark the observed number of selected SNPs in the 1-Mb bin containing the midpoint of each chromosomal fusion site (S1–S3). The grey vertical line indicates the 95th percentile of the null distribution. *P*-values shown next to each dashed line give the right-tail permutation probability that a randomly positioned 1-Mb bin on that chromosome would contain at least as many selected SNPs as the observed fusion-site bin. ns = not significant. Full statistical results are reported in Supplementary Table S8.

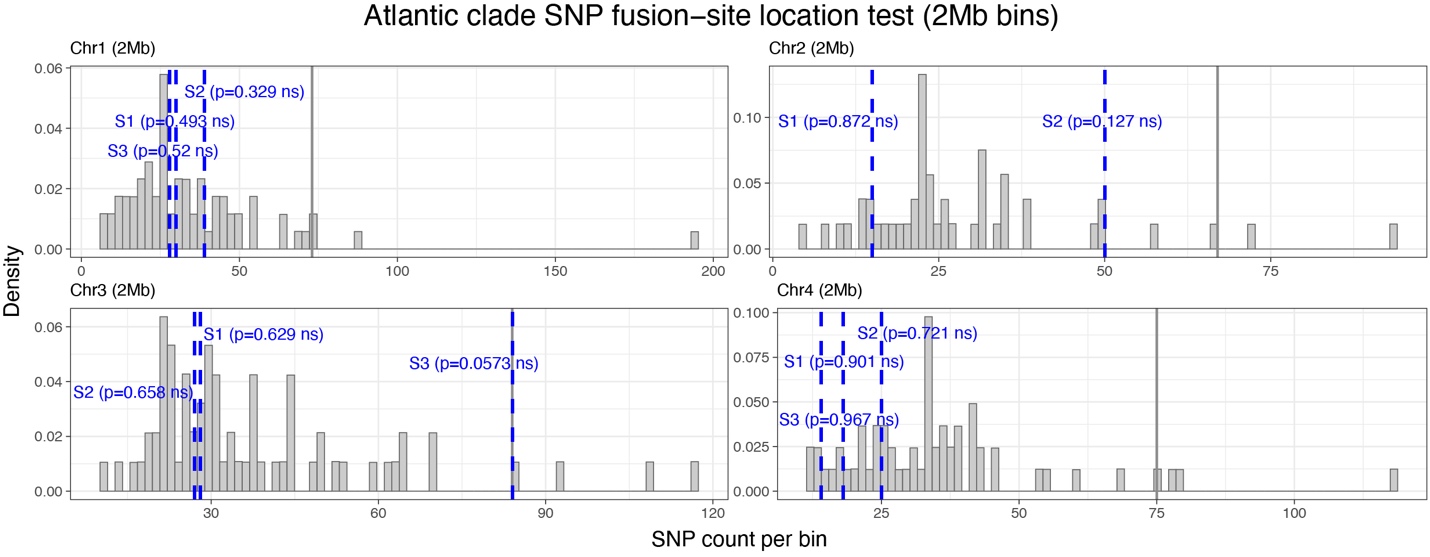

**Supplementary Figure 25. Location-based permutation test of fusion-site enrichment of selected SNPs in the Atlantic clade (2-Mb bins).** Each panel corresponds to one chromosome (Chr1–Chr4) of the Atlantic clade, partitioned into non-overlapping 2-Mb bins. Grey bars represent the null distributions of selected SNP counts per 2-Mb bins on each chromosome, generated by randomly sampling bins from the same chromosome (including telomeric bins) 10^6^ times. The y-axis shows probability density and the x-axis shows the number of selected SNPs per bin. For each chromosome, blue dashed vertical lines mark the observed number of selected SNPs in the 2-Mb bin containing the midpoint of each chromosomal fusion site (S1–S3). The grey vertical line indicates the 95th percentile of the null distribution. *P*-values shown next to each dashed line give the right-tail permutation probability that a randomly positioned 2-Mb bin on that chromosome would contain at least as many selected SNPs as the observed fusion-site bin. ns = not significant. Full statistical results are reported in Supplementary Table S8.

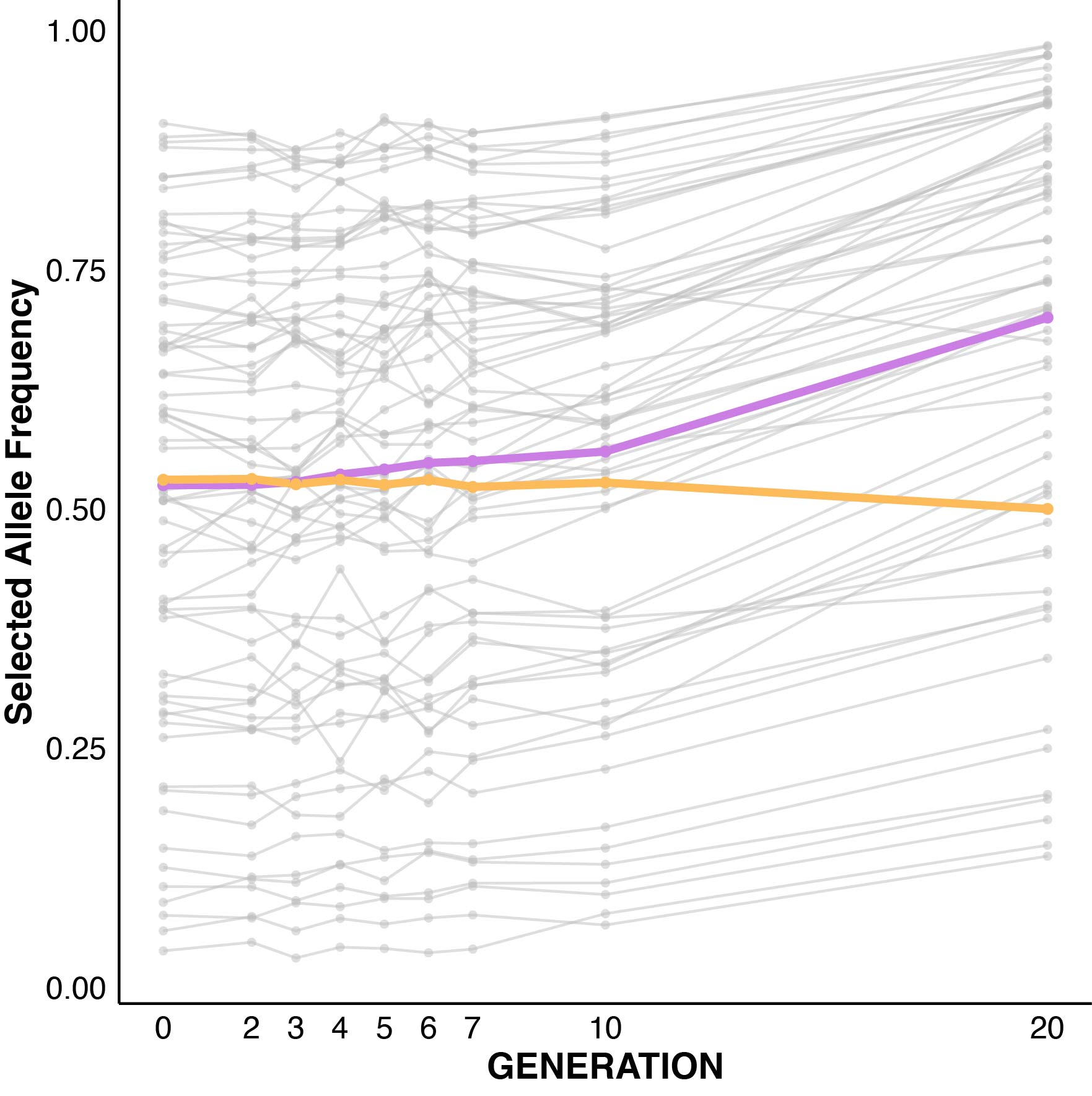

**Supplementary Figure 26. Time-resolved allele-frequency trajectories of selected haplotype blocks in the Atlantic clade across all sampled generations.** Gray lines show mean allele frequencies across replicate treatment lines for each selected allele. The purple line indicates the average frequency of all selected alleles in selection lines; the yellow line shows the corresponding average in control lines.

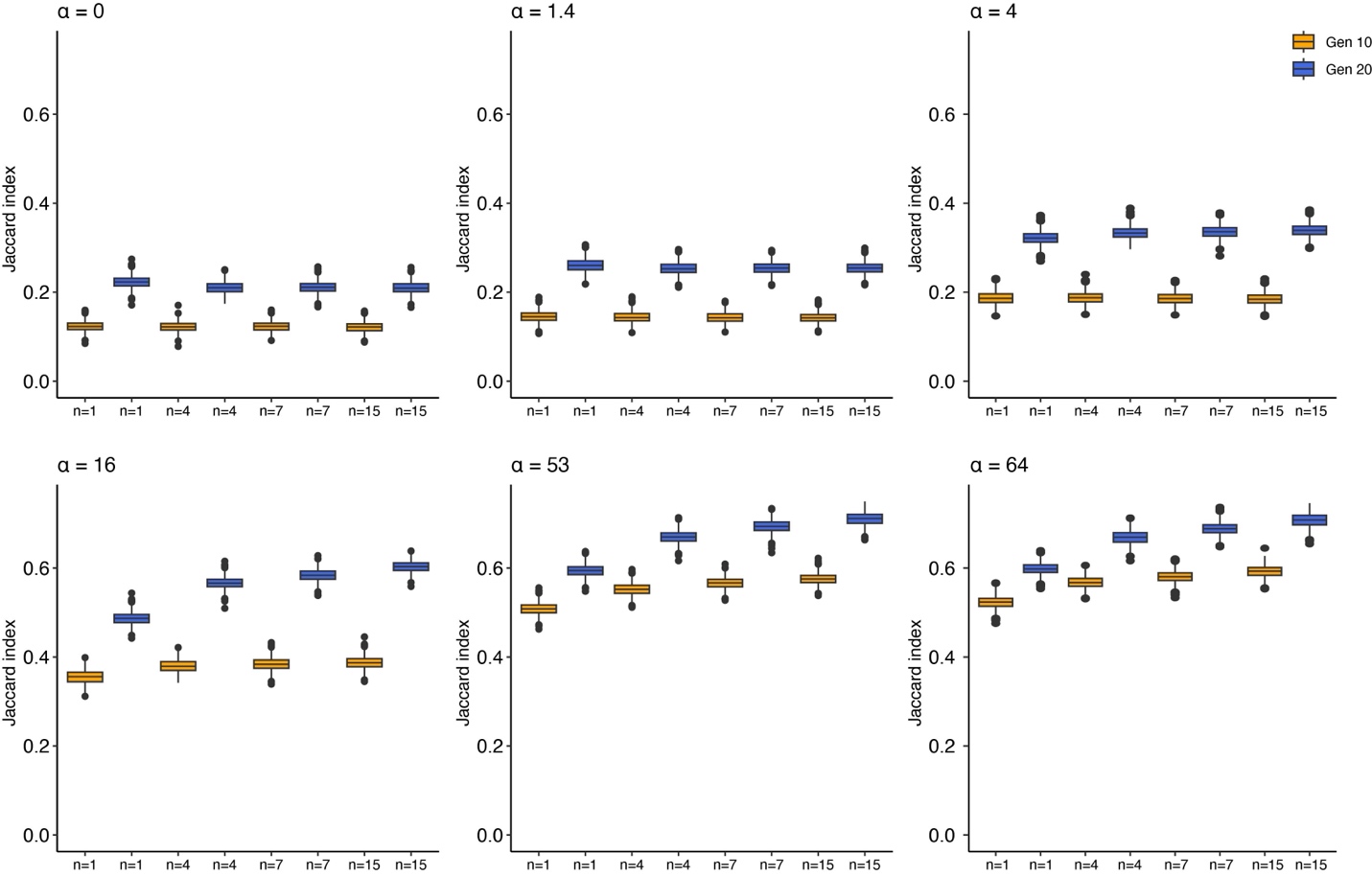

**Supplementary Figure 27. Effects of epistasis (α) and chromosome number (n) on the repeatability of adaptation (Jaccard index) at Generations 10 and 20.** Boxplots show the distribution of pairwise Jaccard indices across 1,000 simulation replicates for each combination of epistasis strength (α = 0, 1.4, 4, 16, 53, 64) and haploid chromosome number (n = 1, 4, 7, 15). Yellow boxes indicate Generation 10 and blue boxes indicate Generation 20. Under no epistasis (α = 0) or very weak epistasis (α = 1.4), Jaccard indices show only limited differences among recombining genome architectures, with a modest elevation in the fully linked genome (n = 1). Under moderate to strong epistasis (α ≥ 4), Jaccard indices generally increase with chromosome number, and this trend is strongest at high epistasis (α = 53 or 64), where 15 chromosome genomes consistently exhibit the highest Jaccard indices by Generation 20.
